# A neural circuit targeting technique for investigating functional input-output organization in the nervous system

**DOI:** 10.1101/2025.04.17.649280

**Authors:** Yusuke Kasuga, Xiaowei Gu, Tomoya Ohnuki, Akira Uematsu, Reiko Yoshida, Dai Yanagihara, Joshua P Johansen

## Abstract

Neurons communicate information across circuits and the function of cells in these circuits is determined by both the afferent inputs they receive and the efferent outputs they send to other brain regions^1,2^. To study the activity and function of specific neuronal populations, transneuronal anterograde^3^ and retrograde^4–6^ viral approaches have been employed to define neural circuit elements by inputs or outputs, respectively. However, what is missing is a way to study the function of neurons based on both their inputs and outputs. Applying a combination of multiple recombinases and transneuronal anterograde/retrograde viruses, we developed a technique called input-output <u>P</u>rojection-based <u>IN</u>tersectional <u>C</u>ircuit-tagging <u>E</u>nabled by <u>R</u>ecombinases (PINCER) to target specific neuronal cell types and investigate functional input-output organization in neural circuits. We show the logic and application of this technique with *in vivo* calcium imaging and optogenetic approaches to reveal the distinct functions and neural dynamics of connectivity defined neuronal populations in the amygdala for emotional processing. Specifically, PINCER allowed the parsing of valence and salience functions of the amygdala to reveal an input-output cell type selectively mediating aversive memory formation. This technique allows neuroscientists to identify novel subclasses of cells based on their combinatorial input-output anatomical connectivity, providing a tool for fine dissection of the functional properties of neural circuits.

## Introduction

In the neural circuits of the central nervous system, form gives rise to function. Brain circuits across species have been shaped by evolution to control many purposes from simple sensory processing and motor reflexes to complex emotional and cognitive experiences. Understanding the functional properties and organization of neural circuits can reveal the fundamental capabilities and constraints of the nervous system. The functionality of brain circuits arises from the combinatorial pattern of connections based on the inputs and outputs of individual neurons across brain regions^1,2^. This neural architecture serves as the foundational infrastructure for the processing and transmission of information within the nervous system (**Extended Data Fig. 1a**). Understanding how the function of cell types in brain circuits is defined by their input-output connectivity is thus a central goal of neuroscience research.

One approach for studying how anatomical connectivity underlies function relies on labelling neurons based on their axonal efferent projections to other brain regions. For example, retrograde viruses^4–6^ expressing recombinases have been used extensively for this purpose^7^. These virus-based techniques have empowered neuroscientists to probe the dynamics of neural circuit elements defined by their efferent projections to other brain regions (**Extended Data Fig. 1b**).

An alternative approach involves labeling cell types based on afferent inputs they receive from other brain regions. Although toxic to neurons over time, early anterograde trans-neuronal/synaptic viruses, such as the modified H129 strain of herpes simplex virus (HSV1-H129ΔTK)^8^, modified vesicular stomatitis virus (VSV)^9^, and rabies virus^10^ (restricted to peripheral nervous system usage) were useful for anatomical studies. However, the recent identification of a low-toxicity AAV serotype for transneuronal anterograde transfection (AAV1)^3^, offered a new approach for investigating properties of neural circuit elements defined by their afferent inputs (**Extended Data Fig. 1c**).

However, because neurons are defined by both their inputs and outputs, there remains a critical methodological gap for studying neuronal function based on anatomical connectivity. The combinatorial integration of transneuronal anterograde and retrograde viral tracing techniques could address this deficiency and open the door to the delineation of neural elements defined by both the afferent inputs they receive and the efferent outputs they make to other brain regions (**Fig. 1a**).

**Fig. 1.**
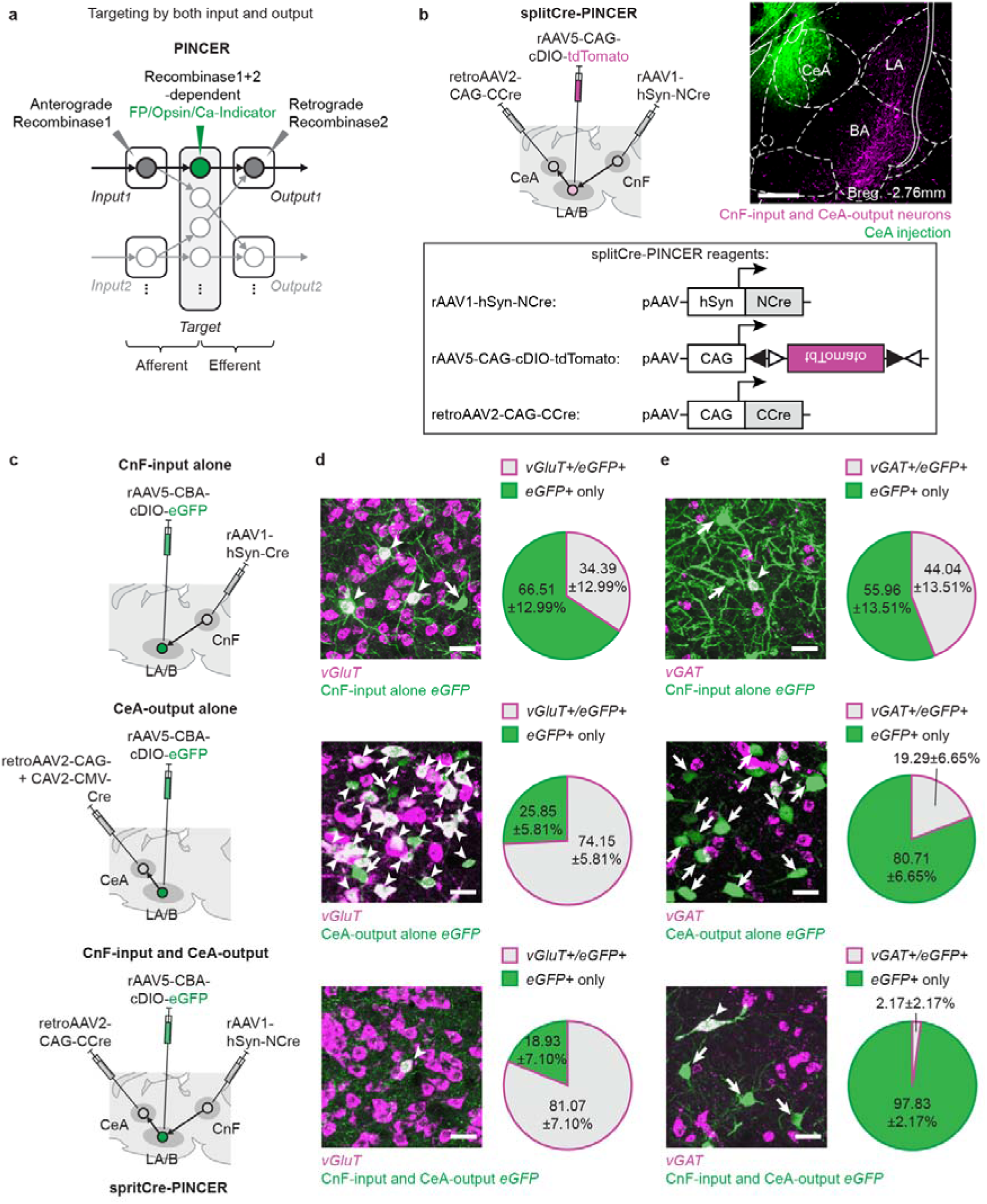
Logic and development of the PINCER method and molecular characterization of anatomical connectivity defined neuronal populations using PINCER. **a-b**, The concept, logic, and reagents for ‘PINCER’ method. **a**, Using transneuronal anterograde and retrograde viruses to express recombinases, PINCER captures a neural population based on both their afferent inputs and efferent outputs. FP: fluorescent protein. **b,** Schematic (*top, left*) showing the splitCre-PINCER approach in the lateral and basal nuclei of the amygdala (LA/B) for labeling cells based on the inputs they receive from the cuneiform nucleus (CnF) and the outputs they send to the central nucleus of the amygdala (CeA). Reagents (*bottom*) used and pAAV plasmid design for the splitCre-PINCER approach. Open and filled triangles, incompatible *loxP* sites. Representative image (*top, right*) showing successful implementation of PINCER to label the CnF-input and CeA-output neurons in LA/B (CnF-input/CeA-output neurons, rAAV5-cDIO-CAG-tdTomato+, magenta cells). Area of green labeled cells from CeA co-injection of rAAV5-CAG-eGFP+ without eGFP expression in LA/B demonstrates no leakage of retrograde virus from CeA into LA/B. **c-e**, Co-expression analysis of glutamatergic (*vGluT*) and GABAergic (*vGAT*) neuronal markers in the CnF-input alone, CeA-output alone and CnF-input/CeA-output neurons in LA/B. **c,** Schematic of viral combinations to label the CnF-input, CeA-output, and CnF-input/CeA-output neurons to co-label them with *vGluT* and *vGAT* mRNAs with the hybridization chain reaction fluorescence in situ hybridization (HCR-FISH) in LA/B. **d,** Representative images (*left*) of the CnF-input alone (*top*), CeA-output alone (*middle*), and CnF-input/CeA-output (*bottom*) neurons expressing *vGluT* in LA/B. Arrowheads and arrows denote anatomical connectivity defined neurons which are *vGluT+* or *vGlut-*, respectively. Pie charts (*right*) showing the percentage (mean±SEM) of the CnF-input alone neurons (*top*, N = 4 animals, total of n = 58 CnF-input neurons analyzed), CeA-output alone neurons (middle, N = 3 animals, total of n = 552 CeA-output neurons analyzed), or CnF-input/CeA-output neurons (bottom, N = 4 animals, total of n = 60 CnF-input/CeA-output neurons analyzed) co-labelled with *vGluT* in LA/B. **e,** Representative images (*left*) of the CnF-input alone (*top*), CeA-output alone (*middle*) and CnF-input/CeA-output (*bottom*) labeled neurons and neurons expressing *vGAT* in LA/B. Arrowheads and arrows denote anatomical connectivity defined neurons which are *vGAT+* or *vGAT-*, respectively. Pie charts (*right*) showing the percentage (mean±SEM) of the CnF-input alone neurons (*top*, N = 4 animals, total of n = 58 CnF-input neurons analyzed), CeA-output alone neurons (*middle*, N = 3 animals, total of n = 552 CeA-output neurons analyzed) or CnF-input/CeA-output neurons (*bottom*, N = 4 animals, total of n = 60 CnF-input/CeA-output neurons analyzed) co-labelled with *vGAT* in LA/B. Scale bars, **b,** 500μm, **d,e,** 30μm. **b,** Bregma distance in anterior-posterior direction.

With the emergence of viral based optogenetic tools, great progress in understanding the neural circuits of emotion has been achieved. This is particularly true for studies of the lateral and basal nuclei of the amygdala (LA/B), a crucial brain region for integrating sensory, salience and valence (aversive and rewarding) information^11–21^ to produce behavioral repertoires representing defensive and appetitive forms of emotional learning and memory^22–26^. Prominent theories suggest that emotions can be defined in a 2-dimensional framework superimposed along salience and valence axes^27–29^. While LA/B neurons encode these features, whether there are identified cell types that serve unique aversive, rewarding or salience functions is unclear. In fact, some results suggest that valence is encoded in population activity and that reward and aversive coding is not represented in separate groups of cells^30,31^. One possibility is that different functional populations of cells are defined by their anatomical connections with other brain regions. Some evidence suggests that specific subpopulations of valence specific LA/B neurons can be defined by their projection targets, but these cell types were typically found to encode and serve multiple behavioral learning roles^32–38^. Other work suggests that valence specific cells can be defined by a combination of molecular identity and projection target, but didn’t consider salience^38^. Given the importance of LA/B neurons in valence and salience processing as well as producing different behaviors (freezing, flight, avoidance, foraging etc), an intriguing hypothesis is that distinct functions of specific cell populations can be identified by their combinatorial input and output connectivity.

LA/B sends extensive efferent outputs to the central nucleus of the amygdala (CeA)^18,35,36,39–41^ and this pathway and CeA itself participates in aversive^35,36,39,40,42–44^, appetitive^34–36,41^ and salience^45–50^ processing. Moreover, recent evidence has revealed a monosynaptic brainstem pathway from the cuneiform nucleus (CnF) which conveys aversive information to LA/B to instruct aversive memory formation^51^. Although LA/B projections to CeA participate in multiple functions, it is possible that CeA projecting cells which receive aversive information from CnF may serve more specific aversive learning related functions. This circuit would thus be ideal for the development and testing of a technique for studying functional populations of cells based on their afferent inputs and efferent outputs.

Here, we introduce a novel technique, termed input-output <u>P</u>rojection-based <u>IN</u>tersectional <u>C</u>ircuit-tagging <u>E</u>nabled by <u>R</u>ecombinases (PINCER), to target a specific LA/B cell population that receives input from CnF and projects to CeA. Using combinatorial transneuronal anterograde and retrograde viruses with multiple recombinase systems, PINCER allows precise anatomical labeling and expression of calcium indicators for neuronal activity readout and opsins for optogenetic manipulation. We used PINCER to explore whether LA/B cell population defined by both CnF-input and CeA-output exhibits unique neural properties compared to those defined by either CnF-input alone or CeA-output alone. Specifically, we characterized the anatomy, activity patterns and functions of LA/B cells defined by both CnF-input and CeA-output during aversive associative learning and salience-based tasks and contrasted these findings with LA/B cell populations defined solely by either receiving CnF-input or projecting to CeA. We found that both input-alone and output-alone labeled cell populations contribute to both salience, reward and aversive learning functions, but that input-output labeled neurons specifically encode and participate in aversive learning. Together, this shows the power of the PINCER technique in defining specific functional cell populations and reveals principles of input-output organization in emotion processing circuits.

## Results

### Establishment of the PINCER input-output cell type specific tagging technique

We first examined the existence of a direct dual-transneuronal circuit from CnF through LA/B to CeA using viral based anatomical tracing methods. To this end, we injected the transneuronal anterograde AAV serotype-1^3^ to express Cre under control of the hSyn promoter (rAAV1-hSyn-Cre) into CnF, non-transneuronal AAV serotype-5 into LA/B to express double-floxed inverted orientation (DIO) tdTomato in a Cre-dependent manner (rAAV5-cDIO-tdTomato), and a viral cocktail of retrograde AAV (retroAAV) and CAV to express eGFP (rAAV2retro-CAG-eGFP and CAV2-CMV-eGFP) into CeA (**Extended Data Fig. 1d**). We previously showed that the projection from CnF to LA/B is unidirectional^51^. Using this combination of viral injections, neurons co-expressing the CnF-input (tdTomato) and CeA-output (eGFP) labels were identified (**Extended Data Fig. 1d**), supporting the existence of a direct CnF→LA/B→CeA circuit.

We then investigated how anterograde transneuronal delivery of Cre from CnF (rAAV1-hSyn-Cre) and retrograde delivery of Cre-dependent fluorophore from CeA (retroAAV2-CBA-cDIO-mCherry) labels cells in LA/B and CeA. This combination of viral injections labelled neurons in both LA/B and CeA (**Extended Data Fig. 1e**). The labeling in CeA is due to connectivity between CnF and CeA^51^. This result indicates that this combination of viral injections is not suitable for specifically targeting LA/B because infection of adjacent, or even distal cell populations could affect LA/B specificity for neuronal manipulations (e.g. opto/chemogenetics) and neural activity readout approaches. This observation motivated us to develop a strategy to target only LA/B neurons with CnF-input/CeA-output connectivity dependence.

To establish the PINCER technique, we used a combinatorial strategy combining both anterograde and retrograde transneuronal viruses to express recombinases and tag a neural population based on both their afferent inputs and efferent outputs (**Fig. 1a**). Two recombinase-based strategies were used in designing PINCER, one using the split-Cre recombinase (**Fig. 1b**, splitCre-PINCER) while the other took advantage of a combination of Cre and Flp recombinases (**Extended Data Fig. 2a**, Cre/Flp-PINCER). The split Cre recombinase system expressed the two halves of the split Cre (NCre and CCre, for N-terminal and C-terminal ends of halved Cre, respectively) in either anterograde or retrograde AAVs. When both NCre and CCre are expressed in the same cell, NCre and CCre fuse to become the functional Cre recombinase that can act on Cre-dependent gene encoding plasmids (**Fig. 1b**). Applying the splitCre-PINCER method, the transneuronal anterograde AAV1 injected into CnF to express NCre under hSyn promoter (rAAV1-hSyn-NCre) in LA/B neurons, retroAAV was injected into CeA to express CCre under CAG promoter (rAAV2retro-CAG-CCre) and the non-transneuronal AAV5 was injected into LA/B to express Cre-dependent tdTomato in Cre-dependent manner (rAAV5-cDIO-tdTomato) (**Fig. 1b**). In the dual recombinase approach Cre/Flp-PINCER was achieved by injecting the transneuronal anterograde rAAV1-hSyn-Cre into CnF, retrogradeAAV and CAV was injected into CeA to express a Cre-dependent Flp (rAAV2retro-CBA-cDIO-Flp and CAV2-CMV-cDIO-Flp) and the non-transneuronal AAV5 was injected into LA/B to express mCherry in Flp-dependent manner (rAAV5-CBA-fDIO-mCherry) (**Extended Data Fig. 2a,b**). This would allow for expression of mCherry only in cells that expressed both Cre (from CnF) and Flp (from CeA). These PINCER methods efficiently labeled neurons in LA/B which receive CnF-input and send CeA-output (CnF-input/CeA-output neurons; **Fig. 1b**). Reversing the viruses which express NCre and CCre also effectively labeled CnF-input/CeA-output neurons (**Extended Data Fig. 1f,g,h**). Moreover, negative control experiments in which different anterograde or retrograde viruses were omitted did not find input-output labeled neurons (**Extended Data Fig. 3**).

Cre is known for its high efficiency recombinase action compared to Flp^52^. Therefore, we compared the expression time-course of splitCre-PINCER and Cre/Flp-PINCER methods for labeling CnF-input/CeA-output neurons in LA/B. This experiment revealed that the splitCre-PINCER method achieved maximum labeling of CnF-input/CeA-output neurons with shorter infection duration (4 weeks) compared to the Cre/Flp-PINCER method (8 weeks) (**Extended Data Fig. 2c,d**). Both methods label similar numbers of the CnF-input/CeA-output neurons in LA/B if the infection duration was adequate (splitCre-PINCER >4 weeks, Cre/Flp-PINCER >8 weeks infection). This result shows that the splitCre-PINCER approach has higher efficiency resulting in faster labeling of neurons compared with the Cre/Flp-PINCER method.

Taken together, these results demonstrate that PINCER methods can target a specific neuronal population in LA/B which receive CnF input and send efferent projections to CeA. There are different combinations of recombinases that can be used for PINCER, but utilizing the split-Cre system in the PINCER approach (splitCre-PINCER) is more efficient in driving transgene expression in input-output neurons.

### Anatomical characterization of input-output circuit organization in LA/B using PINCER

Having established the PINCER approach, we next used it to characterize the organization and molecular makeup of anatomical connectivity defined cell populations in LA/B. We first quantified the proportion of CnF-input labeled cells which also project to CeA. Labeling of eGFP+ CnF-input alone neurons was achieved using injections of the transneuronal anterograde rAAV1-hSyn-Cre in CnF and rAAV5-CBA-cDIO-eGFP in LA/B. In combination, mCherry+ CnF-input/CeA-output neurons were labeled using retrograde viruses expressing Cre-dependent Flp recombinase injected into CeA and AAV5 expressing Flp dependent mCherry in LA/B (Cre/Flp-PINCER). Quantifying the percentage of eGFP+ CnF-input alone labeled and mCherry+ CnF-input/CeA-output labeled cells revealed that 49.11% of CnF-input alone neurons are CnF-input/CeA-output neurons (**Extended Data Fig. 4a**). Next, we examined the proportion of CeA-output alone labeled cells which also receive input from CnF. Combinatorial labeling of eYFP+ CnF-input/CeA-output neurons under the control of Cre and Flp, with the Cre/Flp-PINCER method and co-labeling of tdTomato+ CeA-output alone neurons, with an injection of transneuronal retrograde rAAV2retro-CAG-tdTomato in CeA, in LA/B revealed that 5.10% of CeA-output alone neurons are CnF-input/CeA-output neurons in LA/B (**Extended Data Fig. 4b**). Finally, to determine the topographic distribution of this input-output labeled cell population, we performed serial imaging of CnF-input/CeA-output cells throughout LA/B and found that these neurons are preferably distributed between the mid to posterior aspect of LA/B (**Extended Data Fig. 4c,d,e**).

We next used PINCER to identify the molecular-neurotransmitter identity (glutamatergic excitatory vs. GABAergic inhibitory neuronal types) of the input alone, output alone and input-output defined LA/B neurons. eGFP+ CnF-input alone neurons were labeled through injections of transneuronal anterograde rAAV1-hSyn-Cre in CnF and rAAV5-CBA-cDIO-eGFP in LA/B; eGFP+ CeA-output alone neurons were labeled with injections of rAAV5-CBA-cDIO-eGFP in LA/B and a cocktail of transneuronal retrograde retroAAV2-CAG-Cre and CAV2-CMV-Cre in CeA; and eGFP+ CnF-input/CeA-output neurons were labeled with the splitCre-PINCER approach (**Fig. 1c**). These three eGFP+ anatomical connectivity defined cell classes were stained with glutamatergic neuronal marker *vesicular glutamate transporters* (*vGluT*) and GABAergic neuronal marker *vesicular GABA transporters* (*vGAT*) mRNAs using hybridization chain reaction fluorescence in situ hybridization (HCR-FISH). 34.39% and 44.04% of the CnF-input alone neurons in LA/B were *vGluT*+ and *vGAT*+, respectively. 74.15% and 19.29% of the CeA-output alone neurons in LA/B were *vGluT*+ and *vGAT*+ whereas the vast majority (81.07%) and a small fraction (2.17%) of the CnF-input/CeA-output neurons in LA/B were *vGluT*+ and *vGAT*+, respectively (**Fig. 1d,e**). These results indicate that CnF-input alone and CeA-output alone neurons are a mixture of glutamatergic and GABAergic neurons, whereas the CnF-input/CeA-output neurons are mostly glutamatergic. We then characterized the three anatomical connectivity defined LA/B cell types based on their expression of markers for GABAergic interneuron subclasses based on expression of *somatostatin* (*SST*), *parvalbumin* (*PV*) and *vasoactive intestinal peptide* (*VIP*) mRNAs using HCR-FISH. 23.16%, 11.97% and 9.28% of the CnF-input neurons in LA/B were *SST*+, *PV*+ and *VIP*+, respectively. 2.56%, 1.11% and 1.70% of the CeA-output alone neurons were *SST*+, *PV*+, and *VIP*+, respectively. A small proportion (2.84%) of the CnF-input/CeA-output neurons were *SST*+, but none (0.00%) of these neurons were *PV*+ and *VIP*+, respectively (**Extended Data Fig. 5a,b,c**).

To further extend the PINCER technique, we combined it with cell type specific retrograde transsynaptic rabies virus tracing for identifying which molecularly defined neuronal subtypes in CeA the CnF-input/CeA-output neurons in LA/B project to. We selected CeA SST+ cells as a target because they are known to control defensive responding following aversive learning^53,54^. For this purpose, we injected a combination of viruses into an SST-Cre transgenic rat including transneuronal anterograde rAAV1-hSyn-Flp into CnF, rAAV5-CBA-fDIO-mCherry into LA/B, and the rabies glycoprotein (G) and avian TVA-receptor helper viruses (rAAV9-CAG-cDIO-HA-2A-G and rAAV9-CAG-cDIO-TVA) followed by an injection of EnvA-ΔG-Rabies-eGFP into CeA (**Extended Data Fig. 5d**). This approach allowed for colocalization of the HA-tagged (for G labeling) and rabies eGFP+ SST+ neurons in CeA as starter neurons for rabies transfection (**Extended Data Fig. 5e**). The identification of mCherry+ CnF-input alone neurons in LA/B co-expressing the rabies eGFP with this approach shows that some of the CnF-input/CeA-output neurons in LA/B provide monosynaptic inputs onto SST+ inhibitory interneurons in CeA (**Extended Data Fig. 5f**).

Taken together, these results demonstrate the wide range of PINCER applications for delineating input-output circuit architecture, identifying molecularly defined cell type identities of input-output projection-defined neurons in comparison to input-alone or output-alone cells, and for elucidating which molecularly defined cell types in the output region the input-output projection-defined neurons project to.

### PINCER reveals differential coding of salience and aversive learning in distinct connectivity defined cell types

We next used the PINCER technique to perform cell-type specific calcium imaging^55–57^ and record neural activity in the input-output projection-defined LA/B cell population and compared this to the dynamics of the input or output alone populations during aversive learning and memory tasks in which the amygdala participates^23,24^.

We first expressed the calcium indicator GCaMP6s in the CnF-input defined neurons by injecting the transneuronal anterograde rAAV1-hSyn-Cre into CnF and rAAV5-Syn-cDIO-GCaMP6s injected in LA/B, followed by implantation of an optical fiber in LA/B (**Fig. 2b**). Animals then underwent aversive conditioning while GCaMP signals were recorded (**Fig. 2a**). They exhibited no freezing behavior to CS presentation before aversive conditioning (CS pre-conditioning) but began freezing in response to the CS after it was paired with the shock US (conditioning) and exhibited CS-evoked freezing behavior when it was presented at the memory ‘retrieval’ timepoint following conditioning (**Extended Data Fig. 6**). The CnF-input alone neurons in LA/B responded to the CS before and after conditioning (**Fig. 2c**) and displayed a significant increase in CS-evoked responding after compared to before aversive conditioning (**Fig. 2d**). During aversive training, CS responses did not change significantly but US responses were reduced from early to late conditioning (**Extended Data Fig. 7a,b,c,d**). These results indicate that the CnF-input alone LA/B neurons respond to auditory CSs both before and after aversive conditioning but exhibit a learning induced enhancement, responding more strongly after training.

**Fig. 2.**
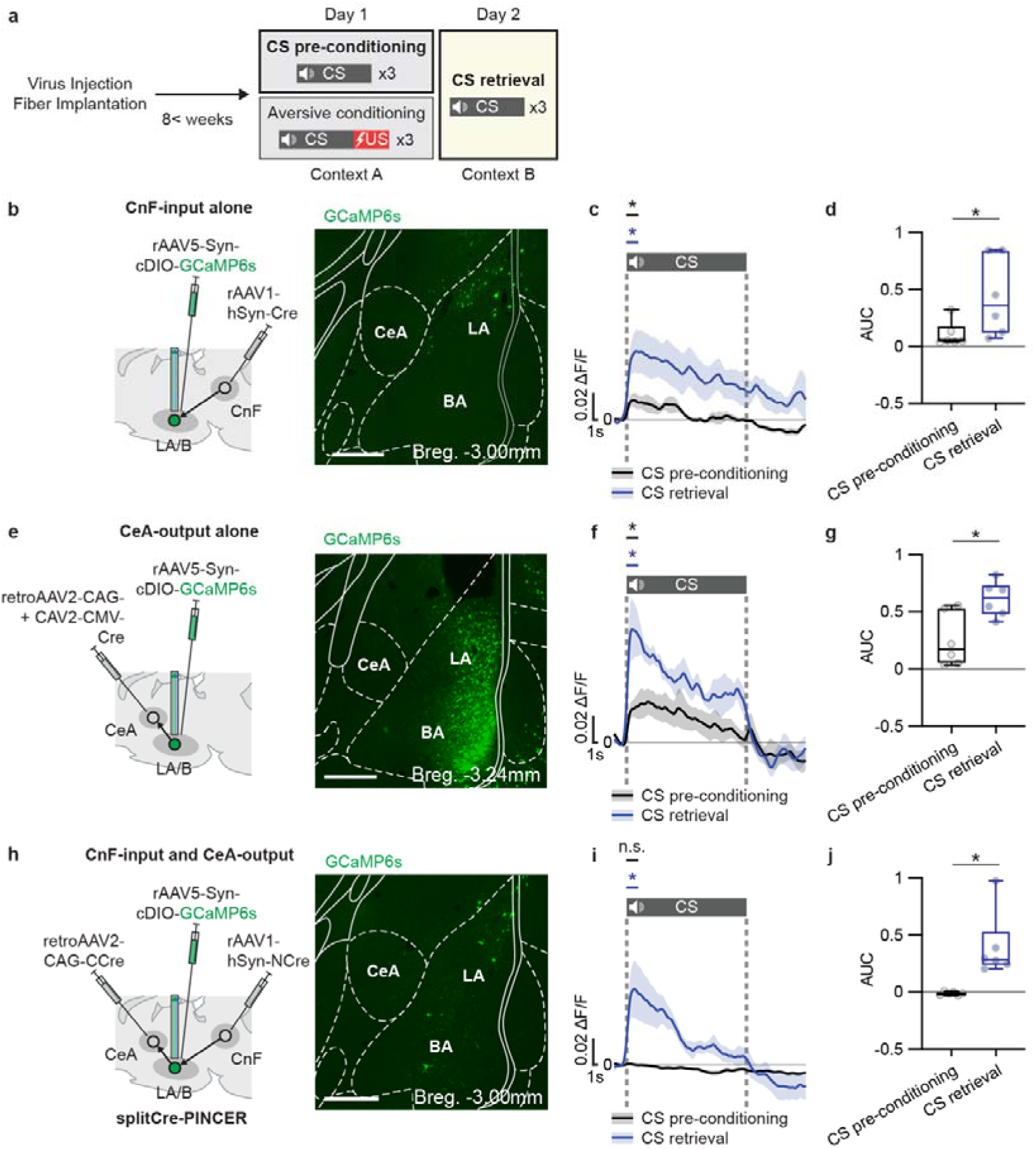
Calcium imaging during aversive learning in anatomical connectivity defined cell types using PINCER. **a,** Schematic of experimental design. **b-d**, Increased CS-evoked calcium responses of the CnF-input neurons in LA/B during pre-conditioning vs. retrieval. **b,** Schematic (*left*) of viral combinations to express GCaMP6s in the CnF-input alone neurons in LA/B for fiber photometry. Representative image (*right*) of GCaMP6s expression in the CnF-input alone neurons. **c,** Peri-event time histogram showing group averaged calcium responses to CS in pre-conditioning (black) and retrieval (blue) (N = 6 animals). CS-evoked calcium responses were both significantly higher than baselines period (2 seconds before and after CS onset, pre-conditioning *P* = 0.0312 and retrieval *P* = 0.0312, Wilcoxon test). **d,** Bar graphs show significantly increased CS-evoked ΔF/F area under curve (AUC) responses at retrieval (blue) vs. pre-conditioning (black) (*P* = 0.0312, Wilcoxon test). **e-g**, Increased CS-evoked calcium response of the CeA-output alone neurons in LA/B during retrieval vs. pre-conditioning. **e,** Schematic (*left*) of viral combinations to express GCaMP6s in the CeA-output alone neurons. Representative image (*right*) of GCaMP6s expression in the CeA-output neurons. **f,** Peri-event time histogram showing group averaged calcium responses to CS in pre-conditioning (black) compared with retrieval (blue) (N = 6 animals). CS-evoked calcium responses were significantly higher than baseline period (pre-conditioning *P* = 0.0312 and retrieval *P* = 0.0312, Wilcoxon test). **g,** Bar graphs show significantly increased CS-evoked ΔF/F AUC responses at retrieval (blue) vs. pre-conditioning (black) (*P* = 0.0312, Wilcoxon test). **h-j**, Increased CS-evoked calcium responses of the CnF-input/CeA-output neurons in LA/B during pre-conditioning vs. retrieval. **h,** Schematic (*left*) of viral combinations to express GCaMP6s in the CnF-input/CeA-output neurons with splitCre-PINCER approach. Representative image (*right*) of GCaMP6s expression in the CnF-input/CeA-output neurons. **i,** Peri-event time histogram showing group averaged calcium responses to CS in pre-conditioning (black) and retrieval (blue) (N = 6 animals). CS-evoked responses in retrieval but not pre-conditioning were significantly higher than baseline periods (pre-conditioning *P* = 0.8348 and retrieval *P* = 0.0312, Wilcoxon test). **j,** Bar graphs show significantly increased CS-evoked ΔF/F AUC responses at retrieval (blue) vs. pre-conditioning (black) (*P* = 0.0312, Wilcoxon test). **b,e,h,** Scale bars: 500μm. **b,e,h,** Bregma distance in anterior-posterior direction. **c,f,i,** Shades: SEM. **d,g,j,** In box plot: center line, median; box limits, upper and lower quartiles; whiskers, two-sided 95^th^ percentile range of distribution. n.s.: *P* > 0.05, *: *P* < 0.05.

Next, we studied the aversive learning induced changes in neural processing in the CeA-output defined LA/B neurons. We expressed GCaMP6s in this cell population using retrograde virus (retroAAV2-CAG-Cre and CAV2-CMV-Cre) injection into CeA and rAAV5-Syn-cDIO-GCaMP6s injection into LA/B followed by optical fiber placement in LA/B (**Fig. 2e**). CeA-output defined LA/B neurons responded to auditory CSs before and after conditioning (**Fig. 2f**) and exhibited a significant increase in CS-evoked responding after compared to before aversive conditioning (**Fig. 2g**). During aversive conditioning, neither CS nor US evoked responses changed significantly (**Extended Data Fig. 7e,f,g,h**). These results indicate that auditory CS-evoked calcium responses of the CeA-output alone neurons in LA/B are evident both before and after aversive conditioning but that CS responses become potentiated at memory retrieval following aversive learning.

Finally, we examined neural processing in the CnF-input/CeA-output defined LA/B neurons during aversive learning and memory. GCaMP6s was expressed in the CnF-input/CeA-output LA/B cells through injections of rAAV5-Syn-cDIO-GCaMP6s in LA/B using the splitCre-PINCER approach described above followed by optical fiber placement in LA/B (**Fig. 2h**). Contrasting with the CnF-input alone and CeA-output alone neurons, the CnF-input/CeA-output neurons did not respond to the CS during the pre-conditioning session prior to aversive training. However, input-output defined neurons did respond to the CS at the memory retrieval timepoint after conditioning (**Fig. 2i**) and demonstrated a significantly larger CS-evoked response at memory retrieval compared with prior to aversive learning (**Fig. 2j**). During aversive conditioning, neither CS nor US evoked responses changed significantly (**Extended Data Fig. 7i,j,k,l**). Together with the previous results, these findings indicate that CnF-input/CeA-output LA/B neurons do not respond to auditory stimuli prior to aversive learning, unlike input and output alone defined neurons. However, all cell populations exhibit an aversive learning induced increase in CS-evoked responding during memory retrieval.

While LA/B is known to be important for aversive learning and memory, it also encodes the salience and novelty of sensory stimuli^14,16,18,19,21^. Indeed, salience and valence are two features which have been hypothesized to define distinct axes underlying emotional processing^29^. While all cell populations we recorded showed a learning induced enhancement of CS responding (**Fig. 3d,g,j)**, one of the unique coding features of the CnF-input/CeA-output neurons in LA/B was the absence of responding to sensory stimuli prior to aversive conditioning **(Fig. 3i)**. This contrasted with the robust pre-learning sensory stimulus evoked activity observed in the CnF-input and CeA-output alone neurons **(Fig. 3c,d,e,f)**. Based on this finding, we hypothesized that activity in the CnF-input and CeA-output alone defined neurons reflects both salience and aversive learning, while the input-output defined cells selectively encode aversive learning. A hallmark of salience coding is strong responding to initial presentations of sensory stimuli and diminution of the response over repeated presentations as attention wanes. The effects of this reduced salience of stimuli can be assayed behaviorally using the latent inhibition paradigm in which repeated CS presentations are thought to reduce the salience of CSs, which then impairs subsequent aversive associative learning^21,58^.

**Fig. 3.**
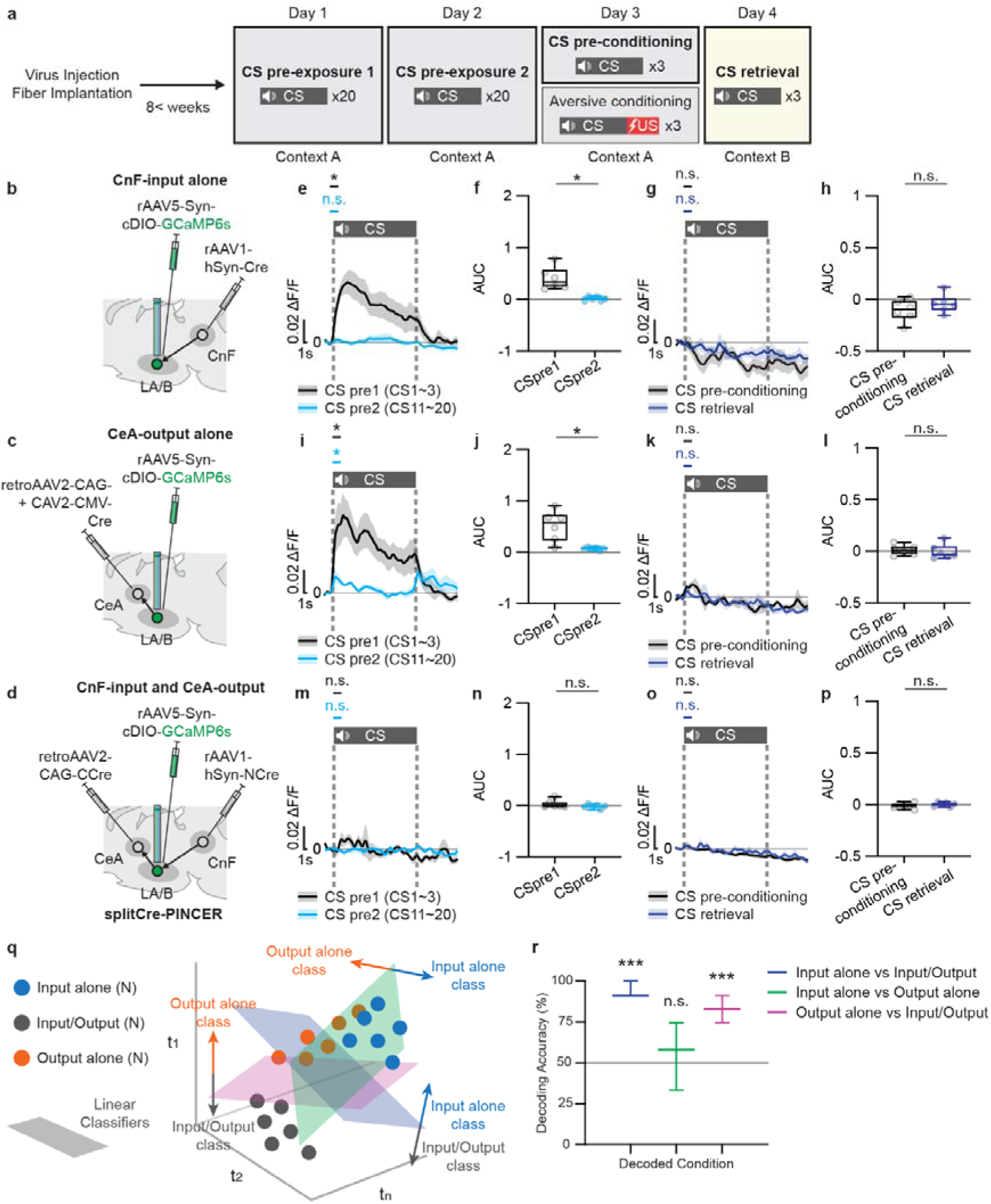
Distinct neural dynamics underlying salience processing in anatomical connectivity defined cell types revealed through PINCER. **a,** Schematic of latent inhibition experimental design**. b-d,** schematic showing viral combinations to express GCaMP6s in the **b,** CnF-input alone, **c,** CeA-output alone and **d**, CnF-input/CeA-output neurons in LA/B during latent inhibition. **b,e-h,** Decreased responses with repeated CS presentations and blockade of conditioning induced enhancement in CS processing in the CnF-input alone neurons during latent inhibition. **e,** Peri-event time histogram showing group averaged CS-evoked calcium responses in the first (black) compared with the last (cyan) trials of CS pre-exposure (N = 6 animals). CS responses to the first (black) but not the last (cyan) trials were significantly higher than baselines prior to CS onset (2 seconds before vs. after CS onset, CS pre-exposure first trials *P* = 0.0312 and last trials *P* = 0.5625, Wilcoxon test by corresponding baselines prior to CS onsets). **f,** Bar graphs show significantly decreased CS-evoked ΔF/F area under curve (AUC) in the last (cyan) vs. the first CS pre-exposure trials (black) (*P* = 0.0312, Wilcoxon test). **g,** Peri-event time histogram showing group averaged CS-evoked calcium responses during pre-conditioning (black) and retrieval (blue) exhibited no responses to CSs (CS pre-conditioning *P* > 0.9999 and retrieval *P* = 0.2188, Wilcoxon test). **h,** Bar graphs show no differences in CS-evoked ΔF/F AUCs comparing pre-conditioning (black) and retrieval (blue) (*P* = 0.5625, Wilcoxon test). **c,i-l**, Decreased responses with repeated CS presentations and blockade of conditioning induced enhancement in CS processing in the CeA-output alone neurons during latent inhibition. **i,** Peri-event time histogram showing group averaged CS-evoked calcium responses in the first (black) compared with the last (cyan) trials of CS pre-exposure (N = 6 animals). CS responses to the first (black) but not the last (cyan) trials were significantly higher than baselines prior to CS onset (CS pre-exposure first trials *P* = 0.0312 and last trials *P* = 0.0312, Wilcoxon test). **j,** Bar graphs show significantly decreased CS-evoked ΔF/F AUC in the last (cyan) vs. the first CS pre-exposure trials (black) (*P* = 0.0312, Wilcoxon test). **k,** Peri-event time histogram showing group averaged CS-evoked calcium responses during pre-conditioning (black) and retrieval (blue) exhibited no responses to CSs (CS pre-conditioning *P* = 0.0625 and retrieval *P* = 0.8348, Wilcoxon test). **l,** Bar graphs show no differences in CS-evoked ΔF/F AUCs comparing pre-conditioning (black) and retrieval (blue) (*P* > 0.9999, Wilcoxon test). **d,m-p,** No CS-responding during pre-exposure and blockade of conditioning induced enhancement in CS processing in the CnF-input/CeA-output neurons during latent inhibition. **m,** Peri-event time histogram showing group averaged CS-evoked calcium responses in the first (black) compared with the last (cyan) trials of CS pre-exposure (N = 6 animals). These neurons did not respond significantly to CS during pre-exposure compared with baseline activity levels (the first CS pre-exposure trials *P* > 0.9999 and the last CS pre-exposure trials *P* > 0.9999, Wilcoxon test). **n,** Bar graphs show no difference in CS-evoked ΔF/F AUCs in the last (cyan) vs. the first CS pre-exposure trials (black) (*P* = 0.8348, Wilcoxon test). **o,** Peri-event time histogram showing group averaged CS-evoked calcium responses during pre-conditioning (black) and retrieval (blue) exhibited no responses to CSs (pre-conditioning *P* > 0.9999 and retrieval *P* = 0.2488, Wilcoxon test). **p,** Bar graphs show no differences in CS-evoked ΔF/F AUCs comparing pre-conditioning (black) and retrieval (blue) (*P* = 0.5625, Wilcoxon). **q,** Schematic of the decoding method (see Figure S10 and Methods). Linear classifiers were trained to classify neural dynamics of the CnF-input/CeA-output from those of the CnF-input or CeA-output alone or CnF-input vs. CeA-output alone using activity patterns of these distinct neural populations during latent inhibition of aversive conditioning. **r,** Decoders could distinguish activity patterns of the CnF-input/CeA-output from the CnF-input or CeA-output alone neurons with high accuracy compared with chance level (input alone vs input-output, *P* = 0.0050, bootstrap test; output alone vs input-output, *P* = 0.0050, bootstrap test). However, decoders could not distinguish the activity patterns of the CnF-input from the CeA-output alone cell types at above chance level (input alone vs output alone, *P* = 0.4930, bootstrap test). n.s.: *P* > 0.05, *: *P* < 0.05, ***: *P* < 0.001. **e,g,i,k,m,o,** Shades: SEM. **f,h,j,l,n,p** In box plot: center line, median; box limits, upper and lower quartiles; whiskers, two-sided 95^th^ percentile range of distribution. **r,** Bars: median, and error bars: two-sided 95^th^ percentile range of distribution.

To test the idea that input or output alone defined LA/B neurons encode salience whereas input-output labeled neurons do not, we compared the calcium dynamics of the three anatomical connectivity defined populations in LA/B while animals engaged in the latent inhibition of aversive conditioning task. We hypothesized that neural populations encoding salience would respond robustly to initial presentations of sensory stimuli, but that responses would reduce with repeated CS presentations. Furthermore, we anticipated that cell populations encoding learning would not show the observed learning induced enhancement if the conditioning phase was preceded by repeated CS presentations. In the latent inhibition phase of the task, CSs were repeatedly presented to the animals prior to paired CS-US presentations followed by standard aversive learning and memory tests (**Fig. 3a**). Animals did not show freezing behavior to CSs during pre-exposure before conditioning (CS pre-exposure 1 and 2), developed mild freezing behavior to CSs during aversive conditioning but did not exhibit CS-evoked freezing behavior to CS presentation during the retrieval test after learning (**Extended Data Fig. 8**). This shows that latent inhibition impaired the formation of aversive memories.

Using the latent inhibition paradigm and transneuronal anterograde or retrograde viral techniques, GCaMP6s was expressed in the CnF-input or CeA output alone neurons in LA/B along with an optical fiber in LA/B to perform fiber photometry from these cell populations (**Fig. 3b,c and Extended Data Fig. 9a,b and Extended Data Fig. 10a,b**). During CS pre-exposure, the CnF-input and CeA-output alone neurons in LA/B responded initially to CSs, but their responses diminished over repeated CS presentations (**Fig. 3e,f,i,j and Extended Data Fig. 9d,e,f,g**). Moreover, these cell populations did not respond to CSs during aversive learning and no aversive learning induced enhancement of CS responding was evident at memory retrieval (**Fig. 3g,h,k,l**). Furthermore, during aversive training, CS and US responses did not change significantly in either cell population (**Extended Data Fig. 10d,e,f,g,h,i**). These results indicate that repeated CS pre-exposure reduces auditory CS-evoked calcium responses in the CnF-input and CeA-output alone LA/B neurons and blocks aversive learning induced enhancement in CS-evoked responding in these cell populations.

Using the splitCre-PINCER approach, we next expressed GCaMP6s in the CnF-input/CeA-output neurons in LA/B and implanted an optical fiber in LA/B (**Fig. 3d and Extended Data Fig. 9c and Extended Data Fig. 10c**). Importantly, unlike the CnF-input or CeA-output alone neurons, the CnF-input/CeA-output neurons in LA/B did not respond to auditory CSs during the pre-exposure session (**Fig. 3m,n and Extended Data Fig. 9h,i**). Moreover, there was no significant increase in CS responding in the input-output defined neurons at memory retrieval (**Fig. 3o,p**) Finally, this cell population did not show any response to the CS but did respond to US during the conditioning session (**Extended Data Fig. 10j,k,l**). These results indicate that the CnF-input/CeA-output neurons in LA/B do not have the auditory CS-evoked calcium responses throughout the CS pre-exposure session and that latent inhibition blocks aversive learning induced enhancement of CS responding.

Taken together, these data show that the calcium responses in the CnF-input and CeA-output alone neurons in LA/B encode both the salience of the auditory CS and aversive memory formation, while the CnF-Input/CeA-output LA/B neurons selectively represent aversive memory and do not participate in salience processing. To test more explicitly whether coding in the input-output defined neurons was distinct from neural processing in the input or output alone defined cell populations, we trained linear support vector machine decoders on the calcium response patterns during pre-exposure, pre-conditioning, aversive conditioning and retrieval from the latent inhibition paradigm and then tested their ability to decode neural activity patterns comparing CnF-input vs. CnF-input/CeA-output, CnF-input vs. CeA-output and CeA-output vs. The CnF-input/CeA-output defined cell classes (**Extended Data Fig. 11a and Fig. 3q**). The decoder had significantly high decoding accuracy when classifying the input-output neuronal population dynamics from the input-alone or output-alone. In contrast, the decoder could not distinguish the activity patterns of input vs. output alone populations (**Fig. 3r**). To determine the response components which contributed most to the decoding of the input-output cells compared to the input-alone or output-alone, we calculated the weights of distinct task evoked activity patterns (e.g. CS responses across pre-exposure, pre-conditioning, conditioning and retrieval and US responses during conditioning). For distinguishing between input-output vs. input cell populations, CS-evoked activity in the first block of CS pre-exposure (Pre1 CS) and US-evoked responses during the conditioning period (Cond. US) had significantly higher weights compared to other epochs (**Extended Data Fig. 11b**), likely due to the fact that the input-alone, but not the input-output defined cells were auditory responsive prior to conditioning. Similarly, the decoding weights to classify the input-output and output alone populations were significantly higher during CS-evoked responses in the first block of CS pre-exposure (Pre1 CS) periods compared to other epochs (**Extended Data Fig. 11c**). These data demonstrate that the task related calcium dynamics of the CnF-input/CeA-output neurons in LA/B are mathematically separable from the activity patterns of the CnF-input alone or CeA-output alone neurons in LA/B, indicating a unique coding property of the CnF-input/CeA-output neurons in LA/B. Together, these findings reveal that LA/B cells defined by both their anatomical inputs and outputs specifically encode aversive memory formation, while cells defined by their input or output alone more broadly encode both salience and aversive learning and memory.

### PINCER reveals a specific aversive behavioral function of input-output defined cells

Given the distinct coding properties of the input-alone and output-alone cell populations from the input-output defined cells, we next used an optogenetic approach^1,7^ to determine whether they served similar or distinct behavioral functions during aversive and attentional learning and memory tasks. To do this, we first targeted the CnF-input alone population by expressing the inhibitory archaerhodopsin ArchT^59^ fused with a fluorescent protein in this LA/B cell population. Transneuronal anterograde rAAV1-hSyn-Cre and non-transneuronal rAAV5-CAG-cDIO-ArchT-tdTomato (as an experimental group) or rAAV5-CAG-cDIO-tdTomato (as a control group) viruses were injected in CnF and LA/B, respectively followed by placement of optical fibers in LA/B (**Fig. 4b**). Because of the importance of aversive stimulus evoked responding in LA/B neurons for producing associative learning^60^, laser illumination (589 nm) was applied through optical fibers bilaterally implanted in LA/B to inhibit neural activity during the US period of conditioning (**Fig. 4a**). Whereas this manipulation did not affect freezing behavior during aversive conditioning (**Fig. 4c**), it caused a significant reduction in freezing behavior during subsequent memory retrieval in the experimental group compared to the control group (**Fig. 4d**). This observation demonstrates that shock-evoked activity in the CnF-input neurons in LA/B is necessary for aversive memory formation.

**Fig. 4.**
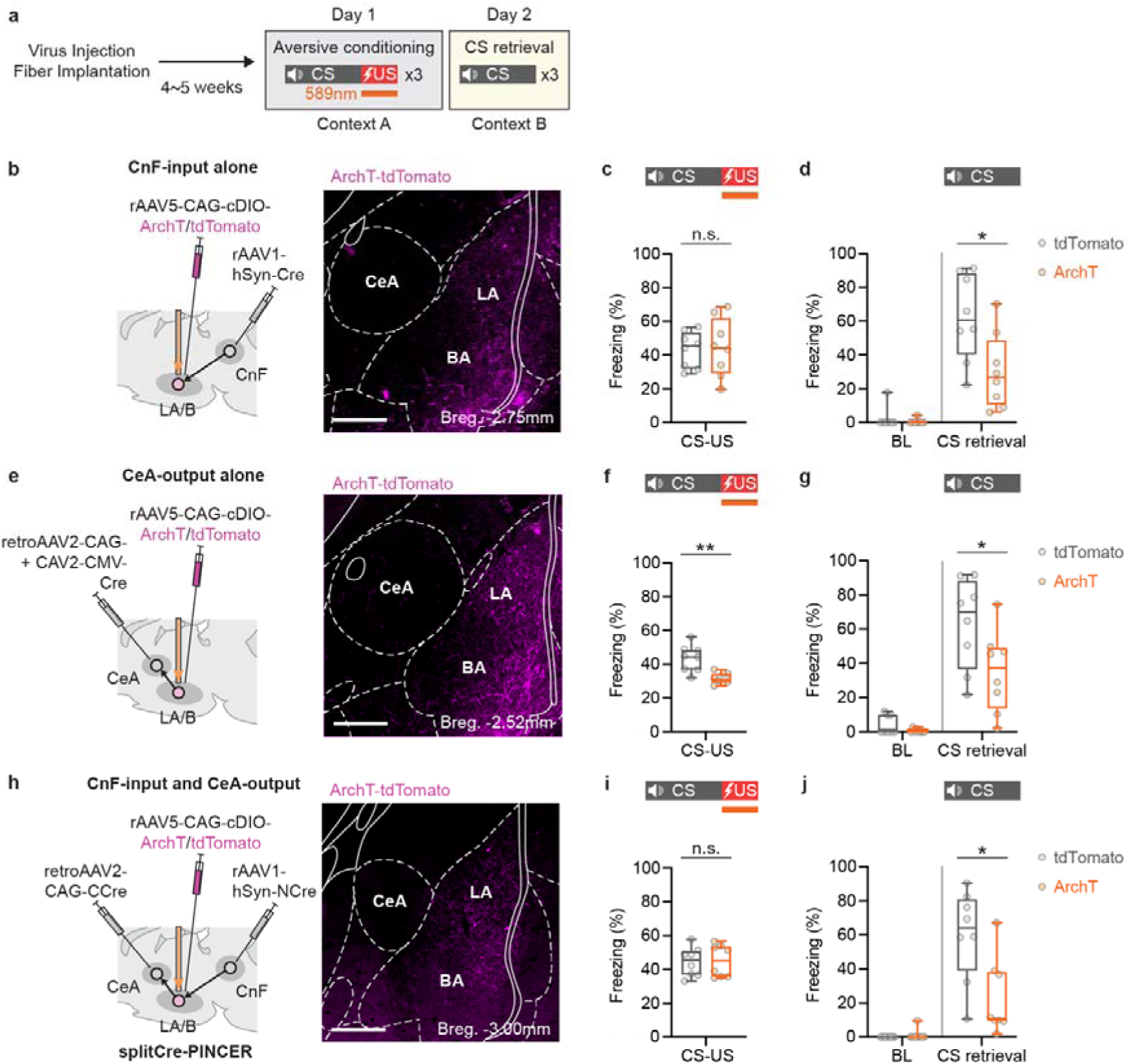
Anatomical connectivity-defined population-specific optogenetic inhibition effects on aversive conditioning. **a,** Experimental design for optogenetic manipulation experiments during aversive conditioning. **b-d,** Inactivation of the CnF-input alone neurons during shock presentation impairs aversive associative memory formation. **b,** Schematic (*left*) of viral combinations to express ArchT-tdTomato in the CnF-input alone neurons in LA/B for optogenetic manipulation. Representative image (*right*) of ArchT-tdTomato expression in the CnF-input alone neurons. **c,** Bar graphs show freezing responses to CS in aversive conditioning. No significant differences between tdTomato only control (N = 8 animals) and ArchT experimental (N = 8 animals) groups (*P* > 0.9999, Mann-Whitney test). **d,** Bar graphs show lower behavioral freezing responses during retrieval in the experimental group vs. control group (*P* = 0.0207, Mann-Whitney test). **e-g,** Inactivation of the CeA-output alone neurons during shock US presentation of aversive conditioning impairs associative learning and long-term memory formation. **e,** Schematic (*left*) of viral combinations to express ArchT-tdTomato in the CeA-output alone neurons in LA/B. Representative image (*right*) of ArchT-tdTomato expression in the CeA-output alone neurons. **f,** Bar graphs show reduced behavioral freezing responses during CS presentations in aversive conditioning in experimental (N = 8 animals) vs. control (N = 8 animals) groups (*P* = 0.0019, Mann-Whitney test). **g,** Bar graphs show lower CS-evoked behavioral freezing responses during retrieval in the experimental group vs. control group (*P* = 0.0379, Mann-Whitney test). **h-j,** Inactivation of the CnF-input/CeA-output neurons during shock presentation of aversive conditioning reduces associative memory formation. **h,** Schematic (*left*) of viral combinations to express ArchT-tdTomato in the CnF-input/CeA-output neurons in LA/B with the splitCre-PINCER approach. Representative image (*right*) of ArchT-tdTomato expression in the CnF-input/CeA-output neurons in LA/B. **i,** Bar graphs show no difference in CS-evoked behavioral freezing responses during aversive conditioning comparing tdTomato only control (N = 8 animals) and experimental (N = 8 animals) groups (*P* = 0.0379, Mann-Whitney test). **j,** Bar graphs show reduced CS-evoked behavioral freezing responses during retrieval in the ArchT group vs. tdTomato group (*P* = 0.0148, Mann-Whitney test). **b,e,h,** Scale bars: 500μm, **b,e,h,** Bregma distance in anterior-posterior direction. **c,d,f,g,i,j,** In box plot: center line, median; box limits, upper and lower quartiles; whiskers, two-sided 95^th^ percentile range of distribution. n.s.: *P* > 0.05, *: *P* < 0.05, **: *P* < 0.01.

Next, to express ArchT in the CeA-output alone neurons in LA/B, rAAV5-CAG-cDIO-ArchT-tdTomato (as an experimental group) or rAAV5-CAG-cDIO-tdTomato (as a control group) and a cocktail of retrograde retroAAV2-CAG-Cre and CAV2-CMV-Cre were injected in LA/B and CeA, respectively, combined with bilateral optical fiber placement bilaterally in LA/B (**Fig. 4e**). Unlike the manipulation of the CnF-input alone and the CnF-input/CeA-output neurons in LA/B (see below), optogenetically inhibiting output defined LA/B neurons during shock-US period of aversive conditioning significantly reduced freezing behaviors during both aversive conditioning (**Fig. 4f**) and retrieval (**Fig. 4g**) in the experimental group compared to the control group. This result indicates that the CeA-output neuron population in LA/B is important for both aversive learning and memory formation.

Finally, ArchT mediated optogenetic inhibition in the CnF-input/CeA-output neurons in LA/B was achieved through injections of non-transneuronal rAAV5-CAG-cDIO-ArchT-tdTomato (experimental group) or rAAV5-CAG-cDIO-tdTomato (control group) in LA/B using the splitCre-PINCER approach, and optical fibers were placed bilaterally in LA/B (**Fig. 4h**). Similar to the manipulation of the CnF-input alone neurons in LA/B, optogenetic inhibition during US presentation of training resulted in no effect on freezing behavior during aversive conditioning (**Fig. 4i**) but significantly reduced freezing behavior during subsequent memory retrieval in the experimental group compared to the control group (**Fig. 4j**). A similar outcome was obtained when the CnF-input/CeA-output neurons in LA/B were targeted using the Cre/Flp-PINCER method (**Extended Data Fig. 12**). These observations together suggest that the CnF-input, CeA-output and CnF-input/CeA-output defined LA/B neuronal populations are necessary for long-term aversive memory formation, while the CeA-output defined cells serve both learning and memory functions.

We next examined whether the CnF-input/CeA-output neurons serve a distinct function for salience processing compared with the CnF-input and CeA-output alone defined neurons as suggested by the calcium imaging results. To do this, we examined the effects of optogenetic inhibition of the three anatomical connectivity defined cell populations in the latent inhibition of aversive conditioning paradigm. Based on the calcium imaging findings showing that the CnF-input and CeA-output alone defined cells specifically encode salience, we hypothesized that inhibition of these cell types would reduce the effects of latent inhibition on aversive memory formation, but that inhibition of the CnF-input/CeA-output cells would have no effect.

To test this hypothesis, optogenetic inhibition in LA/B was applied during each CS during the 2 pre-exposure sessions prior to aversive conditioning (**Fig. 5a**). ArchT and tdTomato were expressed in the CnF-input or CeA-output alone neurons in LA/B using the same methods as the manipulation experiments in aversive conditioning (**Fig. 5b,f**). Laser inhibition of the input or output-alone defined LA/B neurons did not produce freezing behavior in the pre-exposure sessions (**Fig. 5c,g**). However, optogenetic inhibition of the CnF-input alone defined cells during pre-exposure increased freezing during subsequent aversive conditioning and memory retrieval (**Fig. 5d,e**), but the same manipulation in the CeA-output alone defined neurons selectively produced recovery of freezing during retrieval (**Fig. 5h,i**). These results indicate that the CnF-input and CeA-output alone neurons in LA/B are necessary for latent inhibition of aversive conditioning.

**Fig. 5.**
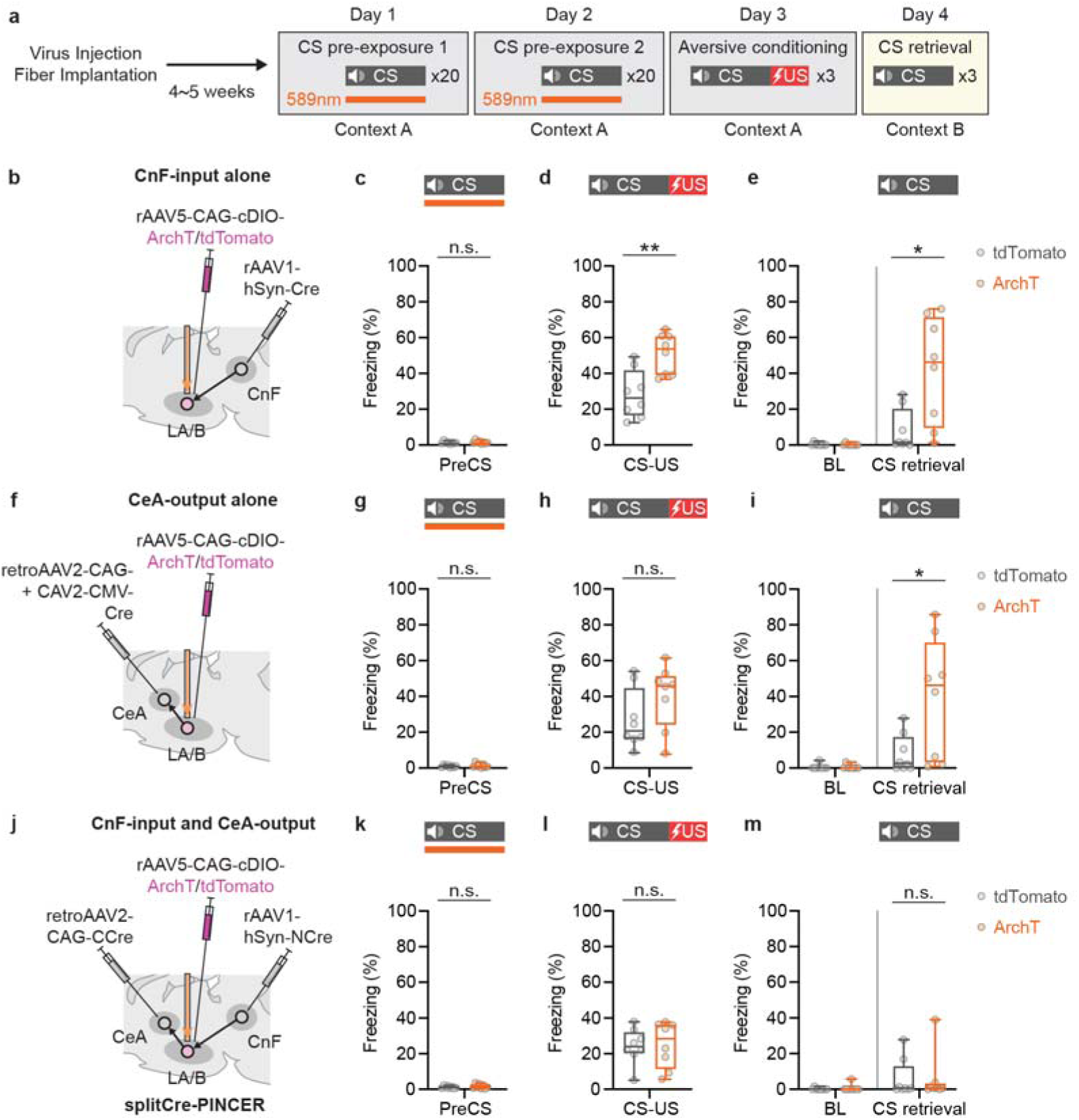
Anatomical connectivity-defined population-specific optogenetic inhibition effects on behavioral models of salience. **a,** Experimental design of optogenetic manipulation in latent inhibition of aversive conditioning in which optogenetic inhibition of LA/B cell populations was applied during auditory CS pre-exposure presentations. **b-e,** Inactivation of the CnF-input alone neurons during CS pre-exposure presentation reverses latent inhibition effects on aversive learning and memory. **b,** Schematic of viral combinations to express ArchT-tdTomato in the CnF-input alone neurons in LA/B for optogenetic manipulation. **c-e,** Bar graphs showing that optogenetic inhibition during CS presentation of pre-exposure does not affect behavioral freezing responses during **c,** pre-exposure (tdTomato N = 8 animals vs. ArchT N = 8 animals, *P* = 0.5545, Mann-Whitney test), but does enhance learned CS-evoked freezing responses during subsequent **d,** aversive conditioning (*P* = 0.0047, Mann-Whitney test) and **e,** memory retrieval (*P* = 0.0207, Mann-Whitney test) in experimental compared to control groups. **f-i,** Inactivation of the CeA-output alone neurons during CS pre-exposure presentation reduces latent inhibition effects on the formation of aversive associative memories. **f,** Schematic of viral combinations to express ArchT-tdTomato in the CeA-output alone neurons in LA/B. **g-i,** Bar graphs showing that optogenetic inhibition during CS presentation of pre-exposure does not affect behavioral freezing responses during **g,** pre-exposure (tdTomato N = 8 animals vs. ArchT N = 8 animals groups, *P* = 0.8785, Mann-Whitney test) or learned freezing responses during subsequent aversive conditioning **h,** (*P* = 0.2786, Mann-Whitney test), but does restore CS-freezing during **i,** memory retrieval (*P* = 0.0494, Mann-Whitney test) in the experimental vs. control groups. **j-m,** Inactivation of the CnF-input/CeA-output neurons during CS pre-exposure presentation does not affect latent inhibition effects on aversive associative learning and memory. **j,** Schematic of viral combinations to express ArchT-tdTomato in the CnF-input/CeA output neurons in LA/B with splitCre-PINCER approach. **k-m,** Bar graphs showing that optogenetic inhibition during CS presentation of pre-exposure does not affect behavioral freezing responses during pre-exposure (tdTomato N = 8 animals vs ArchT N = 8 animals, *P* = 0.8785, Mann-Whitney test) or learned freezing responses during subsequent **l,** aversive conditioning (*P* = 0.7984, Mann-Whitney test) or CS-freezing during **m,** memory retrieval (*P* = 0.7918, Mann-Whitney test) in the experimental vs. control groups. In box plot (**c-e,g-i,k-m**): center line, median; box limits, upper and lower quartiles; whiskers, two-sided 95^th^ percentile range of distribution. n.s.: *P* > 0.05, *: *P* < 0.05, **: *P* < 0.01.

Finally, we used the SplitCre-PINCER approach to express ArchT-tdTomato in the CnF-input/CeA-output neurons in LA/B for optogenetic inhibition during pre-exposures sessions of latent inhibition (**Fig. 5a,j**) as was done for the CnF-input and CeA-output alone neurons. In contrast to the effects seen with manipulation of the CnF-input and CeA-output alone neurons, the inhibition of the CnF-input/CeA-output neurons in LA/B during CS pre-exposure did not produce any effects on freezing behavior during CS pre-exposure (**Fig. 5k**), aversive conditioning (**Fig. 5l**) or retrieval (**Fig. 5m**). This result shows that CS-evoked activity in the CnF-input/CeA-output neurons in LA/B is not required for latent inhibition of aversive conditioning.

Taken together, these optogenetic inhibition experiments demonstrate that the CnF-input alone and CeA-output alone LA/B neurons participate in broad functions including salience processing and aversive learning and memory, while the CnF-input/CeA-output neurons in LA/B are required specifically for aversive memory formation.

### PINCER reveals differential coding of appetitive learning in distinct connectivity defined cell types

As LA/B is known to be involved in not only aversive but also appetitive form of emotional learning^12,15,25,34–38,40,41,61^, we next used the PINCER technique to record neural activity in the input-output projection-defined LA/B cell population and compared this to the dynamics of the input or output alone populations during an appetitive (reward) learning task.

In the appetitive task, CSs alone were presented during a pre-conditioning phase, followed by a conditioning session in which CS presentation was paired with sucrose reward (US) presented at a reward port. The conditioning session were repeated twice per day for two days (4 sessions in total) (**Fig. 6a**). To gain the reward, food-deprived animals learned to approach the reward port (reward-seeking) during the CS period over the course of the conditioning sessions, which resulted in an increase in the probability of port entry during the CS in session 4 compared to session 1 (**Fig.1b and Extended Data Fig.13a**)

**Fig. 6.**
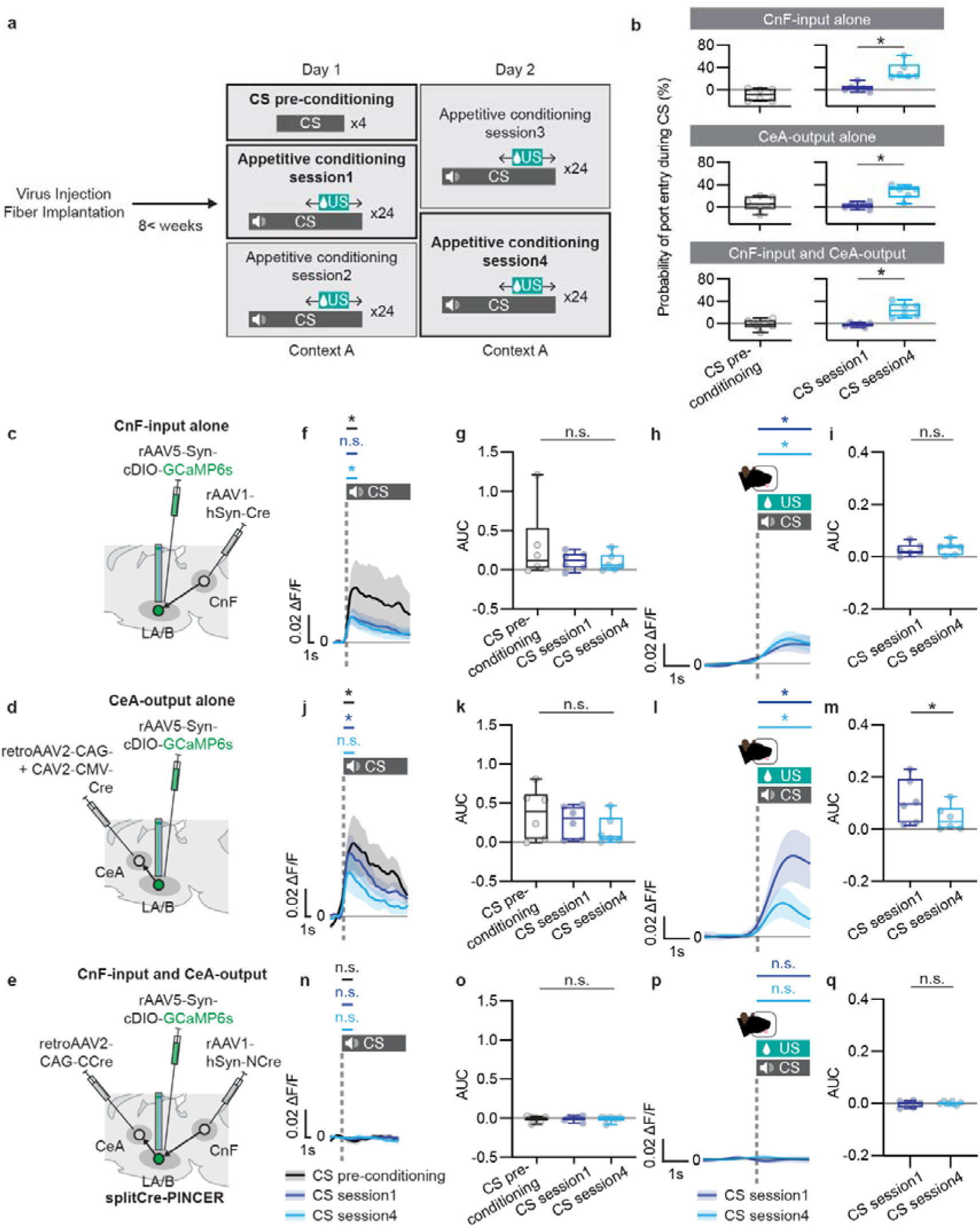
Calcium imaging during appetitive learning in anatomical connectivity defined cell types using PINCER. **a,** Schematic of experimental design. **b,** Development of reward-seeking behavior during CS presentation in appetitive conditioning. Normalized nose-poke probability (see Methods) was used as an index of reward-seeking behavior to quantify appetitive learning. Animals for calcium imaging of CnF-input alone, CeA-output alone and CnF-input/CeA-output neurons in LA/B developed significantly higher probability of nose-poke behaviors during CS presentation in session 4 (cyan) in comparison to session 1 (blue) (N = 6 animals for each of the 3 anatomical connectivity deveined group, *P* = 0.0312 for all of the 3 anatomical connectivity deveined groups, Wilcoxon test). **c-e,** schematic showing viral combinations to express GCaMP6s in the **c,** CnF-input alone, **d,** CeA-output alone and **e**, CnF-input/CeA-output neurons in LA/B during appetitive conditioning. **c,f-i,** CS- and reward acquisition-evoked calcium responses of the CnF-input neurons in LA/B during appetitive conditioning. **f,** peri-event time histogram (PETH) showing group averaged CS-evoked calcium responses in the pre-conditioning (black) compared with the session 1 (blue) and session 4 (cyan) of appetitive conditioning (N = 6 animals). CS responses in pre-conditioning (black) and session 4 (cyan) but not session 1 (blue) of appetitive conditioning were significantly higher than baseline periods prior to CS onset (2 seconds before vs. after CS onset, pre-conditioning *P* = 0.0312, session 1 *P* = 0.0625 and session 4 *P* = 0.0312, Wilcoxon test by corresponding baselines prior to CS onsets). **g,** Bar graphs show no differences in CS-evoked ΔF/F AUCs comparing pre-conditioning (black), session 1 (blue) and session 4 (cyan) (*P* = 0.9563, Friedman test). **h,** PETH showing group averaged reward acquisition-evoked calcium responses during session 1 (blue) and session 4 (cyan) of appetitive conditioning exhibited increased responses to reward compared to calcium dynamics prior to reward acquisition onset (2 seconds before vs. after reward acquisition onset, *P* = 0.0312 for both session 1 and session 4). **i,** Bar graphs show no differences in reward acquisition-evoked ΔF/F AUCs comparing session 1 (blue) and session 4 (cyan) (*P* = 0.2188, Wilcoxon test). **d,j-m**, CS- and decreased reward acquisition-evoked calcium responses of the CeA-output neurons in LA/B during appetitive conditioning. **j,** PETH showing group averaged CS-evoked calcium responses in the pre-conditioning (black) compared with the session 1 (blue) and session 4 (cyan) of appetitive conditioning (N = 6 animals). CS responses in pre-conditioning (black) and session 1 (blue) but not session 4 (cyan) of appetitive conditioning were significantly higher than baseline prior to CS onset (*P* = 0.0312 for pre-conditioning and session 1 and *P* = 0.0625 for session 4, Wilcoxon test by corresponding baselines prior to CS onsets). **k,** Bar graphs show no differences in CS-evoked ΔF/F AUCs comparing pre-conditioning (black), session 1 (blue) and session 4 (cyan) (*P* = 0.4297, Friedman test). **l,** PETH showing group averaged reward acquisition-evoked calcium responses during session 1 (blue) and session 4 (cyan) of appetitive conditioning exhibited increased responses to reward compared to calcium dynamics prior to reward acquisition onset (*P* = 0.0312 for both session 1 and session 4, Wilcoxon test). **m,** Bar graphs show decrease in reward acquisition-evoked ΔF/F AUCs in session 4 (cyan) in comparison to session 1 (blue) (*P* = 0.0312, Wilcoxon test). **e,n-q,** No CS- and reward acquisition-evoked calcium responses of the CnF-input/CeA-output neurons in LA/B observed during appetitive conditioning. **n,** PETH showing group averaged CS-evoked calcium responses in the pre-conditioning (black) compared with the session 1 (blue) and session 4 (cyan) of appetitive conditioning (N = 6 animals). These neurons did not respond significantly to CS during pre-conditioning (black), session 1 (blue) and session 4 (cyan) compared with baseline activity levels (pre-conditioning *P* = 0.6875, session 1 *P* = 0.3125 and session 4 *P* = 0.5625, Wilcoxon test). **o,** Bar graphs show no differences in CS-evoked ΔF/F AUCs comparing pre-conditioning (black), session 1 (blue) and session 4 (cyan) (*P* = 0.9563, Friedman test). **p,** PETH showing group averaged reward acquisition-evoked calcium responses during session 1 (blue) and session 4 (cyan) of appetitive conditioning demonstrate no response to reward compared to calcium dynamics prior to reward acquisition onset (session 1 *P* > 0.9999 and session 4 *P* = 0.6875, Wilcoxon test). **q,** Bar graphs show no differences in reward acquisition-evoked ΔF/F AUCs comparing session 1 (blue) and session 4 (cyan) (*P* = 0.8438, Wilcoxon test). **f,h,j,l,n,p,** Shades: SEM. **b,g,i,k,m,o,q,** In box plot: center line, median; box limits, upper and lower quartiles; whiskers, two-sided 95^th^ percentile range of distribution. n.s.: *P* > 0.05, *: *P* < 0.05.

Using the transneuronal anterograde or retrograde viral techniques, GCaMP6s was expressed in the CnF-input alone, CeA output alone or CnF-input/CeA-output neurons in LA/B along with an optical fiber in LA/B for fiber photometry recordings from these cell populations (**Fig. 6c,d,e**). The CnF-input alone and CeA-output alone neurons showed responses to the CS during pre-conditioning, session 1 and session 4 of appetitive conditioning (**Fig. 6f,j**) but displayed no significant decrease in CS-evoked responding throughout pre-conditioning, session 1 and session 4 (**Fig. 6g,k**). These anatomical connectivity defined LA/B cell types also responded when animals nose poked for reward (reward acquisition) during conditioning (**Fig. 6h,l**). While the CnF-input alone neurons in LA/B did not show difference in reward acquisition-evoked US responses, the CeA-output alone neurons in LA/B decreased the response in session 4 compared to session 1 (**Fig. 6i,m**). By contrast, the LA/B CnF-input/CeA-output neurons did not exhibit CS responses during pre-conditioning, session 1 or session 4 of appetitive conditioning (**Fig. 6n,o**) and did not respond during reward acquisition (**Fig. 6p,q**).

To further examine whether the lack of responses to CS and US in the CnF-input/CeA-output neurons in LA/B is attributable to an absence of functional involvement in this appetitive task, we used the SplitCre-PINCER approach to express ArchT-tdTomato in the CnF-input/CeA-output neurons in LA/B for optogenetic inhibition during sucrose reward presentations during learning (**Extended Data Fig. 13b,c**). The inhibition of the CnF-input/CeA-output neurons in LA/B during US presentation had no effect on the development of reward seeking behavior from session 1 to session 4 during conditioning (**Extended Data Fig. 13d,e**). This result shows that reward acquisition-evoked activity in the CnF-input/CeA-output neurons in LA/B is not required for reward learning.

Taken together, these results indicate that the CnF-input and CeA-output alone LA/B neurons respond to rewards and that reward learning maintains CS responding in these cells, but does not increase it. However, these findings indicate that CnF-Input/CeA-output LA/B neurons do not encode or participate in appetitive learning. Combined with the previous results, this shows that CnF-Input/CeA-output LA/B neurons serve a specific aversive learning function, while input and output alone defined cell populations more broadly encode salience, aversive and reward learning.

## Discussion

### Principles of the PINCER technique

The functionality of brain circuits emerges from the complex patterns of connections between neurons across different brain regions. A key objective of neuroscience research is to understand how the functions of the specific cells embedded in these circuits arise from their anatomical connections. In this study, we introduce the PINCER technique to address this question. We developed two different PINCER approaches using split-Cre or dual recombinases and show the effectiveness of these approaches in labeling, manipulating and imaging from input-output defined cell populations and for comparing their properties to cells defined by their input or output alone. We find that the input-output defined cell types serve more specific neural coding and emotional processing functions compared with LA/B neurons defined by the input or output alone. Specifically, the input-output defined LA/B neurons specifically code for and participate in aversive memory formation whereas LA/B neurons defined by the input or output alone serve more general encoding and behavioral functions including aversive learning, memory formation and salience processing. These findings reveal key principles of brain region organization in which cells defined by their afferent input or efferent output alone serve broader computational roles while neurons defined by both input and output participate in more specific functions.

### Integrating PINCER into the existing neural circuit-based toolkit

Combining PINCER with existing neuroscience techniques will provide deeper insights into the organization of neural circuits. For example, the PINCER technique serves a distinct but complementary role to the ‘tracing the relationship between input and output’ (TRIO/cTRIO) approach^62^. The TRIO/cTRIO technique has been a powerful viral tool for anatomical dissection of input-output circuit organization. A recent study expanded its utility to include calcium imaging and optogenetic manipulation^44^. However, different from PINCER, the TRIO/cTRIO technique targets cells based on their projections to distinct target structures and is used to map the afferent inputs onto the labeled cells from throughout the brain. However, it does not allow for cell type identification based on combinatorial input-output connectivity but could be applied in a complementary way with PINCER. For example, TRIO could be used in anatomical studies to determine brain-wide afferent inputs onto different projection defined cell populations in a given brain region^62–66^ and PINCER could then be employed to target inputs that are enriched in one of the projection defined target populations to selectively capture that input-output defined cell type for functional characterization. The principles of PINCER can also be combined with the GRASP synaptic labeling technique to study synaptic connectivity and strength of inputs onto output defined projection cells^67,68^. Together with existing approaches, PINCER has the potential to significantly advance research by providing more detailed insights into the anatomical and functional input-output organization of neural circuits.

### Molecular cell type identification characterized by PINCER

How to define cell types in the brain is a central topic of discussion in the neuroscience field^69–71^. Cell types have been characterized based on molecular identity^72–74^ as well as afferent inputs^75,76^ and output-projection labeling^7^ and the combination of molecular identify and efferent projection^38^. PINCER offers an approach to define cell types by their input-output connectivity as well as their molecular makeup. In this study, we characterized and compared the molecular-neurotransmitter identities of the CnF-input alone, CeA-output alone and CnF-input/CeA-output defined LA/B neurons. Whereas most of the input-output defined LA/B neurons are *vGluT*+ glutamatergic excitatory neurons, the input-alone and output-alone defined LA/B neurons are mixtures of *vGluT*+ glutamatergic excitatory and *vGAT*+ GABAergic inhibitory neurons. This observation suggests that the input-output defined LA/B neurons represent glutamatergic long-range projection cells while a portion of the input-alone LA/B neurons form interneuron microcircuits inside LA/B and a subset of CeA projection cells are GABAergic. Although we have characterized the expression of the main sub-classes^71,72^ of GABAergic interneurons including SST+, PV+ and VIP+ neurons, characterization of other potential excitatory neuronal subtypes based on developmental lineage specifications^73,74^ will be important in future investigations.

### Amygdala cell type specific coding and behavioral functions revealed by PINCER

A key function of LA/B is to associate sensory stimuli with aversive or rewarding outcomes and store a memory of these associations through plasticity of sensory input synapses onto amygdala neurons^20,23,24,26,77^. LA/B neurons encode aversive and rewarding stimuli in partially separable populations of cells and many respond to sensory stimuli prior to learning^15,25,30,34,61,78,79^, particularly when they are unexpected^16^. Thus, two key features of emotional processing, salience and valence, are encoded in LA/B neurons, prompting the question of whether and how these features relate to biologically defined cell types in amygdala. Indeed, some recent work has suggested that valence is represented in population activity and is not encoded in a selected population of response-defined LA/B neurons^30,31^. Attempts to map functional coding features to output-alone or molecularly defined cell types have had mixed results, with some studies reporting limited enrichment of aversive or reward coding in specific cell types while others find mixed selectivity in response properties^34–38,40,41,79,80^. Many LA/B neurons respond to sensory stimuli prior to conditioning and reducing their responses with repeated presentations^11,12,16,25,61^ suggesting a salience function, but the functional role of LA/B cell types in salience has not been examined and the general role of LA/B in salience processing is unclear. Using PINCER, we examined these questions and find that the input or output alone defined cell types support both salience and valence functions, responding to aversive and rewarding outcomes as well as sensory stimuli prior to learning and serving both salience and aversive learning functions for behavior. In contrast to the broad functional features of these cell groups, the application of PINCER demonstrated that CnF-input/CeA-output defined cells selectively encode and participate in aversive learning **(Fig. 2**, **Fig. 3**, **Fig. 4, and Fig. 5)**.

These observations on the coding and functional properties of neurons in the CnF→LA/B→CeA circuit suggest a hypothetical circuit model for how valence and salience are integrated in different amygdala cell types for emotional processing (**Extended Data Fig. 14a**). During aversive learning (**Extended Data Fig. 14b**), noxious information is conveyed from CnF to the CnF-input receiving LA/B neurons (which include some CeA projecting input-output cells) to instruct aversive learning in cells that also receive sensory CS inputs. Notably, the input-output defined cells do not respond to rewards or CSs paired with rewards and are not required for appetitive conditioning, demonstrating that they specifically function for aversive learning. Recent results support this model by showing that CnF conveys aversive information about the sensory properties of innately aversive experiences and bodily reactions related to these events to LA/B and functions to trigger aversive memory formation^51^. Following aversive conditioning, auditory evoked responses are enhanced in LA/B neuronal subtypes through synaptic plasticity occurring at thalamic/cortical inputs or in local recurrent connections between these cell types^81^. During memory retrieval, these inputs are now strong enough to drive output to CeA to produce conditional freezing responses. The CnF-input and CeA-output alone neurons also respond to rewards and CSs and do not significantly change their CS responding during repeated presentations over the course of reward learning (**Fig. 6**).

Beyond an understanding of how aversive and appetitive learning and salience processing is allocated across anatomically defined cell types, our findings also provide insights into the functional role and mechanisms of salience processing in LA/B neurons. Latent inhibition is considered a behavioral model of salience based on the idea that sensory stimuli lose the ability to attract attention with repeated presentations during pre-exposure, thereby impairing subsequent associative learning. Brain regions such as nucleus accumbens (NAc), hippocampus, amygdala and ventromedial prefrontal cortex (vmPFC) have been implicated in latent inhibition^21,58,82^. Early studies reported that LA/B lesions^14^ or infusion of an NMDA receptor antagonist in LA/B^83^ prior to pre-exposure disrupted latent inhibition, allowing animals to acquire normal associative learning. However, how LA/B neuronal coding is linked to function and cell types during latent inhibition was not known. We found that the CnF-input or CeA-output alone LA/B neurons showed strong responses to the initial presentation of the CS, but that their responses diminished with repeated CS presentations **(Fig. 3f,j and Extended Data Fig. 9e,g**). Furthermore, learning did not enhance CS responses following latent inhibition **(Fig. 3h,l)**. Moreover, optogenetic inhibition of the CnF-input or CeA-output alone cell populations during CS pre-exposure blocked the impairing effect of latent inhibition on learning **(Fig. 5d,e,i)**. During reward learning, input and output alone populations respond to rewards and, despite repeated presentations, do not decrease their CS responding during appetitive conditioning. The reward responding and diminished habituation of CS responding suggests that these cell populations encode reward and/or salience, but possibly not appetitive learning as they do not increase CS responding during learning. By contrast, the input-output defined cells did not respond to sensory stimuli prior to aversive learning **(Fig. 3m and Extended Data Fig. 9h)** and optogenetically inhibiting them did not affect latent inhibition **(Fig. 6l,m)**, though learning related increases in CS-responding **(Fig. 2j)** were reduced **(Fig. 3p)**. This suggests that these cells do not encode salience but do reflect the effect of latent inhibition on aversive learning.

Together, these results suggest a model for how salience and aversive learning functions can be partially segregated in LA/B cell types but integrated across the larger LA/B neuronal population (**Extended Data Fig. 14c**). During latent inhibition, repeatedly presented sensory stimulus activate the CnF-input and CeA-output alone LA/B neurons and the activation of these cells is reduced with repeated CS presentations. This reduction in CS processing reduces the ability of LA/B neurons to undergo plasticity during conditioning, thereby blocking aversive memory formation. Intriguing questions raised by these results relate to the mechanism through which the habituation response during latent inhibition occurs and the causal role of this habituation in impairing subsequent learning. Because postsynaptic activity is required in the CnF-input and CeA-output alone neurons to produce latent inhibition and the fact that NMDA receptor activation in LA/B is also necessary for this effect^83^, one possibility is that this occurs through NMDA receptor mediated long-term depression (LTD)^84^ of auditory inputs to these cell populations. Alternatively, long-term potentiation (LTP)^84^ of auditory inputs to the GABAergic CnF-input alone and CeA-output alone LA/B interneurons that project to other input or output defined cells could underlie this effect. More detailed analysis of the plasticity mechanisms occurring in specific LA/B cell types during latent inhibition will be important in future studies.

### Limitations of the PINCER technique and how to overcome them

The usage of low-toxicity and relatively high anterograde labeling efficiency rAAV1^3^ for PINCER is an effective strategy for labeling input-defined neurons in unidirectional input pathways such as the CnF to LA/B projection^51^. However, the utility of rAAV1 for reciprocally connected circuits can be limited due to the retrograde transfection leakage of rAAV1^3^. This can be overcome using other anterograde transneuronal approaches such as the yellow fever vaccine YFV-17D^85^ or ATLAS^86^. The development of a less-toxic version of the anterograde transneuronal viruses HSV1-H129ΔTK^8^ and VSV^9^ and the integration of these new viruses into the PINCER method may be another effective alternative. In addition, while rAAV1 may have retrograde leakage, if the efferent output cell population being targeted in PINCER does not project back to the site where the anterograde AAV is injected, this approach would still be viable (**Extended Data Fig. 15**).

Comparing the two PINCER approaches, there are clear advantages to the splitCre PINCER version such as its faster incubation time **(Extended Data Fig. 2c,d)**. In addition, the use of a single recombinase opens the possibility of combining it with another recombinase (e.g. Flp) to triple label cells based on input, output and a third factor such as molecular identity or a second input or output connection. However, there are some advantages to using the Cre/Flp-PINCER technique in some applications. Specifically, there can be tropism with retroAAV2 in the splitCre PINCER approach, limiting its utility in some circuits^6^. Because of this, we also developed the Cre/Flp-PINCER approach to deliver Cre-dependent Flp retrogradely using a cocktail of retroAAV2 and CAV2 (Cre/Flp-PINCER), as these two retrograde viruses can have different tropism^87,88^. Although this tropism was not apparent in the circuits we studied here as both PINCER approaches labeled similar numbers of output neurons in LA/B (**Extended Data Fig. 2c,d**), this may not be true in other circuits where retroAAV2 might not be effective. In these instances, Cre/Flp-PINCER may be a better option when combined with other retrograde viruses including CAV or newer, lower toxicity rabies viruses^89–91^.

### Future applications of PINCER-based techniques

An important technical extension of PINCER could be to combine it with cell type specific molecular profile targeting. Currently we use PINCER to label cells defined by anatomical inputs and outputs, but it could also be used to target neurons based on combinations of input (through anterograde transneuronal viruses) and molecular markers in specific downstream cells. Alternatively, cells defined by inputs, outputs and molecular profile could be labeled using triple conditional intersectional strategies^92–94^ in which PINCER is used with molecular marker specific recombinase lines or cell-type specific promoter/enhancer/viral transduction^95–97^. PINCER could also be combined with engram labeling techniques such as the TRAP approach^35,98^ or calcium activity based FLiCRE/Cal-Light^99,100^ to capture cells based on their inputs, output and response properties. These technical extensions would broaden the PINCER toolset and allow for even finer cell type identification.

Current cell type-based identification relies on clustering neuronal subtypes based on the similarity of the transcriptomes^101,102^ of individual cells. However, neuronal function is mostly defined by the combined activity of all the cells they receive afferent input from and all the brain regions they project to. Thus, the goal of neuroscience is to determine the function and coding properties of neuronal subtypes defined by their comprehensive input-output connectivity. PINCER represents a first step in this direction. Future advances in neuronal tracing and labeling technology may allow for the expansion of PINCER to achieve this broader goal.

## Acknowledgements

We thank Yuri Sugiyama, Buddhini Wimarsha Jayathilake, Tomoya Duenki, Yuki Goya and Nisha Jose for technical assistance. We thank Dr. Andrew Murray for gifting the Neuro2A cell lines used for packaging CVS-strain rabies virus. We are grateful to the RIKEN CBS-EVIDENT Open Collaboration Center for imaging equipment, software and the technical assistance with confocal image acquisition. We also thank Nur Zeynep Gungor and other Johansen lab members for advice on experimental design and data analysis.

## Author contributions

Y.K. and J.P.J. conceived the project. Y.K. and R.Y. designed and created the plasmids for PINCER technique. Y.K. designed and carried out the proof of principle and anatomical characterization experiments. Y.K. and X.G. designed and carried out the calcium imaging experiments, and Y.K., X.G. and T.O analyzed the calcium imaging data. Y.K., X.G. and A.U. designed and carried out the optogenetic experiments. Y.K., X.G., T.O and J.P.J. wrote the manuscript. J.P.J. and D.Y. supervised the work. All authors read and approved the final manuscript.

## Methods

### Animals

All experimental procedures were approved by the Animal Care and Use committee of the RIKEN Center for Brain Science. Male wild-type Long-Evans (lar: Long-Evans, Japan SLC, Inc) or SST-Cre (HsdSage: LE-*SSt*.*Cre^em1sage^*, inotiv but originally created at SAGE Labs) rats (8-10 weeks old at the time of the first surgery) were used in all experiments. Rats were singly housed under a 12-hour light/dark cycle. Food and water were provided *ad libitum*. For appetitive conditioning experiments, rats were food-deprived for 2 weeks of limited food access (6∼9g of food pellets per a day). The food-deprived rats were habituated to 30% sucrose water (30mL) on the first day of food deprivation. All behavioral experiments were performed during the light cycle.

### Plasmid Preparation

Recombinant DNAs for recombinases in splitCre-PINCER: DNA fragments encoding FLAG-GCN4cc-linker-NLS-NCre and Myc-GCN4cc-linker-NLS-CCre were amplified by PCR from pCMV-Tag2B-NCre^103^ (addgene plasmid #106368) and pCMV-Tag3B-CCre^103^ (addgene plasmid #106369) and inserted into between after hSyn promoter and before WPRE sequences in pAAV-hSyn plasmid or into between after CAG promoter and before WPRE sequences in pAAV-CAG plasmid using NEBuilder HiFi DNA Assembly Master Mix (New England Biolabs, E2321S) to produce pAAV-hSyn-NCre, pAAV-CAG-CCre, pAAV-hSyn-CCre and pAAV-CAG-NCre for following packaging into AAVs.

Recombinant DNAs for reporter genes:

DNA fragments encoding mtagBFP (gifted from Dr. Ian Wickersham) were amplified by PCR and inserted into between CAG promoter and WPRE sequences in pAAV-CAG plasmid using NEBuilder HiFi DNA Assembly Master Mix (New England Biolabs, E2321S) to produce pAAV-CAG-mtagBFP.

Recombinant DNAs for recombinase-dependent reporters and optogenetic genes:

DNA fragment of CBA promoter sequence was replaced with Ef1a promoter sequence sequences in pAAV-Ef1a-fDIO-eYFP^92^ (addgene plasmid #55641) and pAAV-CAG-fDIO-ArchT-eYFP^104^ plasmids to produce pAAV-CBA-fDIO-eYFP and pAAV-CBA-fDIO-ArchT-eYFP plasmids, respectively. Subsequently, DNA fragments encoding mCherry were amplified, inverted and replaced with inverted EYFP sequence after the promoter sequence, between F5 and FRT and before WPRE sequences in pAAV-CBA-fDIO-eYFP and pAAV-CBA-fDIO-ArchT-eYFP plasmids to produce pAAV-CBA-fDIO-mCherry and pAAV-CBA-fDIO-ArchT-mCherry, respectively. DNA fragments encoding mCherry were amplified, inverted and inserted after the promoter sequence, between loxP and lox2722 and before WPRE sequences in pAAV-CBA-cDIO-eGFP, a gift from Edward Boyden (Addgene plasmid #28304) to produce pAAV-CBA-cDIO-mCherry. pAAV-CAG-fDIO-eYFP were produced in our lab as described previously^104^. pAAV-CBA-cDIO-eGFP was a gift from Edward Boyden (Addgene plasmid #28304).

Recombinant DNAs for Cre recombinase-dependent Flp recombinase gene:

pAAV-CBA-cDIO-Flp was produced in our lab as described previously^66^.

Recombinant DNAs for helper AAV gene used in rabies tracing:

pAAV-CAG-cDIO-TVA was produced in our lab as described previously^105^. pAAV-CAG-cDIO-H2B-HA-2A-G^89^ was a gift from Thomas Jassel (Addgene plasmid #73477).

### Viruses

rAAV1-hSyn-Cre (pENN.AAV.hSyn.Cre.WPRE.hGH, a gift from James M. Wilson, Addgene viral prep #10553-AAV1), rAAV5-CAG-tdTomato (pAAV-CAG-tdTomato, a gift from Edward Boyden, Addgene viral prep #59462-AAV5), rAAV5-CAG-eGFP (pAAV-CAG-GFP, a gift from Edward Boyden, Addgene viral prep #37825-AAV5), rAAV5-CAG-cDIO-tdTomato (pAAV-FLEX-tdTomato, a gift from Edwad Boyden, Addgene viral prep #28306-AAV5), rAAV5-Syn-cDIO-GCaMP6s (pAAV.Syn.Flex.GCaMP6s.WPRE.SV40, a gift from Douglas Kim and GENIE Project, Addgene viral prep #100845-AAV5)^106^, rAAV5-CAG-cDIO-ArchT-tdTomato (pAAV-FLEX-ArchT-tdTomato, a gift from Edward Boyden, Addgene viral prep #28305-AAV5)^59^, retroAAV2-CAG-GFP (pAAV-CAG-GFP, a gift from Edward Boyden, Addgene viral prep #37825-AAVrg) and retroAAV2-CAG-tdTomato (pAAV-CAG-tdTomato, a gift from Edward Boyden, Addegene viral perp #59462-AAVrg) were purchased from Addgene.

CAV2-CMV-GFP^107^, CAV2-CMV-cDIO-eGFP^62^ and CAV2-CMV-Cre^108^ were obtained from the Montpellier vector core.

rAAV1-hSyn-NCre, rAAV1-hSyn-CCre, rAAV1-hSyn-Flp, rAAV5-CAG-mtagBFP2, rAAV5-CBA-cDIO-GFP, rAAV5-CBA-fDIO-mCherry, rAAV5-CAG-fDIO-eYFP^104^, rAAV5-CAG-fDIO-ArchT-mCherry, rAAV9-CAG-cDIO-TVA^105^, rAAV9-CAG-cDIO-H2B-HA-2A-G^105^, retroAAV2-CBA-cDIO-mCherry retroAAV2-CAG-CCre, retroAAV2-CBA-cDIO-Flp^54^, retroAAV2-CAG-NCre and retroAAV2-CAG-Cre were packaged in our lab. rAAV1-hSyn-NCre, rAAV1-hSyn-CCre and rAAV1-hSyn-Flp viruses with estimated titer >1×10^13^ GC/mL were used in experiments for anterograde trans-neuronal infection^3^. All the other viruses with estimated titer >1×10^12^ GC/mL were used in experiments

Plasmid for EnvA-coated CVS-strain glycoprotein-deleted Rabies virus fused with EGFP (EnvA-CVS-N2cΔG-eGFP)^89^ was purchased from addgene (RabV CVS-N2cΔG-eGFP, a gift from Thomas Jessel, Addgene plasmid #73461) and packaged in our laboratory^66,105^ using the Neuro2A cell lines (mouse neuroblasts) provided by Dr. Andrew Murray.

### Stereotaxic Surgery

All surgeries were conducted using isoflurane anesthesia (3% for induction, 1.5–2.5% for maintenance) in a stereotaxic apparatus (model 942, David Kopf Instruments). Viruses were administered via a stainless-steel injection cannula (28-gauge, Plastic One), which was connected to a 2 mL Hamilton syringe with polyethylene tubing. After advancing the virus-loaded cannula to the target site, a syringe pump (PHD2000, Harvard Apparatus) continuously infused the viruses at a flow rate of 0.07 μL/min. The cannula remained at the injection site for 15 minutes after the infusion was completed, before being slowly withdrawn. Immediately following the virus injection, an optical fiber (400 μm core, N.A. = 0.66, Doric Lenses) was implanted for *in vivo* calcium imaging experiments, while optical fibers held in ferrules (200 μm core, N.A. = 0.37, Doric Lenses) were used for *in vivo* optogenetic studies. These optical fibers were secured to the skull with stainless steel screws, super-bond, and acrylic dental cement. For all experiments except those involving rabies tracing (**Extended Data Fig. 5d,e,f**), viruses were injected into the target brain regions of wild-type rats, while rabies tracing experiments involved injections in SST-Cre rats.

For all proof of principle and anatomical characterization experiments, the following injection coordinates and viral volumes were used: CnF (AP: −8.3 mm from Bregma, ML: ±4.0 mm from Bregma, DV: −5.5 mm from pia with 20° lateral angle, 0.3 μL injection volume), LA/B (AP: −3.0 mm from Bregma, ML: +5.0 mm from Bregma, DV: −7.1 mm from pia, 1.0 μL injection volume), and CeA (AP: −2.3 mm from Bregma, ML: ±2.15 mm from Bregma, DV: −7.8 mm from pia with 14° medial angle, 0.3 μL injection volume).

For the experiment examining the existence of the CnF-LA/B-CeA circuit (**Extended Data Fig. 1d**), rAAV1-hSyn-Cre virus was bilaterally injected into CnF, rAAV5-CAG-cDIO-tdTomato virus was unilaterally injected into LA/B, and retroAAV2-CAG-eGFP and CAV2-CMV-eGFP and rAAV5-CAG-mTagBFP2 viruses (mixed in 1:1:1 ratio by volume) were bilaterally injected into CeA. Rats were singly housed for 4+ weeks to allow viral infection and expression and then terminated as described in the Histology section. Viral expression of mTagBFP in CeA was used to determine whether leakage of virus injected into CeA was apparent in LA/B (data not shown).

For the anterograde AAV-retrograde Cre-dependent fluorophore experiments (**Extended Data Fig. 1e**), rAAV1-hSyn-Cre virus was bilaterally injected into CnF, and retroAAV2-CBA-cDIO-mCherry and rAAV5-CAG-eGFP viruses (mixed in 1:1 ratio by volume) were bilaterally injected into CeA. Rats were singly housed for 4+ weeks to allow viral infection and expression and then terminated as described in the Histology section. Viral expression of eGFP in CeA was used to determine whether leakage of virus injected into CeA was apparent in LA/B (data not shown).

For the CnF-input/CeA-output LA/B neuronal labeling and the expression time-course experiments (**Fig. 1b, Extended Data Fig. 1f,g,h and Extended Data Fig. 2**), rAAV1-hSyn-NCre, rAAV1-hSyn-Cre, or rAAV1-hSyn-CCre virus was bilaterally injected into CnF, rAAV5-CAG-cDIO-tdTomato, rAAV5-CBA-fDIO-mCherry, or rAAV5-CAG-cDIO-tdTomato virus was unilaterally injected into LA/B, and retroAAV2-CAG-CCre and rAAV5-CAG-eGFP (mixed in 1:1 ratio by volume); retroAAV2-CBA-cDIO-Flp, CAV2-CMV-cDIO-Flp and rAAV5-CAG-eGFP (mixed in 1:1:1 ratio by volume); or retroAAV2-CAG-NCre and rAAV5-CAG-eGFP (mixed in 1:1 ratio by volume) were bilaterally injected into CeA for splitCre-PINCER (NCre for anterograde and CCre for retrograde, **Fig. 1b and Extended Data Fig. 2**), Cre/Flp-PINCER (**Fig. 1b and Extended Data Fig. 2c,d**) or reversed splitCre-PINCER (CCre for anterograde and NCre fore retrograde, **Extended Data Fig. 1f,g,h**) approaches, respectively. Rats were singly housed for the duration of the time-course experiments and then sacrificed as described in the Histology section. eGFP expression in CeA was used to determine whether leakage of virus injected into CeA was apparent in LA/B (representative images in **Fig. 1b**).

For the negative control experiments in the initial PINCER characterization (**Extended Data Fig. 3**), the following combinations of viruses were injected into the target regions based on the purpose of each experiment. Viral cocktails were bilaterally injected into CnF or unilaterally injected into LA/B or bilaterally injected into CeA. For the anterograde labeling negative control experiment using the splitCre-PINCER approach (**Extended Data Fig. 3a**), rAAV1-hSyn-NCre injection into CnF was omitted but rAAV5-CAG-cDIO-tdTomato virus was unilaterally injected into LA/B and retroAAV2-CAG-CCre and rAAV5-CAG-eGFP viruses (mixed in 1:1 ratio by volume) were bilaterally injected into CeA. For the retrograde labeling negative control experiments using the splitCre-PINCER approach (**Extended Data Fig. 3b**), retroAAV2-CAG-CCre injection into CeA was omitted but rAAV1-hSyn-NCre and rAAV5-CAG-cDIO-tdTomato viruses were bilaterally and unilaterally injected into CnF and LA/B, respectively. For the anterograde labeling of Cre negative control experiment using the Cre/Flp-PINCER approach (**Extended Data Fig. 3c**), rAAV1-hSyn-Cre injection into CnF was omitted but rAAV5-CBA-fDIO-mCherry virus was unilaterally injected into LA/B and retroAAV2-CBA-cDIO-Flp, CAV2-CMV-cDIO-Flp and rAAV5-CAG-eGFP viruses (mixed in 1:1:1 ratio by volume) were bilaterally injected into CeA. For the retrograde labeling of Cre-dependent expression of Flp negative control experiments using the Cre/Flp-PINCER approach (**Extended Data Fig. 3d**), retroAAV2-cDIO-Flp and CAV2-CMV-cDIO-Flp injection into CeA was omitted but rAAV1-hSyn-Cre and rAAV5-CBA-fDIO-mCherry viruses were bilaterally and unilaterally injected into CnF and LA/B, respectively. For the anterograde labeling of CCre negative control experiments using the reversed splitCre-PINCER approach (**Extended Data Fig. 3e**), rAAV1-hSyn-CCre injection into CnF was omitted but rAAV5-CAG-cDIO-tdTomato virus was unilaterally injected into LA/B and retroAAV2-CAG-NCre and rAAV5-CAG-eGFP viruses (mixed in 1:1 ratio by volume) were bilaterally injected into CeA. For the retrograde labeling of NCre negative control experiments using the reversed splitCre-PINCER approach (**Extended Data Fig. 3f**), retroAAV2-CAG-NCre injection into CeA was omitted but rAAV1-hSyn-CCre and rAAV5-CAG-cDIO-tdTomato viruses were bilaterally and unilaterally injected into CnF and LA/B, respectively. Rats were singly housed for 4+ or 8+ weeks (to allow viral infections and expression) for the splitCre-PINCER and Cre/Flp-PINCER negative control experiments, respectively, and then sacrificed as described in the Histology section. eGFP expression in CeA was used to determine whether leakage of virus injected into CeA was apparent in LA/B (data not shown).

For the anatomical characterization of the CnF→LA/B→CeA circuit (**Extended Data Fig. 4**), the following combinations of viruses were injected into the target regions based on the purpose of each experiment. Each combination of viruses was bilaterally injected into CnF or unilaterally injected into LA/B or bilaterally injected into CeA. For the quantification of the percentage of CnF-input alone neurons which are also CnF-input/CeA-output neurons in LA/B (**Extended Data Fig. 4a**), rAAV1-hSyn-Cre virus was bilaterally injected into CnF; rAAV5-CBA-cDIO-eGFP and rAAV5-CBA-fDIO-mCherry viruses (mixed in 1:1 ratio by volume) were unilaterally injected into LA/B; and retroAAV2-CBA-cDIO-Flp, CAV2-CMV-cDIO-Flp and rAAV5-CAG-mtagBFP viruses (mixed in 1:1:1 ratio by volume) were bilaterally injected into CeA. The mtagBFP expression in CeA was used to check no leakage of CeA injection into LA/B (data not shown). For the quantification of the percentage of CeA-output alone neurons which are also CnF-input/CeA-output neurons in LA/B (**Extended Data Fig. 4b**), rAAV1-hSyn-Cre virus was bilaterally injected into CnF; rAAV5-CAG-fDIO-eYFP virus was unilaterally injected into LA/B; and retroAAV2-CAG-tdTomato, retroAAV2-CBA-cDIO-Flp, CAV2-CMV-cDIO-Flp, and rAAV5-CAG-mtagBFP viruses (mixed in 1:1:1:1 ratio by volume) were bilaterally injected into CeA. The mtagBFP expression in CeA was used to check no leakage of CeA injection into LA/B (data not shown). For the distribution analysis of the CnF-input/CeA-output neurons in the anterior-to-posterior LA/B (**Extended Data Fig. 4c,d,e**), rAAV1-hSyn-NCre virus was bilaterally injected into CnF, rAAV5-CAG-cDIO-tdTomato virus was unilaterally injected into LA/B, and retroAAV2-CAG-CCre and rAAV5-CAG-eGFP viruses (mixed in 1:1 ratio by volume) were bilaterally injected into CeA. Rats were singly housed for 4+ weeks to allow viral infections and expressions and then terminated as described in the following Histology section. The eGFP expression in CeA was used to check no leakage of CeA injection into LA/B (data not shown).

For the molecular characterization of the three input-output projection defined LA/B neurons (**Fig. 1c,d,e and Extended Data Fig. 5a,b,c**), the following combinations of viruses were injected into the target regions based on the purpose of each experiment. Each combination of viruses were bilaterally injected into CnF or unilaterally injected into LA/B or bilaterally injected into CeA. To label the CnF-input alone LA/B neurons, rAAV1-hSyn-Cre virus was bilaterally injected into CnF and rAAV5-CBA-cDIO-eGFP virus was unilaterally injected into LA/B. To label the CeA-output alone LA/B neurons, rAAV5-CBA-cDIO-eGFP virus was unilaterally injected into LA/B and retroAAV2-CAG-Cre, CAV2-CMV-Cre and rAAV5-CAG-tdTomato viruses (mixed in 1:1:1 ratio by volume) were bilaterally injected into CeA. The tdTomato expression in CeA was used to check no leakage of CeA injection into LA/B. To label the CnF-input/CeA-output LA/B neurons, rAAV1-hSyn-NCre virus was bilaterally injected into CnF, rAAV5-CBA-cDIO-eGFP virus was unilaterally injected into LA/B, and retroAAV2-CAG-CCre and rAAV5-CAG-tdTomato viruses (mixed in 1:1 ratio by volume) were bilaterally injected into CeA. Rats were singly housed for 4+ weeks to allow viral infections and expressions and then terminated as described in the following Histology section. The tdTomato expression in CeA was used to check no leakage of CeA injection into LA/B (data not shown).

For the extended PINCER approach to investigate CeA output cell-type identification of the input-output projection defined LA/B neurons (**Extended Data Fig. 5d,e,f**), rAAV1-hSyn-Flp virus was bilaterally injected into CnF; rAAV-CBA-fDIO-mCherry virus was unilaterally injected into LA/B; and rAAV9-CAG-cDIO-TVA and rAAV9-CAG-cDIO-H2B-HA-2A-G helper viruses (mixed in 1:1 ratio by volume) were bilaterally injected into CeA. 4+ weeks after the helper AAV injection, EnvA-ΔG-Rabies-eGFP (EnvA-CVS-N2cΔG-eGFP) was bilaterally injected into CeA (same coordinate but 0.5 uL injection volume). Rats were singly housed for additional <2 weeks after rabies virus injection to allow viral infections and expressions and then terminated as written in the following Histology section. Immunohistochemistry based HA-tag expression in CeA was used to check no leakage of CeA injection into LA/B (data not shown).

For all calcium imaging and optogenetic inhibition experiments, following injection coordinates and viral volumes were used: CnF (AP: −8.3 mm from Bregma, ML: ±4.0 mm from Bregma, DV: −5.5 mm from pia with 20° angle to lateral direction, 0.3 μL injection volume), LA/B (AP: −3.0 mm from Bregma, ML: +5.2 or ±5.2 mm from Bregma, DV: −7.4 mm from pia, 0.5 μL injection volume), and CeA (AP: −2.3 mm from Bregma, ML: ±2.15 mm from Bregma, DV: −7.8 mm from pia with 14° angle to medial direction, 0.3 μL injection volume).

For the calcium imaging experiments in the three input-output projection defined LA/B neurons, following combinations of viruses were injected into the target regions based on the purpose of each experiment. Each combination of viruses were bilaterally injected into CnF or unilaterally injected into LA/B or bilaterally injected into CeA. Optical fiber (400-μm N.A. 0.66 mono fiber-optic canula, Doric Lenses) was unilaterally placed in LA/B (AP: −3.0 mm from Bregma, ML: +5.2 mm from Bregma, DV: −5.9 mm from pia) following virus injections. To target the CnF-input alone LA/B neurons, rAAV1-hSyn-Cre virus was bilaterally injected into CnF and rAAV5-Syn-cDIO-GCaMP6s virus was unilaterally injected into LA/B. To target the CeA-output alone LA/B neurons, rAAV5-Syn-cDIO-GCaMP6s virus was unilaterally injected into LA/B and retroAAV2-CAG-Cre, CAV2-CMV-Cre and rAAV5-CAG-tdTomato viruses (mixed in 1:1:1 ratio by volume) were bilaterally injected into CeA. The tdTomato expression in CeA was used to check no leakage of CeA injection into LA/B (data not shown). To target the CnF-input/CeA-output LA/B neurons, rAAV1-hSyn-NCre virus was bilaterally injected into CnF, rAAV5-Syn-cDIO-GCaMP6s virus was unilaterally injected into LA/B, and retroAAV2-CAG-CCre and rAAV5-CAG-tdTomato viruses (mixed in 1:1 ratio by volume) were bilaterally injected into CeA. Rats were singly housed for 7+ weeks to allow viral infections and expressions and then processed for calcium imaging experiments. After imaging experiments were finished, rats were terminated as written in the following Histology section. The tdTomato expression in CeA was used to check no leakage of CeA injection into LA/B (data not shown).

For the optogenetic inhibition experiments in the three input-output projection defined LA/B neurons, following combinations of viruses were injected into the target regions based on the purpose of each experiment. Each combination of viruses were bilaterally injected into CnF, LA/B or CeA. Optical fibers (200-μm core, N.A. 0.37, Thorlabs) were bilaterally placed in LA/B (AP: −3.0 mm from Bregma, ML: ±5.2 mm from Bregma, DV: −6.1 mm from pia) following virus injections. To target the CnF-input alone LA/B neurons, rAAV1-hSyn-Cre virus was bilaterally injected into CnF and rAAV5-CAG-cDIO-ArchT-tdTomato (for experimental group) or rAAV5-CAG-cDIO-tdTomato (for control group) viruses were bilaterally injected into LA/B. To target the CeA-output alone LA/B neurons, rAAV5-CAG-cDIO-ArchT-tdTomato (for experimental group) or rAAV5-CAG-cDIO-tdTomato (for control group) viruses were bilaterally injected into LA/B and retroAAV2-CAG-Cre, CAV2-CMV-Cre and rAAV5-CAG-eGFP viruses (mixed in 1:1:1 ratio by volume) were bilaterally injected into CeA. To target the CnF-input/CeA-output LA/B neurons with the splitCre-PINCER approach, rAAV1-hSyn-NCre virus was bilaterally injected into CnF, rAAV5-CAG-cDIO-ArchT-tdTomato (for experimental group) or rAAV5-CAG-cDIO-tdTomato (for control group) viruses were bilaterally injected into LA/B, and retroAAV2-CAG-CCre and rAAV5-CAG-eGFP viruses (mixed in 1:1 ratio by volume) were bilaterally injected into CeA. Rats were singly housed for 4+ weeks to allow viral infections and expressions and then processed for optogenetic experiments. To target the CnF-input/CeA-output LA/B neurons with the Cre/Flp-PINCER approach, rAAV1-hSyn-Cre virus was bilaterally injected into CnF; rAAV5-CBA-fDIO-ArchT-mCherry (for experimental group) or rAAV5-CBA-fDIO-mCherry (for control group) viruses were bilaterally injected into LA/B; and retroAAV2-CBA-cDIO-Flp, CAV2-CMV-cDIO-Flp and rAAV5-CAG-eGFP viruses (mixed in 1:1:1 ratio by volume) were bilaterally injected into CeA. Rats were singly housed for 7+ weeks to allow viral infections and expressions and then processed for optogenetic experiments. After optogenetic experiments were finished, rats were terminated as written in the following Histology section. The eGFP expression in CeA was used to check no leakage of CeA injection into LA/B (data not shown).

### Histology

To evaluate virus expression and fiber locations, rats were overdosed with intraperitoneal injections of 2.0 mL of 25% chloral hydrate and then transcardially perfused with 50 mL of PBS followed by 50 mL of ice-cold 4% paraformaldehyde (PFA) in PBS. After extraction, brains were post-fixed in 4% PFA in PBS overnight at 4°C and then processed in 20% sucrose in PBS for >3 days at 4°C. The brains were then embedded in Tissue-Tek OCT compound (Sakura, 4583) at −80°C for rapid freezing and then sliced into 40-μm (for fluorescent *in situ* hybridization experiment) or 50-μm (for the other experiments) coronal sections using a cryostat.

### Fluorescence *in situ* Hybridization

Brain sections were fixed in ice-cold 4% PFA in PBS for 15 minutes at 4°C, then washed three times in PBS for 5 minutes each. They were subsequently transferred through increasing concentrations of 50%, 75%, and 100% EtOH, with each step lasting 5 minutes at room temperature. The sections were then immersed in 100% EtOH for an additional 5 minutes at room temperature before being washed in PBS. Next, the sections were incubated in a 10 μg/mL Proteinase K (Promega, V3021) solution for 10 minutes at room temperature, followed by three washes in PBS for 5 minutes each at room temperature.

Multiplexed RNA fluorescence *in situ* hybridization (FISH)^109^ was performed using the HCR^TM^ reagents including RNA probes produced by Molecular Instruments, some of which are customized based on our requests (**Table 1**). The FISH procedures follow a protocol from Molecular Instruments (https://www.molecularinstruments.com/hcr-rnafish-protocols).

**Table 1.** HCR probes used in this study.

| Staining Group | Gene Name | Organism | Accession | Initiator | # Probes Pairs | Fluorophore |
| --- | --- | --- | --- | --- | --- | --- |
| 1,2,3 | *d2eGFP* | Transgenic | N/A | B1 | 12 | AlexaFluor 488 |
| 1 | *vGluT1*<br>(*Slc17A7*) | Rat | NM_053859.3 | B2 | 30 | AlexaFluor 546 |
|  | *vGluT2*<br>(*Slc17A6*) | Rat | NM_053527.1 | B2 | 30 | AlexaFluor 546 |
|  | *vGAT*<br>(*Slc32a1*) | Rat | NM_031782.2 | B3 | 30 | AlexaFluor 647 |
| 2 | *SST* | Rat | NM_012659.2 | B2 | 9 | AlexaFluor 546 |
|  | *PV* | Rat | NM_022499.2 | B3 | 9 | AlexaFluor 647 |
| 3 | *VIP* | Rat | NM_053991.1 | B2 | 20 | AlexaFluor 546 |

Brain sections were incubated in pre-warmed 37°C probe hybridization buffer (Molecular Instruments) for 10 minutes, then left overnight (>12 hours) at the same temperature with the addition of three pairs of probes at a concentration of 4 pmol/mL (**Table 1**). The hybridization solution was replaced with 100% probe wash buffer (Molecular Instruments), followed by three washes with decreasing concentrations of 75%, 50%, and 25% probe wash buffer diluted in increasing concentrations of 25%, 50%, and 75% 5X SSCT (5X SSC, Nacalai Tesque, 32146-91; 0.1% Tween 20, Sigma, P9416), each for 15 minutes at 37°C. Sections were then washed with 100% 5X SSCT for 15 minutes at 37°C and for 5 minutes at room temperature. The 5X SSCT was briefly exchanged for amplification buffer (Molecular Instruments) for 30 minutes at room temperature. The even (h1) and odd (h2) hairpins (Molecular Instruments) for each of the three genes were pooled, snap-cooled by heating to 95°C for 90 seconds, and then cooled to room temperature for 30 minutes in the dark. These hairpins were added to the amplification buffer at a final concentration of 60 pmol/mL, and the sections were incubated in the hairpin amplification solution overnight (>12 hours) at room temperature in the dark. This was followed by a brief wash with 5X SSCT, then two 30-minute incubations and a final 5-minute incubation at room temperature in 5X SSC. After washing, the sections were mounted on slides and cover-slipped with ProLong^TM^ Gold antifade reagent with DAPI (Thermo Fisher Scientific, P36935).

### Immunohistochemistry

Brain sections were blocked in 2% donkey serum in PBS containing 0.3% Triton-X (Nacalai Tesque, 35501-15; PBST) for 30 minutes and incubated in primary antibodies overnight at 4° in the dark. Sections were then rinsed in PBS and incubated in secondary antibodies diluted in PBST for 1 hour at room temperature in the dark. After rinsing with PBS, sections were mounted on slides and cover-slipped with ProLong^TM^ Gold antifade reagent (Thermo Fisher Scientific, REF P36934).

To amplify tdTomato and mCherry fluorescence, primary rabbit anti-RFP antibody (abcam, ab62341, 1:2000 dilution) was used with secondary donkey Alexa Fluor 555 anti-rabbit antibody (Thermo Fisher Scientific, A-31572, 1:1000 dilution). For enhanced expression of eGFP, eYFP and GCaMP6s, primary mouse anti-GFP antibody (Thermo Fisher Scientific, A-11120, 1:2000 dilution) was used with secondary donkey Alexa Fluor 488 anti-mouse antibody (Thermo Fisher Scientific, A-21202, 1:1000 dilution). For fluorescently visualizing helper AAV expression which were tagged with HA (**Extended Data Fig. 5d,e**), primary rabbit anti-HA (Cell Signaling Technology, #3724, 1:5000 dilution) was used with secondary donkey Alexa Fluor 555 anti-rabbit antibody (same as above). For fluorescently visualizing nuclei, Hoechst nucleic acid stain (Thermo Fisher Scientific, H21492, 1:1000 dilution) was used.

### Microscopy Imaging and Analysis

Brain sections processed with FISH and immunohistochemistry were imaged using fluorescence microcopy (BX63, EVIDENT) or confocal microscopy (FV3000, EVIDENT) with 4x magnification for local viral expressions to determine optical fiber placement, 10x magnification was used for cell identification and counting, 20x magnification was used for colocalization analysis of cellular fluorescence expression. Serial z-plane images for cell identification and counting and colocalization analysis of cellular fluorescence expression were taken across the range of tissue where fluorescence labeling was evident. Fluorescence microscopy images were exported using OlyVIA software (EVIDENT). Confocal microscopy images were analyzed for cell identification and counting and colocalization analysis of cellular fluorescence expression then exported using FV31S-SW software (EVIDENT). Exported images were then processed to adjust brightness/contrast using FIJI^110^ for presentation in Figures.

For determining optical fiber placement and viral expression in the calcium imaging experiments and the optogenetic inhibition experiments in the three anatomical connectivity defined LA/B subpopulations, LA/B tissue sections (−2.40 mm to −3.48 mm from Bregma) were analyzed. To check leakage of CeA viral injection into LA/B, section images including both CeA and LA/B (ranging from −1.92 mm to −3.00 mm from Bregma) were analyzed for viral expression of the fluorescent marker used in CeA injection experiments. Rats with no GCaMP or ArchT/tdTomato or ArchT/mCherry expressions in LA/B or leakage of CeA injection into LA/B were excluded from the experiments.

For cell identification in the CnF→LA/B→CeA circuit validation experiment (**Extended Data Fig. 1d**), the anterograde AAV-Cre and retrograde Cre-dependent fluorophore experiment (**Extended Data Fig. 1e**), the PINCER proof of principle experiments (CnF-input/CeA-output LA/B neuronal labeling) (**Fig. 1b, Extended Data Fig. 1f,g,h and Extended Data Fig. 2a,b**), and the PINCER negative control experiments (**Extended Data Fig. 3**), LA/B tissue sections (around −3.00 mm from Bregma) were analyzed for viral expression.

For cell counting in the proof of principle experiments (CnF-input/CeA-output LA/B neuronal expression time-course analysis) (**Extended Data Fig. 2c,d**), LA/B tissue sections (−2.40 mm to −3.36 mm from Bregma) were analyzed for counting tdTomato+ (splitCre-PINCER) or mCherry+ (Cre/Flp-PINCER) neurons. For cell counting to investigate the distribution of CnF-input/CeA-output neurons in the anterior-to-posterior LA/B (**Extended Data Fig. 4c,d,e**), all LA/B tissue sections (−1.56 mm to −4.08 mm) were analyzed by counting tdTomato+ neurons. For each animal, the number of tdTomato+ cells at each AP position were counted and divided by the total number of counted cells across all AP positions. The average of this percentage was then calculated across all animals to obtain “% tdTomato+ cells” (**Extended Data Fig. 4e**).

For colocalization analysis in the anatomical characterization of the CnF→LA/B→CeA circuit (**Extended Data Fig. 4a,b**), LA/B sections (−1.56 mm to −4.08 mm from Bregma) were analyzed to determine co-expression of eGFP/eYFP and mCherry/tdTomato. For the molecular characterization of the three input-output projection defined LA/B neurons (**Fig. 1c,d,e and Extended Data Fig. 5a,b,c**), LA/B sections (−1.56 mm to −4.08 mm from Bregma) were analyzed to determine co-expression of eGFP for connectivity-defined cell labels and labeled RNA probes.

For the extended PINCER approach used to investigate the overlap between *SST*+ CeA output neurons and the input-output projection defined LA/B neurons (**Extended Data Fig. 5d,e,f**), amygdala tissue sections including LA/B and CeA (−1.92 mm to −3.00 mm from Bregma) were analyzed for starter cells co-expressing rabies eGFP and immunohistochemistry-based HA labeling (for ΔG).

### Calcium Imaging Experiments

For all photometry experiments, rats were handled for two days and then habituated to being tethered to the patch cord (400-μm low-autofluorescence mono fiber-optic patch chords, Doric Lenses) for two days prior to recording in the pre-exposure/conditioning context. For aversive conditioning, rats were placed in a conditioning chamber (Med Associates Inc.) nested in a sound-attenuating cubicle with an audio speaker and electric foot-shock grid controlled by Med-PC IV (Med Associates Inc.). For appetitive conditioning, rats were placed in a conditioning chamber with reward port (Med Associates Inc.) delivering sucrose-water reward placed in a cup connected to an arm controlled by MED-PC IV (MED Associates Inc.) in a sound-attenuating cubicle with an audio speaker. Bulk calcium transient recording began 7+ weeks after viral GCaMP injections using a fiber photometry system (Doric Lenses) to measure 405-nm isosbestic and 470-nm GCaMP-dependent signals at 12 kHz sampling rate. Med-PC IV was used to simultaneously and independently control the photometry and video recording systems as well as stimulus presentation timing.

For the Pavlovian aversive conditioning experiments, on day 1 rats were first exposed to three auditory conditioned stimuli (CSs; 5.5 kHz pip, 1 Hz with 250ms on and 750-ms off, 20 seconds, 75 dB) during a pre-conditioning period without shock then conditioned with paired CS and unconditioned stimulus (US; electrical foot-shock, 0.7 mA, 1 second co-terminated with CS) three-times, in the conditioning context A. On day 2 rats were presented with three CSs in a separate retrieval context B (with different visual background and odor) to test aversive memory formation.

For the latent inhibition of Pavlovian aversive conditioning experiments, on day 1 and 2 rats were pre-exposed to 20 auditory CSs (CS pre-exposure, 5.5 kHz pip, 1 Hz with 250-ms on and 750-ms off, 20 seconds, 75 dB) each day (in total 40 auditory CSs for two days) in conditioning context A. On day 3 rats were first exposed to three auditory CSs (same as CS pre-exposure) then conditioned with three CS-US pairings in the conditioning context A. On day 4 rats were presented with three CSs in the retrieval context B to test aversive memory expression.

Behavioral freezing responses during CS presentation were manually scored by trained experimenters. Rats with significant contextual freezing, scored 4+ seconds in 20 seconds before the first CS presentation on each day, were excluded from experiments.

For the appetitive conditioning experiments, on day 1, food-deprived rats were first presented with three auditory CSs (5.5 kHz pip, 1 Hz with 250ms on and 750-ms off, 20 seconds, 75 dB) during a pre-conditioning period without sucrose water reward presentation. They were then conditioned with pairings of CS and unconditioned stimulus (US; 30% sucrose water reward delivery at a reward port in the conditioning chamber for 5 seconds at random timing within the last 10 seconds of the CS period) 24-times (appetitive conditioning session 1). Rats then rested in the home cage for 2+ hours, followed by another conditioning session with 24 CS-US pairings (appetitive conditioning session 2). On day 2, rats again received the same training protocol as on day 1 (appetitive conditioning session 3 and session 4). All these behavioral task procedures occurred in the same context.

Reward-seeking behavior was quantified as nose-poke behaviors into a reward port. These nose-poke events were detected through the breaking of a photo-beam located adjacent to the reward port. For each session, the probability of nose-poking in a given time block (e.g. CS period) was calculated as the number of nose-pokes across all trials during that time period divided by the number of trials. To quantify reward learning, nose-poke probabilities in session 1 and session 4 were compared.

### Data Analysis for Calcium Imaging

All photometry data were analyzed using custom MATLAB scripts (MATLAB R2021b, MathWorks). Raw calcium and isosbestic data were first down-sampled using a sampling rate of 120 Hz. Based on the timestamps of events (trial start, CS onset, etc.) during the experiments, the down-sampled data were extracted for each trial from 20 seconds before CS onset until 20 seconds after CS offset. The down-sampled isosbestic data for each trial were smoothed by a Gaussian kernel with 100 data-point width (0.8333 seconds). The smoothed isosbestic data were shifted vertically to maintain the same minimum as the corresponding down-sampled calcium data. ΔF/F was calculated using the down-sampled calcium data subtracted by the median of the shifted, smoothed isosbestic data and then divided by the median of the shifted, smoothed isosbestic data.

For CS- and US-evoked calcium peri-event time histograms in aversive conditioning and latent inhibition experiments, ΔF/F curves were normalized by subtracting the average values for 2 seconds before CS onset for all trials in a session and then subtracting this from each time bin across the pre-CS and CS periods for each trial. Subsequently the ΔF/F curves were smoothed by a Gaussian kernel with 200 data-point width (1.6667 seconds). All data were separated by experimental group. The area under curve (AUC) of ΔF/F was calculated for CS responses (0-10 seconds from CS onset) and US responses (0-11 seconds from US onset for aversive conditioning).

In appetitive conditioning experiments, ΔF/F peri-event time histograms were generated by normalized activity by calculating the average values for 2 seconds before CS onset for all trials in a session and then subtracting this from each time bin across the pre-CS and CS periods for each trial. We used a similar approach for reward (US) acquisition-evoked peri-event time histograms except that we used the 2 seconds before nose-poke onset during sucrose water reward (US) presentation as the normalization factor. Subsequently the ΔF/F curves were smoothed by a Gaussian kernel with 200 data-point width (1.6667 seconds). All data were separated by experimental group. The area under curve (AUC) of ΔF/F was calculated for CS responses (0-10 seconds from CS onset) and reward acquisition-evoked responses (0-2 seconds from nose-poke onset during reward presentation).

### Decoding Analyses

We trained a linear support vector machine (SVM) to predict distinct input-output projection-defined cell types in LA/B from fiber photometry ΔF/F traces, by using LIBSVM for MATLAB (http://www.csie.ntu.edu.tw/~cjlin/libsvm)^111^. We first down-sampled the ΔF/F signals from 120 Hz to 20 Hz, which reduced the sample size while retaining the temporal dynamics of neuronal activity. To extract task-relevant neuronal activity in the latent inhibition of aversive conditioning in each rat, we concatenated neuronal activity aligned with CS presentation during pre-exposure, pre-conditioning, CS-US paired presentation during conditioning, and retrieval for each rat. Each trial window spans 20 seconds before the onset of CS until 20 seconds after the offset of CS. For the CS pre-exposure, we included only the first three CSs on Day 1 and the last three CS on Day 2, in which we observed the most significant changes in ΔF/F signals (**Extended Data Fig. 9**). The ΔF/F trace in each trial was z-scored, using all data points in each trial, to make the signals comparable across rats. We subsequently concatenated the z-scored response vectors across rats. This process yielded a matrix of dimensions 18105 time-bins × 18 rats (6 rats’ data for each projection-defined cell types), with rows representing 6 rats with three distinct projection-defined cell type recordings (**Extended Data Fig. 11**). Neuronal responses at each time bin across the rats were then normalized to a range between 0 and 1. We performed one-vs-one decoding of different projection-defined cell types using 3-fold cross-validation test our hypotheses and control for model overfitting. Specifically, we trained the decoder iteratively on data randomly selected from 4 of 6 rats for each connectivity defined cell type and then tested the classification performance of the decoder on 2 ‘left-out’ rats’ data from each group. A model cost function (C) was optimized to a value giving minimum loss of validation by searching in a range between 10^-3^ and 10^3^ using the training data^112^. To obtain decoder performance distributions, this decoding procedure with cross validation was repeated 200 times. The statistical significance of decoding performance was determined by estimating the *P* value based on the obtained decoder performance distributions and a 50 % chance level. Specifically, it was computed as (1 + *X*) / (*N* + 1), where *X* represents the number of decoding performances below the chance level, and *N* represents the number of iterations. Since we repeated the decoding performance measurement 200 times, a decoder with no below chance performance resulted in *P* < 0.005, and a distribution with *x*% overlap with chance level performance range resulted in *P* ≈ *x* /100^113^.

To investigate essential time periods in behavioral tasks for projection-defined cell type decoding, we analyzed weights obtained from the SVM decoders trained with data from the aversive conditioning and latent inhibition. We took the absolute value of the weights in each decoder and then normalized them to a range between 0 and 1. This normalization enabled us to evaluate the relative contributions of neuronal activity during various task epochs and learning stages in predicting distinct projection-defined cell types. In each learning stage (CS pre-exposures, CS pre-conditioning, aversive conditioning, and CS retrieval), CS and US epochs were defined as the 0-18s and 0-19s windows after the onset of the CS and US, respectively. Since the US was presented 19 seconds after the CS onset in aversive conditioning, we took windows of equivalent size as post CS epochs during CS pre-exposures, CS pre-conditioning and CS retrieval to specifically analyze the influence of post CS responses or US expectation. These were defined as the 19-38 seconds after CS onset. The remaining time periods with minimum confounding from trial events were defined as baseline epochs, including −20-0 seconds before and 38-40 seconds after CS onset.

### Optogenetic Inhibition Experiments

Rats were handled for two days and then habituated to being tethered to the patch cord (200-μm mono fiber-optic patch chords, Doric Lenses) for two days prior to recording. For aversive conditioning, rats were placed in an aversive conditioning chamber nested in a sound-attenuating chamber with an audio speaker and electric foot-shock grid controlled by Med-PC IV. For appetitive conditioning, rats were placed in a reward conditioning chamber with reward port (Med Associates Inc.) in a sound-attenuating cubicle with an audio speaker and a cup connected to an arm delivering sucrose-water reward controlled by MED-PC IV (MED Associates Inc.). Behavior experiments were started 4+ (Cre only dependent viral infection including the splitCre-PINCER) or 8+ weeks (Cre and Flp dependent viral infection: Cre/Flp-PINCER) after viral injections for ArchT/tdTomato or ArchT/mCherry expression. Stimulus presentation timing was controlled and synchronized by Med-PC to trigger video frame recordings for rat behavioral analysis.

For the Pavlovian aversive conditioning experiments, on day 1 rats were conditioned with CS (5.5 kHz pip, 1 Hz with 250-ms on and 750-ms, 20 seconds, 75 dB) and unconditioned stimulus (US; electrical foot-shock, 0.7 mA, 1 second co-terminated with CS) pairings three-times, in a conditioning context A. On day 2 rats were presented with three CSs in separate retrieval context B (with different background and odor in the chamber from the context A) to test aversive memory formation. For optogenetic inhibition, 589-nm orange laser (SDL-589-XXXT, Shanghai Dream Laser Technology Co., Ltd) was delivered through an implanted fiber optic during US presentation (laser onset 500-ms before US onset, laser offset 500-ms after US offset).

For the latent inhibition of Pavlovian aversive conditioning, on day 1 and 2, rats were pre-exposed to 20 auditory CSs (CS pre-exposure, 5.5 kHz pip, 1 Hz with 250-ms on and 750-ms, 20 seconds, 75 dB) each day (in total 40 auditory CSs for two days) in a conditioning context A. On day 3 rats were conditioned with three CS-US pairings in the conditioning context A. On day 4 rats were presented with three CSs in another retrieval context B to test aversive memory expression. For optogenetic inhibition, 589-nm orange laser (Shanghai Dream Laser Technology Co., Ltd) was delivered through the implanted fiber optic probe during pre-exposure CS presentations (laser onset 500-ms before CS onset, laser offset 500-ms after CS offset) on day 1 and 2.

Behavioral freezing responses during CS presentation were manually scored by trained experimenters blind to the experimental conditions. Rats with significant contextual freezing, scored 4+ seconds in 20 seconds before the first CS presentation on each day, were excluded from experiments.

For the appetitive conditioning experiments, on day 1 food-deprived rats were conditioned with pairing of the CS and unconditioned stimulus (US; 30% sucrose water reward presentation at a reward port in the conditioning chamber for 5 seconds at random timing within the last 10 seconds the CS period) 24-times (appetitive conditioning session 1). Rats then rested in their home cages for 2+ hours, followed by another conditioning session with another 24 CS and US 24 pairings (appetitive conditioning session 2). On day 2, animals underwent the same procedure as on Day 1 (appetitive conditioning session 3 and session 4). For optogenetic inhibition, 589-nm orange laser (Shanghai Dream Laser Technology Co., Ltd) was delivered through the implanted fiber optic probe during US presentations at the reward port (laser onset 500-ms before US presentation onset, laser offset 500-ms after US termination) in all four conditioning sessions. All behavioral tasks were conducted in the same context.

Reward-seeking behavior was quantified as nose-poke behaviors into a reward port. These nose-poke events were detected through the breaking of a photo-beam located adjacent to the reward port and the beam break event was sent to a data acquisition (DAQ) system (National Instruments) to record the times-stamps of nose-poke and align them with CS and US timestamps. As above, for each session, the probability of nose-poking in a given time block (e.g. CS period) was calculated as the number of nose-pokes across all trials during that time period divided by the number of trials. To quantify reward learning, nose-poke probabilities in session 1 and session 4 were compared.

### Statistical Analysis and Data Presentation

All data were tested for normality using Shapiro-Wilk normality test. When data were normally distributed, parametric statistics were used. If the data did not pass a normality test, all statistical tests were run non-parametrically. Error-bars for parametric tests represent the mean ± SEM. Box plots were presented for non-parametric tests with median, box limits for upper and lower quartiles, whiskers for two sided 95^th^ percentile range of the distribution, and grey dots for data points outside of the 95^th^ percentile range of the distribution (outliers). All data points including outliers were used in statistical tests. For parametric comparisons of infection duration time points or sequential trials, 2way repeated measures (RM) ANOVA was used. The significant main effects of 2way RM ANOVA were followed by post-hoc Sidak’s multiple comparisons tests to detect differences between infection duration time-course.

Non-parametric comparisons between paired and unpaired two groups were tested using Wilcoxon and Mann-Whitney tests, respectively. For non-parametric comparisons of paired and unpaired sequential trials, Friedman and Kruskal-Wallis tests were used, respectively. The significant main effects of Friedman and Kruskal-Wallis tests were followed by post-hoc Dunn’s and Tukey-Kramer’s multiple comparisons tests.

All these statistical analyses were conducted using GraphPad Prism 8 (GraphPad Software Inc.).

## Supplementary Material

**Extended Data Fig. 1.**
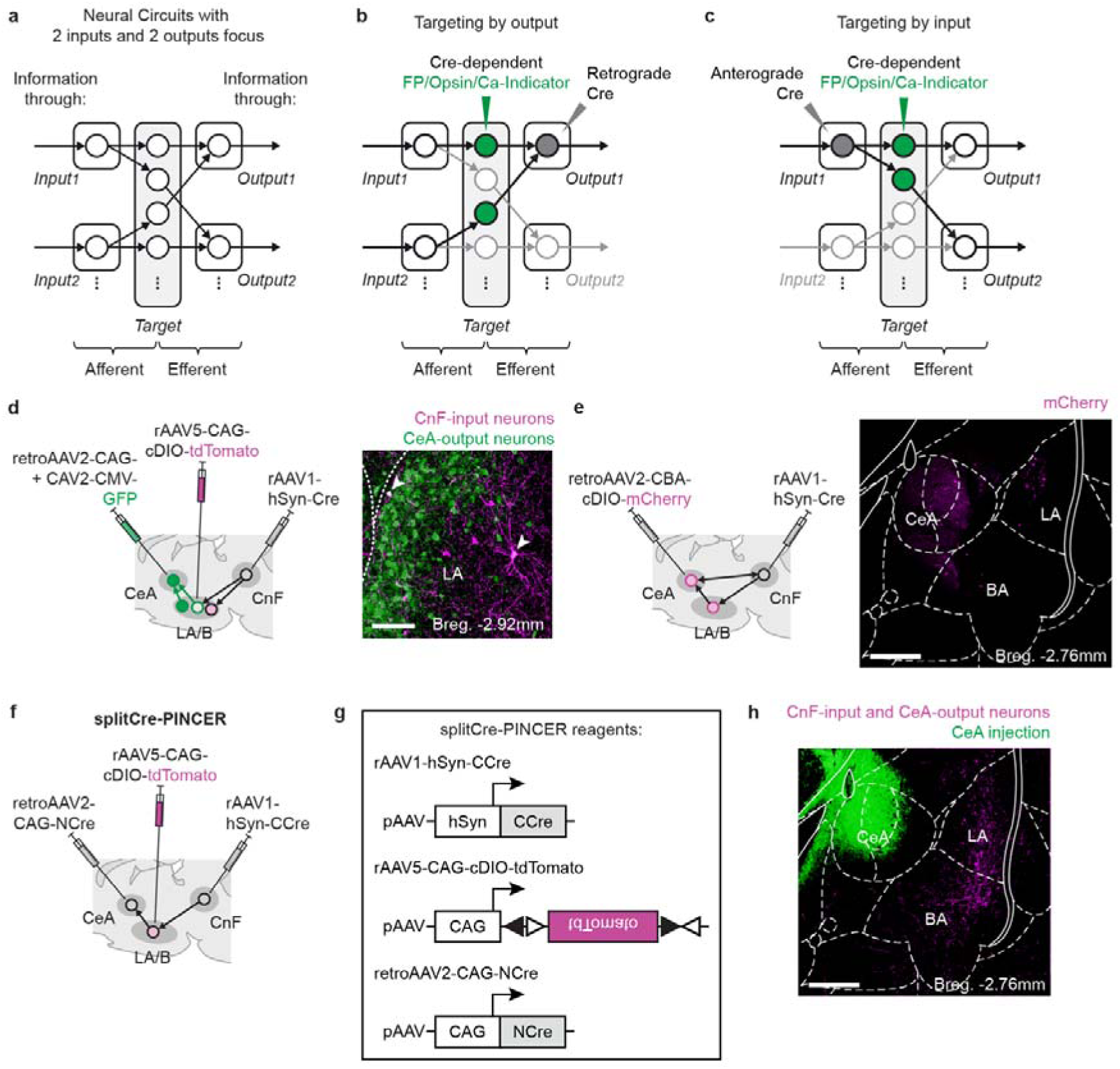
Identification of the CnF→LA/B→CeA input-output circuit and supporting data for a proof of principle of the PINCER method. **a-c,** Description of conventional anatomical connectivity-defined targeting of neuronal populations by anterograde transneuronal or retrograde viruses in a simplified input-output circuit. **a,** Schematic diagram of neural circuitry showing information flow through a target region receiving afferent inputs from two regions and sending efferent outputs to two target regions. **b,** Conventional output-based circuit-based cell type targeting with retrograde virus. The targeted cells project to a given region and receive input from multiple regions. **c,** Input-based circuit element targeting using transneuronal anterograde virus. These cells receive inputs from a specific region and project to multiple regions. **d,** Identification of the dual-transneuronal input-output circuit from the cuneiform nucleus (CnF) through the lateral and basal nuclei of the amygdala (LA/B) to the central nucleus of the amygdala (CeA). Schematic (left) of viral combinations to co-label LA/B neurons receiving input from CnF and sending output to CeA. Representative image of neurons (arrowheads) in LA co-expressing CnF-input (tdTomato) and CeA-output (eGFP) labels (replicated in N = 3 animals). **e,** Anterograde Cre and retrograde Cre-dependent fluorophore experiment. Schematic (left) of viral combinations and example image (right) showing labeling in LA/B and CeA. **f-h,** Reversing the injection site of the two splitCre-PINCER components (NCre and CCre) can capture the CnF-input/CeA-output neurons in LA/B. **f,** Schematic showing of reversed viral combinations. **g,** Reagents and pAAV plasmid sequence design used in the target site reversed splitCre-PINCER approaches. Open and filled triangles, incompatible *loxP* sites. **h,** Representative image of the CnF-input/CeA-output neurons (rAAV5-CAG-cDIO-tdTomato+) labeled by the reversed splitCre-PINCER approach (replicated in N = 3 animals). Scale bars, **d,** 100μm, **e,h,** 500μm. **e,h** Bregma distance in anterior-posterior direction.

**Extended Data Fig. 2.**
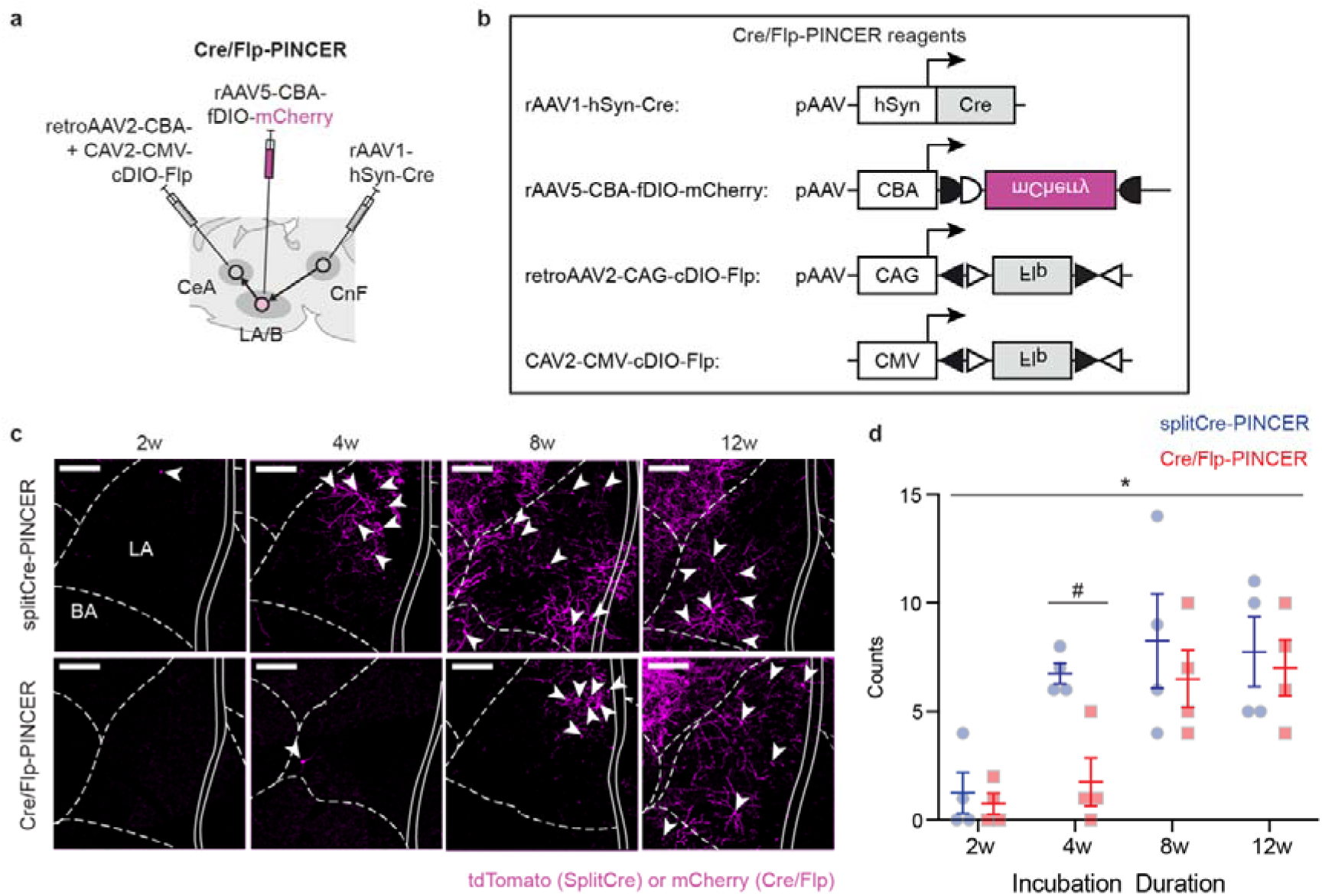
Comparison of the different patterns of PINCER methods. **a,** Schematic showing the Cre/Flip-PINCER approaches in the lateral and basal nuclei of the amygdala (LA/B) for labeling cells based on the inputs they receive from the cuneiform nucleus (CnF) and the outputs they send to the central nucleus of the amygdala (CeA). **b,** Reagents used and pAAV plasmid design for the Cre/Flp-PINCER approaches. Open and filled triangles, incompatible *loxP* sites; open and filled half circles, incompatible *FRT* sites. **c-d,** Viral expression time-course analysis shows more efficient labeling of the CnF-input/CeA-output neurons with the splitCre-PINCER approach compared with the Cre/Flp-PINCER approach. **c,** Representative images showing time-course of the CnF-input/CeA-output neuron labeling (denoted by arrowheads) with the splitCre- (top) and Cre/Flp-PINCER (bottom) approaches in LA/B. **d,** Quantification of cell labeling with the Cre/Flp- or splitCre-PINCER approaches with various infection durations (N = 4 animals per group). Four weeks after viral injection, the splitCre-PINCER (blue) approach labels more CnF-input/CeA-output neurons than the Cre\Flp-PINCER (red) approach but no difference at eight weeks and later (*P* = 0.0381, *F*_1,24_ = 4.815, 2way ANOVA, splitCre- vs Cre/Flp-PINCER in 2-12 weeks of infection duration; *P* = 0.0446, Sidak’s post-hoc multiple comparison test, splitCre- vs Cre/Flp-PINCER in four weeks infection duration). Scale bars, **c,** 200μm. **d,** Error bars: SEM. *: *P*<0.05, #: *P*<0.05.

**Extended Data Fig. 3.**
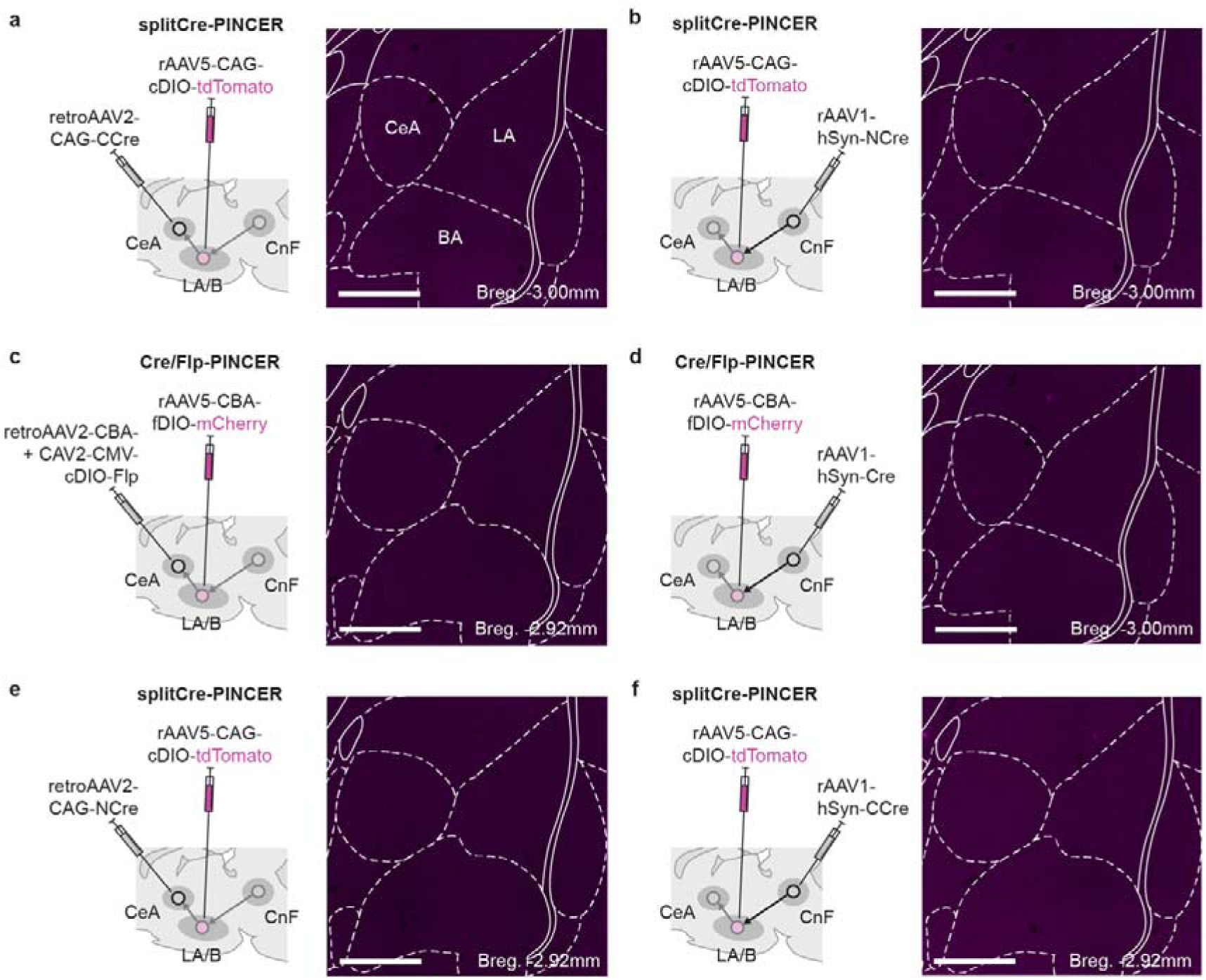
Negative control experiments for the splitCre-PINCER and Cre/Flp-PINCER approaches. **a,** Schematic (*left*) showing viral combinations for the splitCre-PINCER approach without rAAV1-hSyn-NCre injection into CnF. No labeling was found in LA/B (N = 3 animals; representative image, *right*). **b,** Schematic (*left*) showing viral combinations for the splitCre-PINCER approach without retroAAV2-CAG-CCre injection into CeA. No labeling was found in LA/B (N = 3 animals; representative image, *right*). **c,** Schematic (*left*) showing viral combinations for the Cre/Flp-PINCER approach without rAAV1-hSyn-Cre injection into CnF. No labeling was found in LA/B (N = 3 animals; representative image, *right*). **d,** Schematic (*left*) of viral combinations for the Cre/Flp-PINCER approach without retroAAV2-CBA-Cre and CAV2-CMV-Cre injection into CeA. No labeling was found in LA/B (N = 3 animals; representative image, *right*). **e,** Schematic (*left*) showing viral combinations for the reversed splitCre-PINCER approach without rAAV1-hSyn-CCre injection into CnF. No labeling was found in LA/B (N = 3 animals; representative image, *right*). **f,** Schematic (*left*) showing viral combinations for the reversed splitCre-PINCER approach without retroAAV2-CAG-NCre injection into CeA. No labeling was found in LA/B (N = 3 animals; representative image, *right*). **a-f,** Scale bars, 500μm. **a-f,** Bregma distance in anterior-posterior direction.

**Extended Data Fig. 4.**
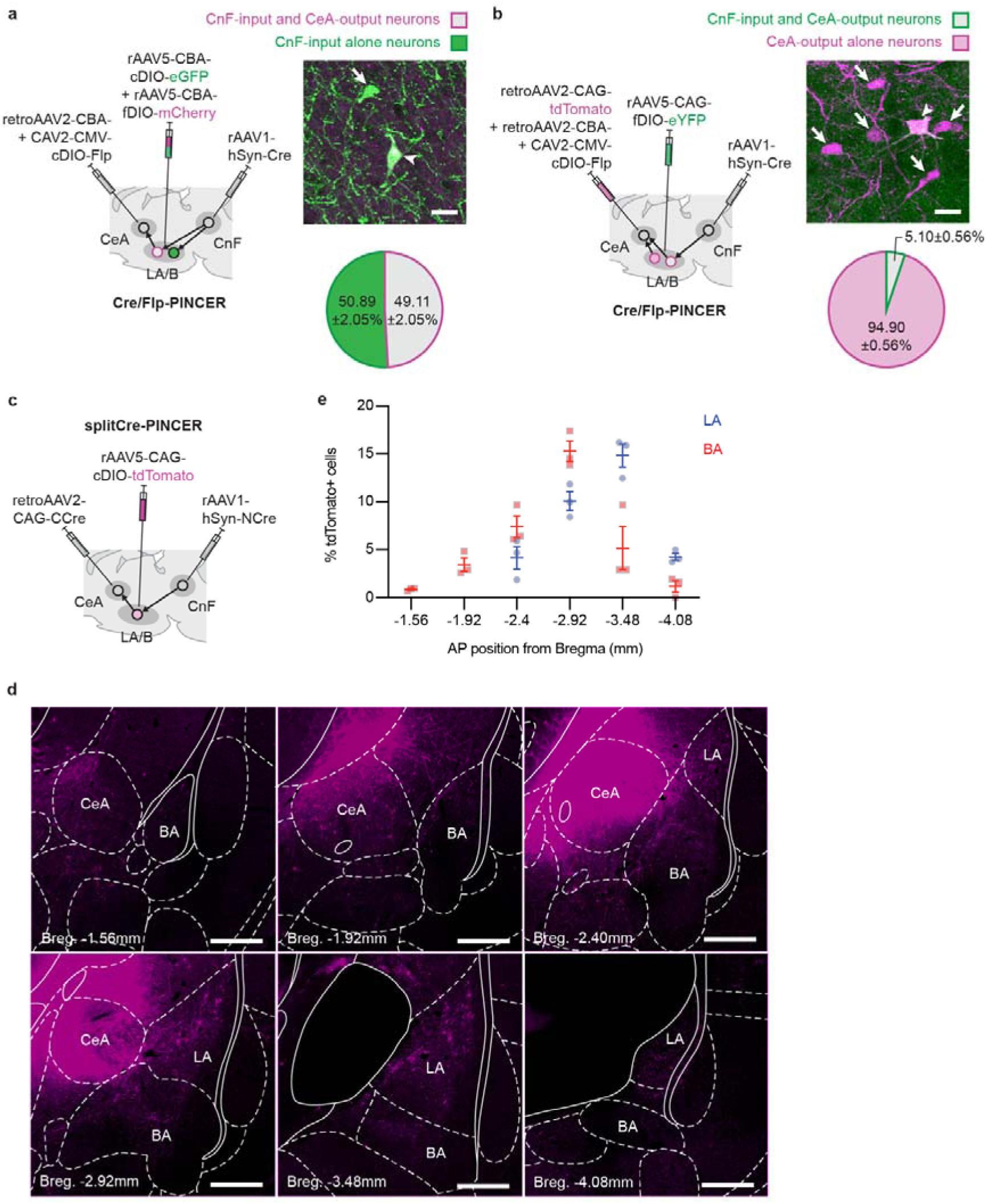
Anatomical characterization of anatomical connectivity defined neuronal populations using PINCER. **a,** Quantification of the percentage of CnF-input alone neurons which are also CnF-input/CeA-output neurons. Schematic (*left*) of Cre/Flp-PINCER viral combinations to label the input-output neurons with the input alone neurons. Representative image (*right top*) of a neuron expressing eGFP only (CnF-input alone neuron; denoted by arrow) and a neuron co-expressing eGFP and tdTomato (CnF-input/CeA-output neuron; denoted by arrowhead). Pie chart (*right bottom*) showing the percentage (mean±SEM) of CnF-input neurons which were also CnF-input/CeA-output labeled (N = 3 animals, total of n = 372 CnF-input neurons analyzed). **b,** Quantification of the percentage of CeA-output alone neurons which are also CnF-input/CeA-output neurons. Schematic (*left*) of Cre/Flp-PINCER viral combinations to label input-output neurons with output alone neurons. Representative image (*right top*) of a neuron expressing tdTomato only (CeA-output alone neurons; denoted by arrows) and a neuron co-expressing eYFP and tdTomato (CnF-input/CeA-output neuron; denoted by arrowhead). Pie chart (right *bottom*) showing the percentage (mean±SEM) of CeA-output neurons which were also CnF-input/CeA-output labeled (N = 3 animals, total of n = 2128 CeA-output neurons analyzed). **c-e,** Topographic distribution of the CnF-input/CeA-output neurons in anterior-to-posterior LA/B. **c,** Schematic of viral combinations used to label the CnF-input/CeA-output neurons in LA/B with the splitCre-PINCER method. **d,** Representative images showing labeled CnF-input/CeA-output neurons in LA/B throughout the anterior-to-posterior axis (N = 3 animals). **e,** Quantification of labeled CnF-input/CeA-output neurons reveals that their distribution throughout the anterior-to-posterior LA/B but more concentrated in the mid-to-posterior LA/B regions (N = 3 animals). Scale bars, **a-b,** 30μm, **d,** 500μm. **d,** Bregma distance in anterior-posterior direction. **e,** Error bars: SEM.

**Extended Data Fig. 5.**
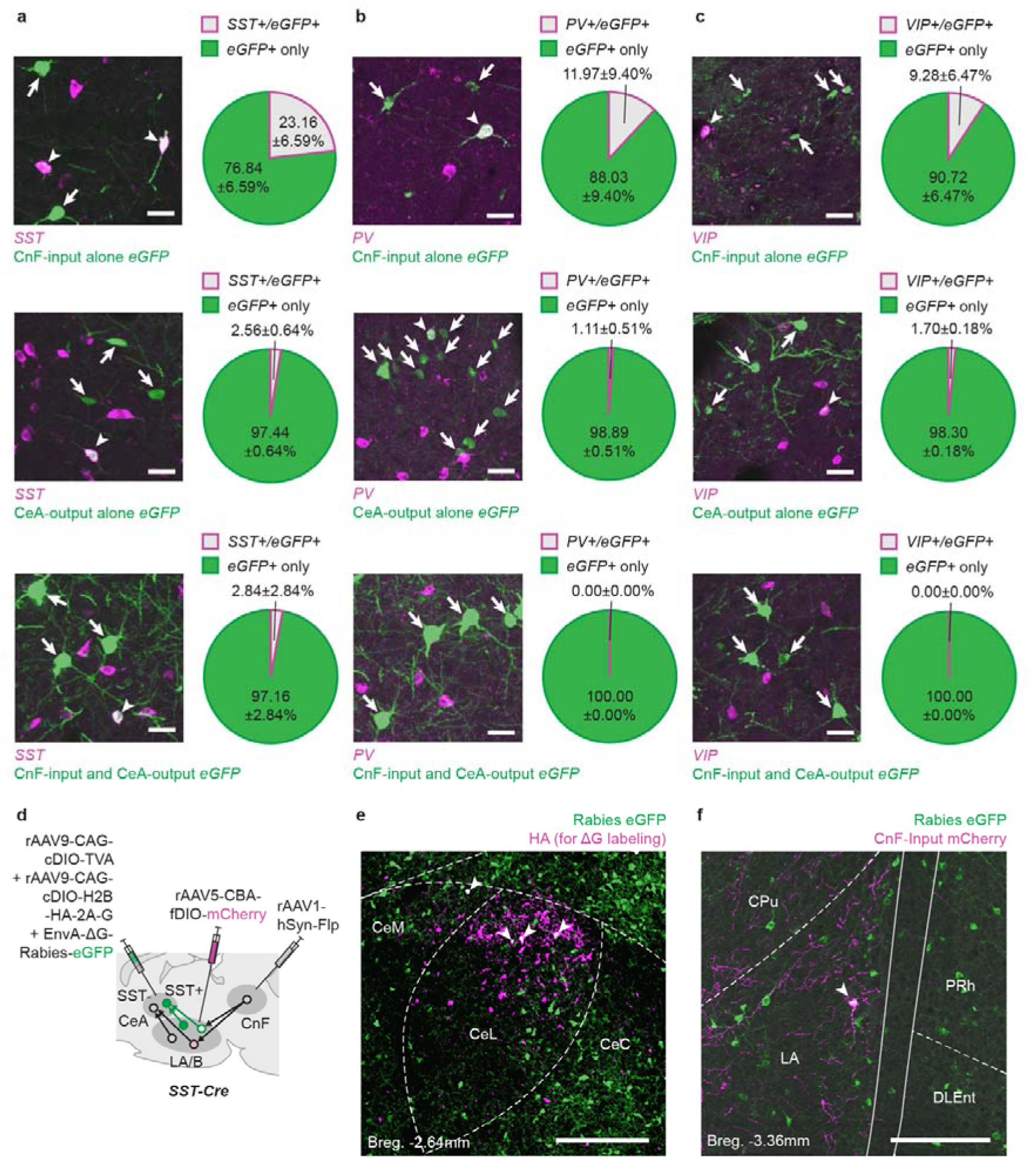
Further molecular/connectivity characterization of LA/B connectivity-defined neuronal populations. **a-c,** Co-expression analysis of molecular markers in the CnF-input alone, CnF-input/CeA-output, and CeA-output alone neurons for characterization of interneuron cell types. **a,** Representative images (*left*) of the CnF-input alone (*top*), CeA-output alone (*middle*), and CnF-input/CeA-output (*bottom*) labeled neurons and neurons expressing *somatostatin* (*SST*) in LA/B. Arrowheads and arrows denote some of the input-output projection-defined neurons co-expressing or not co-expressing *SST*. **a,** Pie charts (r*ight*) showing the percentage (mean±SEM) of the CnF-input alone neurons (*top*, N = 4 animals, total of n = 100 CnF-input neurons analyzed), CeA-output alone (*middle*, N = 3 animals, total of n = 840 CeA-output neurons analyzed), and CnF-input/CeA-output (*bottom*, averaged from N = 3 animals, total of n = 160 CnF-input/CeA-output neurons analyzed) neurons co-expressing *SST*. **b,** Representative images (*left*) of CnF-input only (*top*), CeA-output alone (*middle*), and CnF-input/CeA-output (*bottom*) labeled neurons and neurons expressing *parvalbumin* (*PV)* in LA/B. Arrowheads and arrows denote some of the input-output connectivity-defined neurons co-expressing or not co-expressing *PV*. Pie charts (*right*) showing the percentage of the CnF-input alone (*top*, N = 4 animals, total n = 100 CnF-input neurons analyzed), CeA-output alone (*middle*, N = 3 animals, total of n = 840 CeA-output neurons analyzed), and no (0.00%) CnF-input/CeA-output (*bottom*, N = 3 animals, total of n = 160 CnF-input/CeA-output neurons analyzed) neurons co-express *PV*. **c,** Representative images (*left*) showing the CnF-input alone (*top*), CeA-output alone (*middle*), and CnF-input/CeA-output (*bottom*) labeled neurons and neurons expressing *vasoactive intestinal peptide* (*VIP*) in LA/B. Arrowheads and arrows denote some of the input-output projection-defined neurons co-expressing or not co-expressing *VIP*. Pie charts (*right*) showing the percentage the CnF-input alone (top, N = 3 animals, total of n = 117 CnF-input neurons analyzed), CeA-output alone (*middle*, N = 3 animals, total of n = 664 CeA-output neurons analyzed), and no (0.00%) of CnF-input/CeA-output (*bottom*, N = 4 animals, total of n = 96 CnF-input/CeA-output neurons analyzed) neurons co-express *VIP*. **d-f,** An extension of the PINCER approach to investigate the molecular cell-type identification of target neurons which receive input from the input-output projection defined cells using PINCER in combination with trans-synaptic rabies virus labeling (N = 3 animals). **d,** Schematic showing viral combinations to label the CnF-input receiving LA/B neurons with mCherry and the SST+ CeA-projecting LA/B neurons with eGFP in SST-Cre rats. **e,** Representative image of starter cells (denoted by arrowheads) in CeA. Helper virus expresses immuno-stained HA which is linked to G glycoprotein. **f,** Representative image of a neuron (denoted by arrowhead) in the LA co-expressing eGFP (the SST+ CeA projecting neuron) and mCherry (the CnF-input receiving neuron). Scale bars, **a-c,** 30μm, **e-f,** 200μm. **e-f,** Bregma distance in Anterior-Posterior direction.

**Extended Data Fig. 6.**
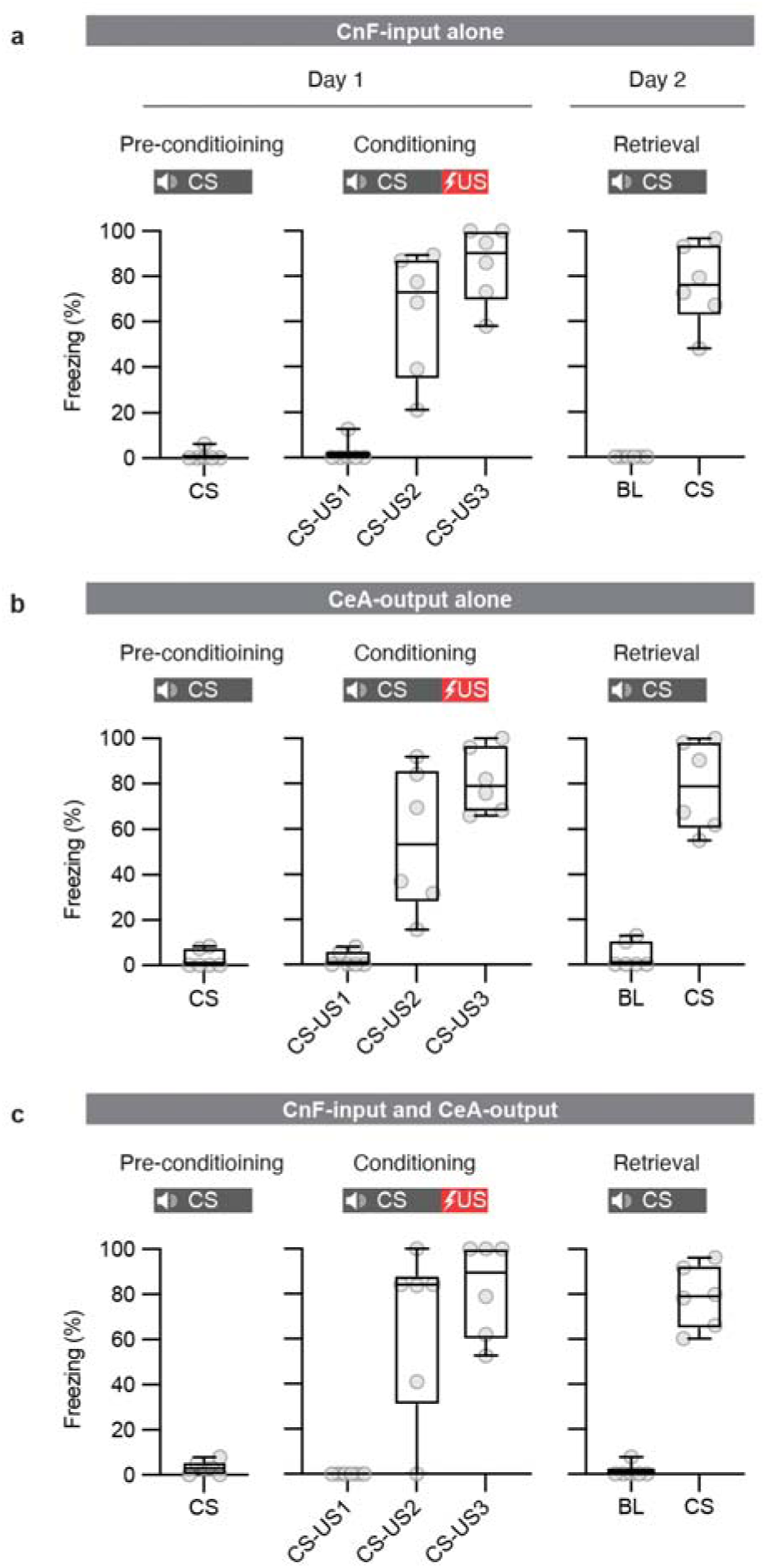
Freezing behavior during aversive conditioning for the fiber photometry experiment. **a-c,** Box plots showing behavioral freezing responses to auditory CSs increased during (middle) and after (right) aversive conditioning in the fiber photometry recording experiment. Behavioral data for photometry recordings in **a,** CnF-input-alone (N = 6 animals), **b,** CeA output-alone (N = 6 animals) and **c,** CnF-input/CeA-output (N = 6 animals) anatomical connectivity-defined LA/B neurons. **a-c,** In box plot: center line, median; box limits, upper and lower quartiles; whiskers, two-sided 95^th^ percentile range of distribution.

**Extended Data Fig. 7.**
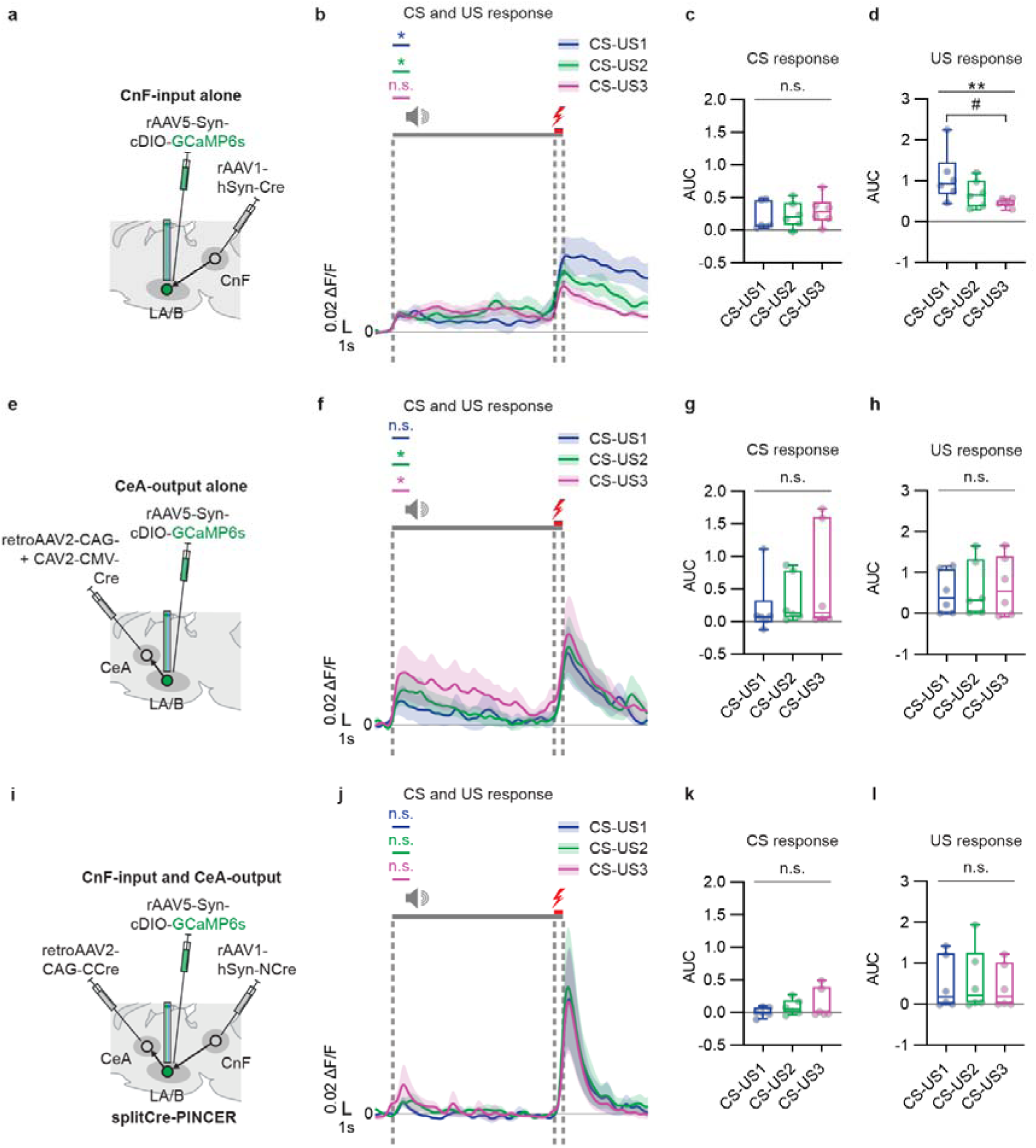
Population-specific calcium dynamics of LA/B anatomical connectivity-defined neurons during aversive conditioning. **a-d,** No change in auditory CS-evoked calcium response but decreasing US-evoked calcium responses in the CnF-input alone LA/B neurons over aversive conditioning trials. **a,** Schematic of viral combinations used to express GCaMP6s in the CnF-input alone neurons in LA/B for fiber photometry experiment. **b,** Peri-event time histogram (PETH) showing group averaged calcium responses to CS and US during aversive conditioning (N = 6 animals). CS calcium responses were significantly higher than baseline activity prior to CS onset except CS response in CS-US3 (2 seconds before CS-onset in vs. CS1 *P* = 0.0312, CS2 *P* = 0.0312, CS3 *P* = 0.0625, Wilcoxon test). **c,** Bar graphs show no significant differences for comparisons of CS-evoked ΔF/F area under the curve (AUC) responses of CS-US1 to CS-US3 of aversive conditioning. CS-US1 (blue), CS-US2 (green) and CS-US3 (magenta) (*P* = 0.7402, Friedman test, CS-US1 to CS-US3). **d,** Bar graphs show a significant decrease in US-evoked ΔF/F AUC responses from CS-US1 to CS-US3. CS-US1 (blue), CS-US2 (green) and CS-US3 (magenta) in aversive conditioning (*P* = 0.0055, Friedman test, CS-US1 to CS-US3; *P* = 0.0117, post-hoc multiple comparison test, CS-US1 vs. CS-US3). **e-h,** No change in auditory CS-evoked and US-evoked calcium response of the CeA-output alone neurons in LA/B during aversive conditioning. **e,** Schematic of viral combinations used to express GCaMP6s in the CeA-output alone LA/B neurons for fiber photometry experiment. **f,** PETH showing group averaged CS and US evoked calcium responses during aversive conditioning (N = 6 animals). CS calcium responses were significantly higher than baseline activity prior to CS onset except CS response in CS-US1 (CS1 *P* = 0.1562, CS2 *P* = 0.0312, CS3 *P* = 0.0312, Wilcoxon test by corresponding baselines). **g,** Bar graphs show no significant differences for the comparisons of CS-evoked ΔF/F AUC responses from CS-US1 to CS-US3 of aversive conditioning. CS-US1 (blue), CS-US2 (green) and CS-US3 (magenta) (*P* = 0.7402, Friedman test, CS-US1 to CS-US3). **h,** Bar graphs show no significant change in US-evoked ΔF/F AUC responses from CS-US1 to CS-US3. CS-US1 (blue), CS-US2 (green) and CS-US3 (magenta) in aversive conditioning (*P* = 0.4297, Friedman test, CS-US1 to CS-US3). **i-l,** No change in auditory CS-evoked and US-evoked calcium response of the CnF-input/CeA-output neurons in LA/B during aversive conditioning. **i,** Schematic of viral combinations used to express GCaMP6s in the CnF-input/CeA-output LA/B neurons for fiber photometry experiment. **j,** PETH showing group averaged CS and US evoked calcium responses during aversive conditioning (N = 6 animals). Calcium responses to CS were not significantly higher than baseline prior to CS onset (CS1 *P* = 0.0938, CS2 *P* = 0.0938, CS3 *P* = 0.1562, Wilcoxon test by corresponding baselines). **k,** Bar graphs show no significant difference for the comparison of CS-evoked ΔF/F AUC responses from CS-US1 to CS-US3 of aversive conditioning. CS-US1 (blue), CS-US2 (green) and CS-US3 (magenta) (*P* = 0.4297, Friedman test, CS-US1 to CS-US3). **l,** Bar graphs show no significant change in US-evoked ΔF/F AUC responses from CS-US1 to CS-US3. CS-US1 (blue), CS-US2 (green) and CS-US3 (magenta) in aversive conditioning (*P* = 0.4297, Friedman test, CS-US1 to CS-US3). **b,f,j,** Shades: SEM. **c,d,g,h,k,l** In box plot: center line, median; box limits, upper and lower quartiles; whiskers, two sided 95th percentile range of the distribution. n.s.: *P* > 0.05, *: *P* < 0.05, **: *P* < 0.01, #: *P* < 0.05.

**Extended Data Fig. 8.**
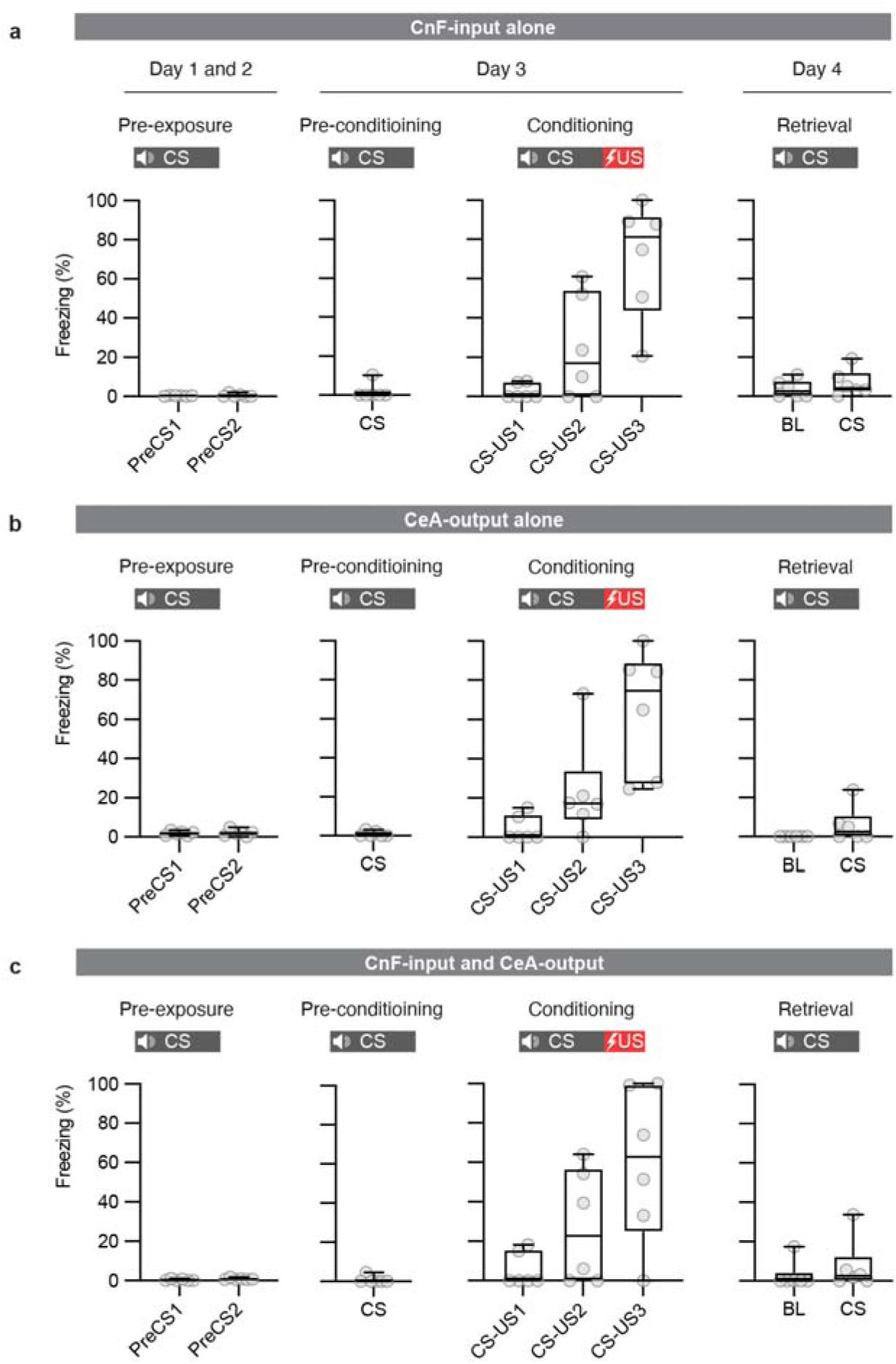
Freezing behavior during aversive conditioning of the latent inhibition with fiber photometry experiment. **a-c**, Box plots showing behavioral freezing responses to auditory CSs during pre-exposure, pre-conditioning, conditioning and retrieval in the fiber photometry recording experiment. Behavioral data for photometry recordings in **a,** CnF-input-alone (N = 6 animals), **b,** CeA output-alone (N = 6 animals) and **c,** CnF-input/CeA-output (N = 6 animals) anatomical connectivity-defined LA/B neurons. **a-c,** In box plot: center line, median; box limits, upper and lower quartiles; whiskers, two-sided 95^th^ percentile range of distribution.

**Extended Data Fig. 9.**
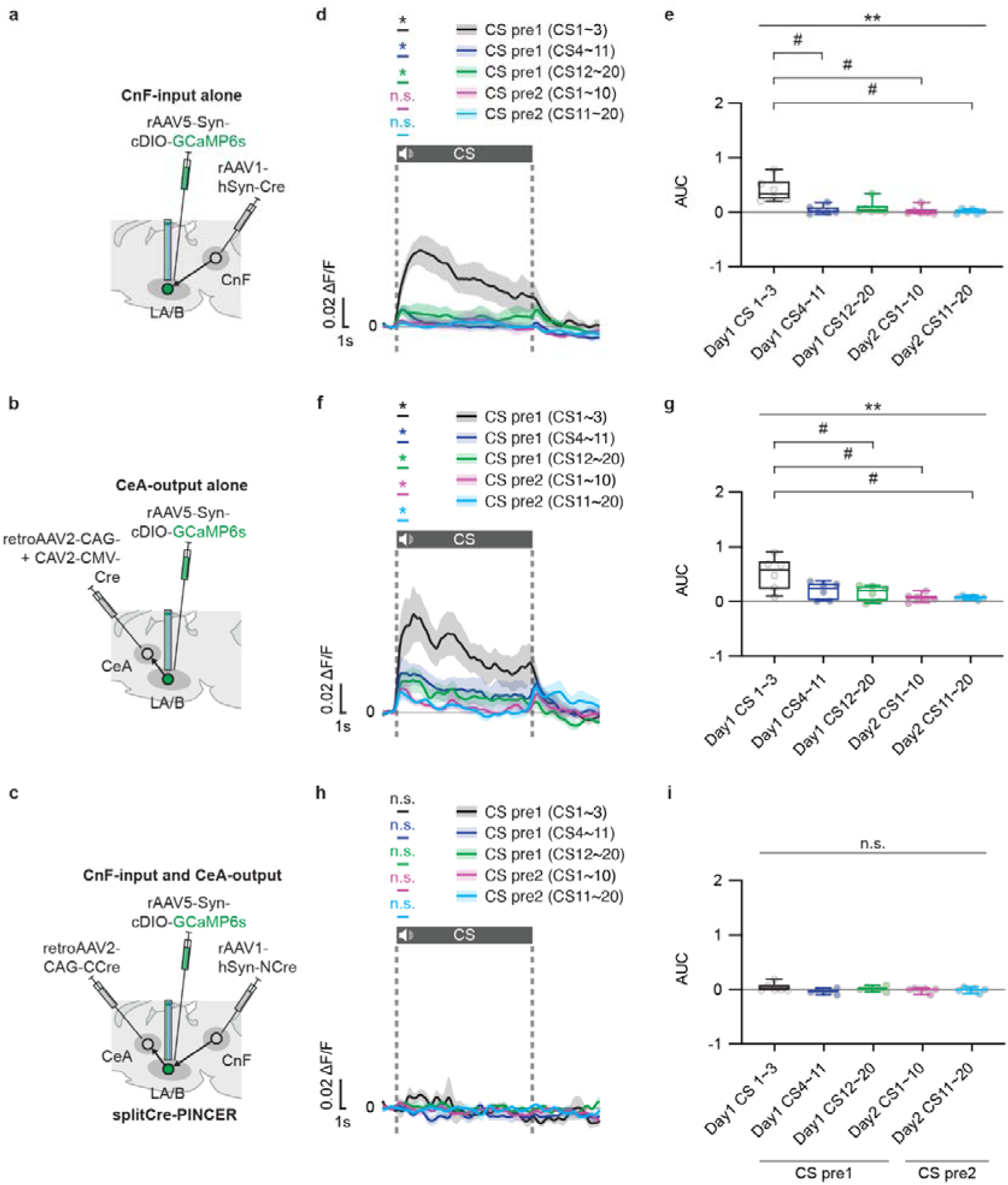
Population-specific calcium response dynamics of LA/B anatomical connectivity-defined neurons during CS pre-exposure of latent inhibition. **a-c,** schematics showing viral combinations used to express GCaMP6s in the **a,** CnF-input alone, **b,** CeA-output alone and **c,** CnF-input/CeA-output neurons in LA/B during latent inhibition. **d-e,** Significant reduction in auditory CS-evoked calcium response in the CnF-input alone neurons in LA/B during CS pre-exposure prior to aversive conditioning. **d,** Peri-event time histogram (PETH) showing a significant reduction in group averaged CS-evoked calcium responses to CS pre-exposure in the latent inhibition experiment (N = 6 animals). CS-evoked calcium responses were significantly higher than baseline activity prior to CS onset in CS pre-exposure trial blocks on day 1 (CS pre1) but not day 2 (CS pre2) (2 seconds before vs. after CS onset; CS pre1 CS1∼CS3 *P* = 0.0312, CS4∼CS11 *P* = 0.0312, CS12∼CS20 *P* = 0.0312; CS pre2 CS1∼CS10 *P* = 0.1562, CS11∼CS20 *P* = 0.5625; Wilcoxon test). **e,** Bar graphs show a significant decrease in CS-evoked ΔF/F area under the curve (AUC) responses for across trials during CS pre-exposure trial blocks. CS pre1 CS1∼CS3 (black), CS4∼CS11 (blue), and CS12∼CS20 (green); and CS pre2 CS1∼CS10 (magenta) and CS11∼CS20 (cyan) in CS pre-exposure (*P* = 0.0065, Friedman test, CS pre1 CS1∼CS3 to CS pre2 CS11∼CS20; *P* = 0.0191, *P* = 0.0349 and *P* = 0.0191, post-hoc multiple comparison test, CS pre1 CS1∼CS3 vs. CS pre1 CS4∼CS11, CS pre2 CS1∼CS10 and CS pre2 CS11∼CS20, respectively). **f-g,** Significant reduction in auditory CS-evoked calcium response in the CeA-output alone neurons in LA/B during CS pre-exposure prior to aversive conditioning. **f,** PETH showing a significant reduction in group averaged CS-evoked calcium responses to CS pre-exposure in the latent inhibition experiment (N = 6 animals). CS-evoked calcium responses were significantly higher than baseline activity prior to CS onset in CS pre-exposure trial blocks on day 1 and day 2 (CS pre1 CS1∼CS3 *P* = 0.0312, CS4∼CS11 *P* = 0.0312, CS12∼CS20 *P* = 0.0312; CS pre2 CS1∼CS10 *P* = 0.0312, CS11∼CS20 *P* = 0.0312; Wilcoxon test). **g,** Bar graphs show a significant decrease in CS-evoked ΔF/F AUC responses across trials in CS pre-exposure trial blocks. CS pre1 CS1∼CS3 (black), CS4∼CS11 (blue), and CS12∼CS20 (green); and CS pre2 CS1∼CS10 (magenta) and CS11∼CS20 (cyan) in CS pre-exposure(*P* = 0.0061, Friedman test, CS pre1 CS1∼CS3 to CS pre2 CS11∼CS20; *P* = 0.0349, *P* = 0.0349 and *P* = 0.0102, post-hoc multiple comparison test, CS pre1 CS1∼CS3 vs. CS pre1 CS12∼CS20, CS pre2 CS1∼CS10 and CS pre2 CS11∼CS20, respectively). **h-i,** No auditory CS-evoked calcium response of the CnF-input/CeA-output LA/B neurons during CS pre-exposure prior to aversive conditioning. **h,** PETH showing no significant CS-evoked calcium responses to CS pre-exposure in the latent inhibition experiment (N = 6 animals). CS-evoked calcium responses were not significantly higher than baseline activity prior to CS onset in CS pre-exposure trial blocks on day 1 and day 2 (CS pre1 CS1∼CS3 *P* > 0.9999, CS4∼CS11 *P* > 0.9999, CS12∼CS20 *P* = 0.0938; CS pre2 CS1∼CS10 *P* = 0.3125, CS11∼CS20 *P* = 0.4375; Wilcoxon test). **I,** Bar graphs show no change in CS-evoked ΔF/F AUC responses. CS pre1 CS1∼CS3 (black), CS4∼CS11 (blue), and CS12∼CS20 (green); and CS pre2 CS1∼CS10 (magenta) and CS11∼CS20 (cyan) (*P* = 0.3711, Friedman test, CS pre1 CS1∼CS3 to CS pre2 CS11∼CS20). **d,f,h,** Shades: SEM. **e,g,I,** In box plot: center line, median; box limits, upper and lower quartiles; whiskers, two sided 95th percentile range of the distribution. n.s.: *P* > 0.05, *: *P* < 0.05, **: *P* < 0.01, #: *P* < 0.05.

**Extended Data Fig. 10.**
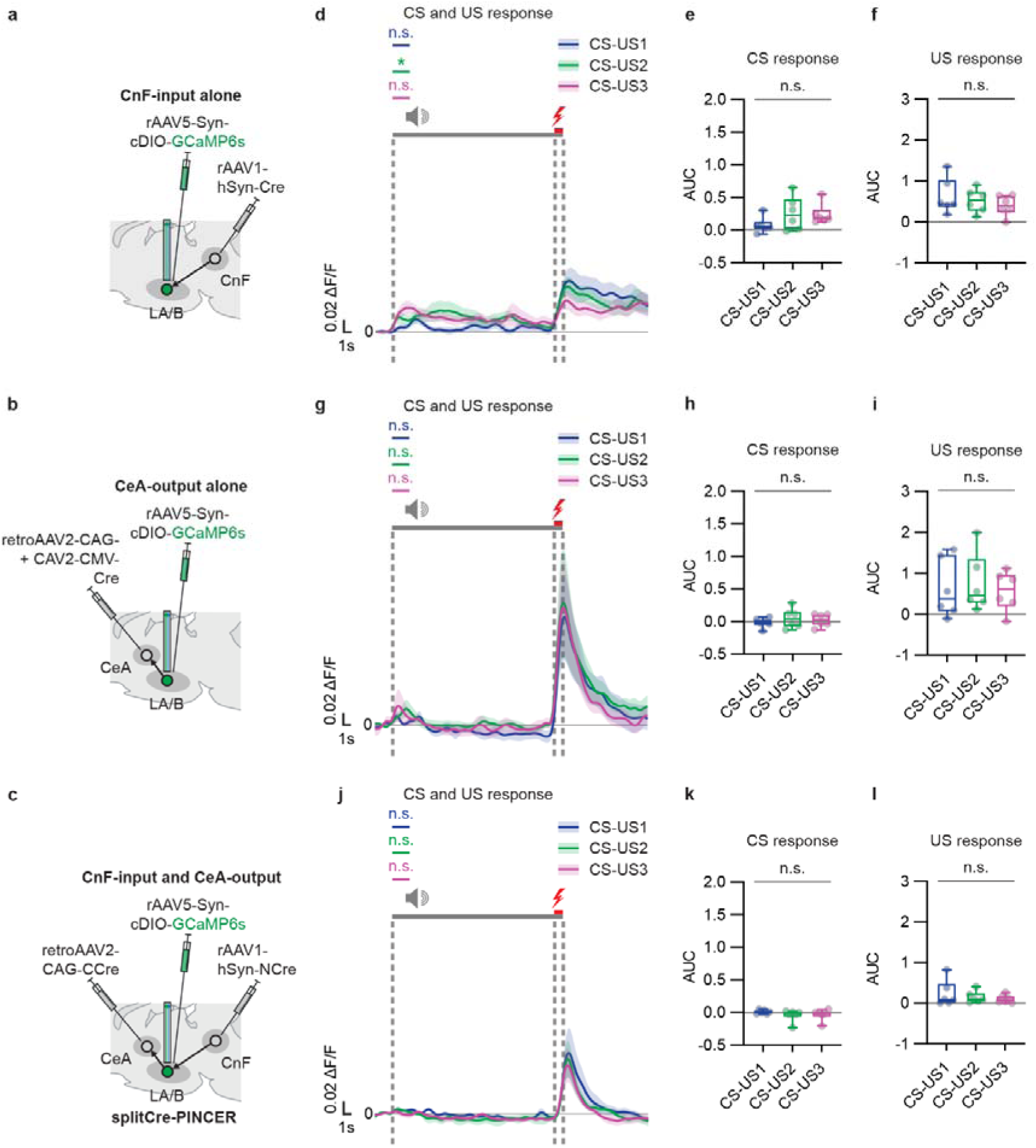
Population-specific calcium response dynamics of LA/B anatomical connectivity-defined neurons during latent inhibition. **a-c,** schematics showing viral combinations to express GCaMP6s in the CnF-input alone **a**, CeA-output alone **b,** and CnF-input/CeA-output **c,** neurons in LA/B during latent inhibition. **d-f,** No changes in auditory CS-evoked and US-evoked calcium response of the CnF-input alone neurons in LA/B during CS and US presentation across trials during aversive conditioning after latent inhibition. **d**, Peri-event time histogram (PETH) showing significant group averaged calcium responses to CS in CS-US2 during aversive conditioning (N = 6 animals). CS-evoked calcium responses were not significantly higher than baseline activity prior to CS onset in CS-US1 and CS-US3 (2 seconds after CS-onsets in CS1 *P* = 0.4375, CS2 *P* = 0.0312, CS3 *P* = 0.0625, Wilcoxon test). **e,** Bar graphs show no change in CS-evoked ΔF/F AUC responses across trials. CS-US1 (blue), CS-US2 (green) and CS-US3 (magenta) (*P* = 0.4297, Friedman test). **f,** Bar graphs show no change in CS-evoked ΔF/F AUC responses across trials. CS-US1 (blue), CS-US2 (green) and CS-US3 (magenta) (*P* = 0.4297, Friedman test, CS-US1 to CS-US3). **g-i**, No changes in auditory CS-evoked and US-evoked calcium response of the CeA-output alone neurons in LA/B to CS and US presentation during aversive conditioning following latent inhibition. **g,** PETH showing no significant group averaged calcium responses to CS during aversive conditioning (N = 6 animals). Calcium responses to CSs were not significantly higher than baseline activity prior to CS onset (CS1 *P* = 0.1562, CS2 *P* = 0.1562, CS3 *P* = 0.3125, Wilcoxon test by corresponding baselines). **h,** Bar graphs show no change in CS-evoked ΔF/F AUC responses across trials. CS-US1 (blue), CS-US2 (green) and CS-US3 (magenta) (*P* = 0.4297, Friedman test, CS-US1 to CS-US3). **i,** Bar graphs show no change in US-evoked ΔF/F AUC responses. CS-US1 (blue), CS-US2 (green) and CS-US3 (magenta) (*P* = 0.7402, Friedman test, CS-US1 to CS-US3). **j-l,** No changes in auditory CS-evoked and US-evoked calcium response of the CnF-input/CeA-output neurons in LA/B to CS and US presentation during aversive conditioning. **j,** PETH showing no significant group averaged calcium responses to CS during aversive conditioning (N = 6 animals). Calcium responses to CSs were not significantly higher than baseline activity prior to CS onset (CS1 *P* = 0.0938, CS2 *P* = 0.1562, CS3 *P* = 0.5625, Wilcoxon test by corresponding baselines). **k,** Bar graphs show no change in CS-evoked ΔF/F AUC responses across trials. CS-US1 (blue), CS-US2 (green) and CS-US3 (magenta) (*P* > 0.9999, Friedman test, CS-US1 to CS-US3). **l,** Bar graphs show no change in US-evoked ΔF/F AUC responses. CS-US1 (blue), CS-US2 (green) and CS-US3 (magenta) (*P* = 0.2522, Friedman test, CS-US1 to CS-US3). **d,g,j,** Shades: SEM. **e,f,h,I,k,l,** In box plot: center line, median; box limits, upper and lower quartiles; whiskers, two sided 95th percentile range of the distribution. n.s.: *P* > 0.05, *: *P* < 0.05.

**Extended Data Fig. 11.**
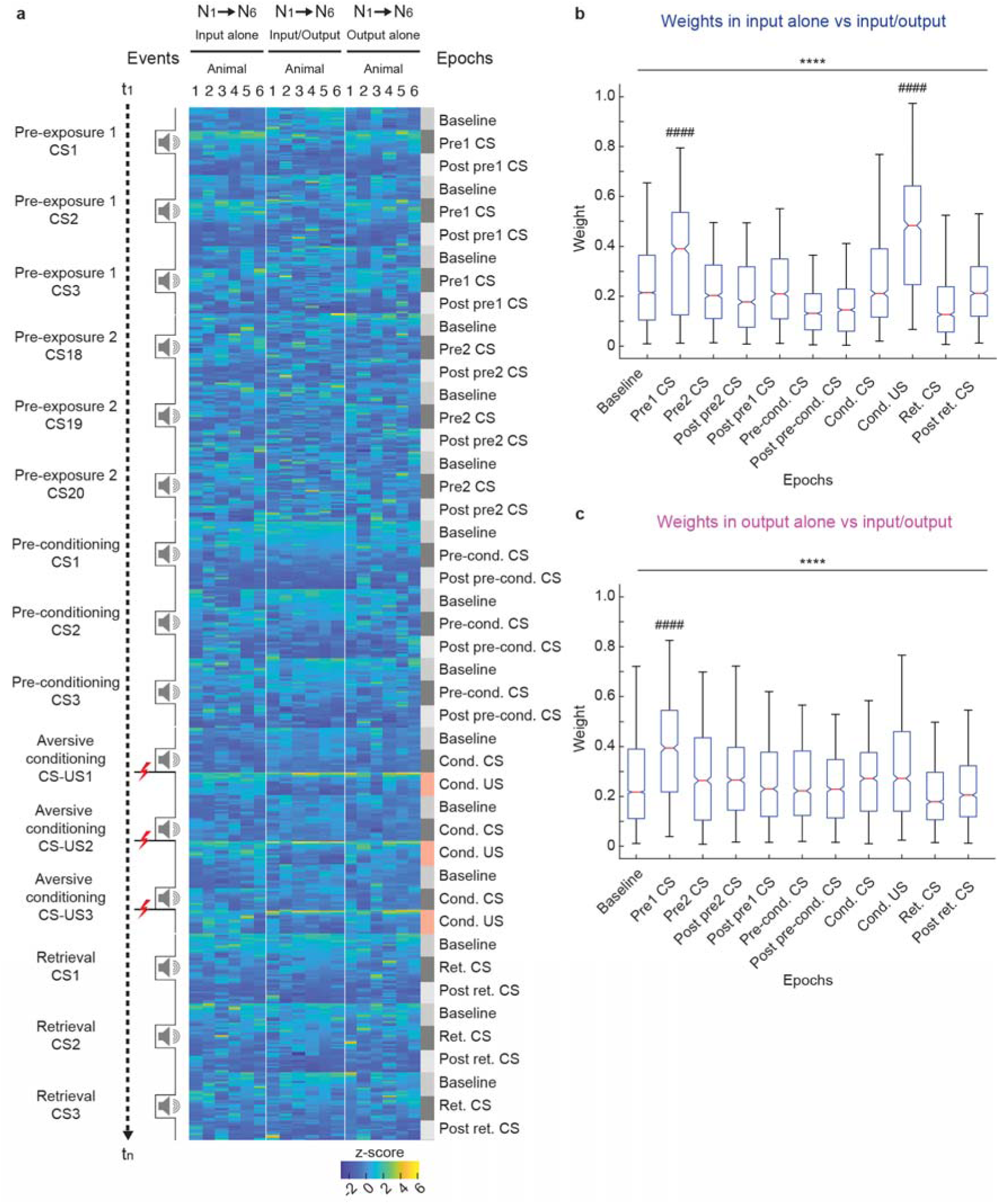
Decoding calcium response dynamics of anatomical connectivity defined neuronal populations from latent inhibition experiments. **a,** A heat map displays the z-scored data of ΔF/F GCaMP6s signals obtained during fiber photometry experiments from animals in all of the three anatomical connectivity defined populations (the input alone, input-output and output alone). x-axis represents individual animal numbers; y-axis shows calcium response for each trial (t) during individual training epochs. **b,** Box plots display decoding weights for classifying input-alone vs. input-output populations throughout epochs in the latent inhibition experiments. The pre1 CS epoch in CS pre-exposure and cond. US epoch in aversive conditioning (US cond.) have significantly higher decoding weights compared to all other epochs (*P* < 0.0001, Kruskal-Wallis test, all epochs; *P* < 0.0001 and *P* < 0.0001, post-hoc multiple comparison test, pre1 CS epoch vs. each of the other epochs and cond. US epoch vs. each of the other epochs, respectively). **c,** Box plots show decoding weights for classifying output alone vs. input-output populations throughout epochs in the latent inhibition experiment. The pre1 CS epoch in CS pre-exposure has significantly higher decoding weight compared to all the other epochs (*P* < 0.0001, Kruskal-Wallis test, all epochs; *P* < 0.0001, post-hoc multiple comparison test, pre1 CS epoch vs. each of the other epochs). **b-c,** In box plot: center red line, median; box limits, upper and lower quartiles; whiskers, two sided 95th percentile range of the distribution. ****: *P* < 0.0001, ####: *P* < 0.0001.

**Extended Data Fig. 12.**
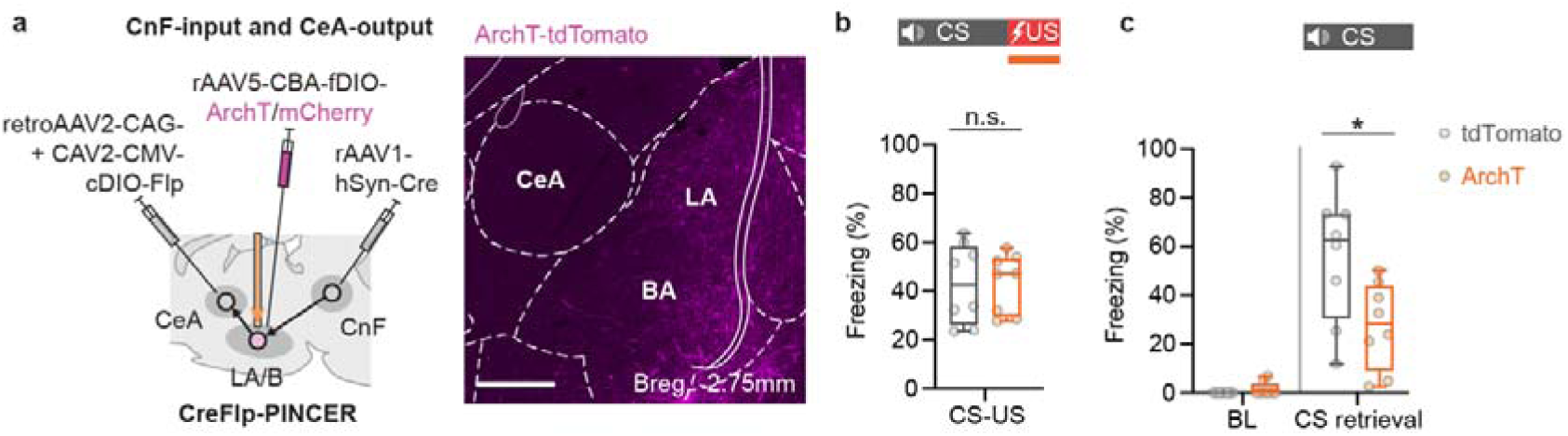
Photoinhibition of the CnF-input/CeA-output targeted neurons in LA/B with the Cre/Flp-PINCER approach. **a-c,** Inactivation of the CnF-input/CeA-output neurons targeted with the Cre/Flp-PINCER approach during shock presentation of aversive conditioning perturbs aversive associative memory formation. **a,** Schematic (*left*) of viral combinations used to express ArchT-tdTomato in the CnF-input/CeA-output neurons in LA/B with the Cre/Flp-PINCER method. Representative image (*right*) of ArchT-tdTomato expression in the CnF-input/CeA-output neurons in LA/B. Scale bar = 500μm and Breg. = anterior-posterior bregma distance. **b,** Bar graphs of CS-evoked behavioral freezing responses during the CS period of aversive conditioning show no significant difference between tdTomato only control (N = 8 animals) and experimental (N = 8 animals) groups (*P* = 0.9591, Mann-Whitney test). **c,** Bar graphs for behavioral freezing responses during retrieval show lower freezing responses in the ArchT group compared to the tdTomato only group (*P* = 0.0148, Mann-Whitney test). **b-c,** In box plot: center red line, median; box limits, upper and lower quartiles; whiskers, two sided 95th percentile range of the distribution. n.s.: *P* > 0.05, *: *P* < 0.05.

**Extended Data Fig. 13.**
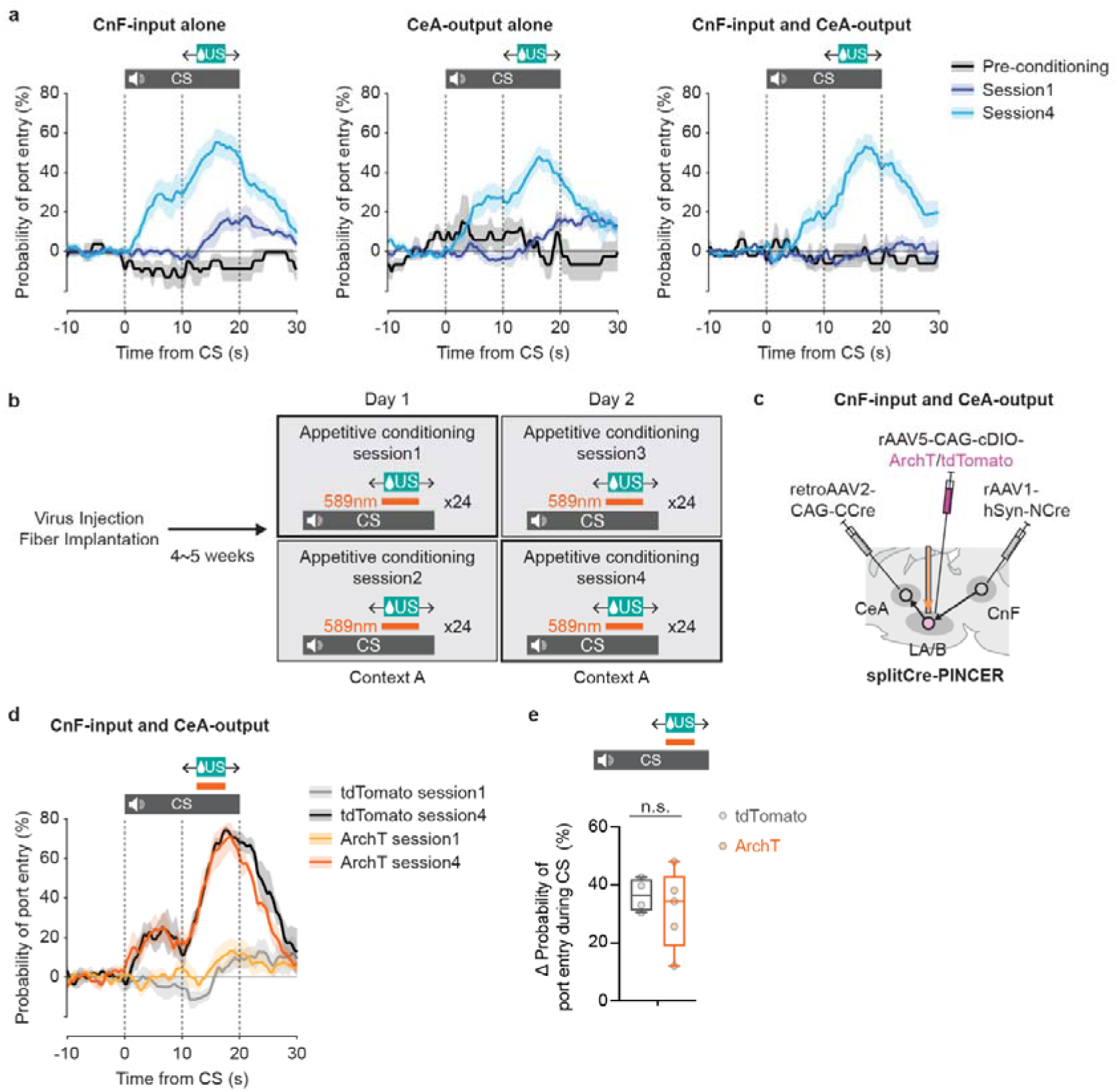
Reward-seeking behavior during appetitive conditioning with fiber photometry and optogenetics photoinhibition experiments. **a**, Peri-event time histogram (PETH) showing normalized probabilities of reward port entry (see methods) at CS and US presentation in pre-conditioning, session 1 and session 4 during appetitive conditioning for fiber photometry experiment (N = 6 animals for each of the 3 anatomical connectivity deveined group). Probabilities of reward port entry were normalized by subtracting the average probability of port entry for 10 seconds before CS onset for all trials in a session and then subtracting this from each time bin across the pre-CS, CS and post-CS periods for each trial. **b,** Experimental design of optogenetic manipulation in appetitive conditioning in which optogenetic inhibition of LA/B cell populations was applied during US presentations in reward port. **c-e,** Inactivation of the CnF-input/CeA-output neurons during US presentation in reward port does not affect nose poke behavior during appetitive associative learning. **c,** Schematic of viral combinations to express ArchT-tdTomato in the CnF-input/CeA output neurons in LA/B with splitCre-PINCER approach. **d**, PETH showing normalized probability of port entry at CS and US presentation during pre-conditioning, session 1 and session 4 during appetitive conditioning in tdTomato control and experimental groups (N = 4 and N = 5 animals for control and experimental groups, respectively). As mentioned in **a**, probabilities of reward port entry were normalized by subtracting the average probability of port entry for 10 seconds before CS onset for all trials in a session and then subtracting this from each time bin across the pre-CS, CS and post-CS periods for each trial. **e,** Bar graphs showing that optogenetic inhibition during US presentation in reward port does not affect the increase of the nose poke behavior calculated as the Δ probability of port entry to the port during CS from session 1 to session 4 between control and experimental groups (tdTomato N = 4 animals vs ArchT N = 5 animals, *P* = 0.7302, Mann-Whitney test). **a,d,** Shades: SEM. **e,** In box plot: center red line, median; box limits, upper and lower quartiles; whiskers, two sided 95th percentile range of the distribution. n.s.: *P* > 0.05.

**Extended Data Fig. 14.**
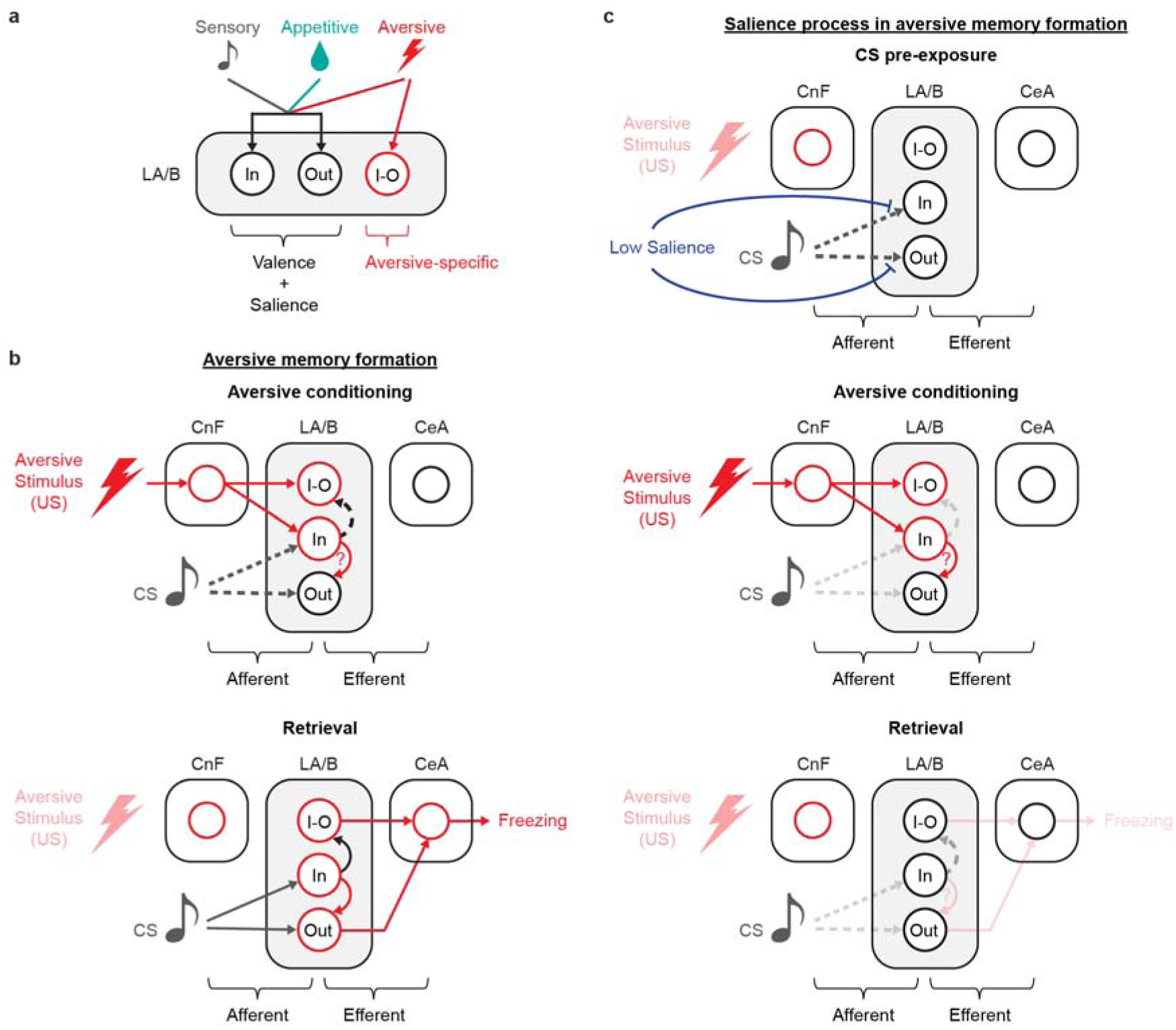
Hypothetical circuit models for the functional input-output organization underlying aversive and appetitive learning and salience processing in amygdala. **a,** A hypothetical circuit model for how valence and salience are integrated in anatomical connectivity-defined cell types in the LA/B. Whereas all of sensory, appetitive and aversive stimulus are integrated in the CnF-input and CeA-output alone neurons in the LA/B for valence and salience functions, only aversive stimuli is processed in the CnF-input and CeA-output neurons in the LA/B for serving an aversive function specifically. **b,** A hypothetical circuit model for aversive memory formation. During aversive conditioning (*middle*), auditory CS signals arrive at the CnF-input alone and CeA-output alone neurons in LA/B whereas aversive US signal is delivered directly to the CnF-input alone and CnF-input/CeA-output neurons, and to CeA-output alone neurons through the CnF-input alone neurons in LA/B (or other multimodal input pathways). The auditory CS and aversive US signals are associated in the CnF-input alone and CeA-output alone neurons in LA/B. During the consolidation process, local connections between the CnF input-alone cells and CnF-input/CeA-output neurons in LA/B associate auditory CS and aversive US signals. During retrieval (*bottom*), the CS can activate all three anatomical connectivity defined LA/B populations directly or indirectly (input-alone) trigger freezing through projections to CeA. **c,** A hypothetical circuit model for salience process during aversive memory formation. During preCS exposure (*top*), repeated auditory CS presentations produce depotentiation of auditory CS inputs to the CnF-input alone and CeA-output alone neurons in LA/B potentially through long-term depression (LTD) in the auditory area synaptic inputs onto these two projection-defined populations or long-term potentiation (LTP) of synaptic inputs from auditory thalamic/cortical regions onto inhibitory interneurons which are connected to these two connectivity-defined populations. During aversive conditioning (*middle*), the depotentiation mechanism occurring during preCS exposure desensitizes auditory CS signals delivered to the CnF-input alone and CeA-output alone neurons in LA/B. This prevents the initial association of auditory CS and aversive US signals in the CnF-input alone and CeA-output alone neurons as well as subsequent recruitment of the input-output defined cells, resulting in impaired behavioral learning (*bottom*).

**Extended Data Fig. 15.**
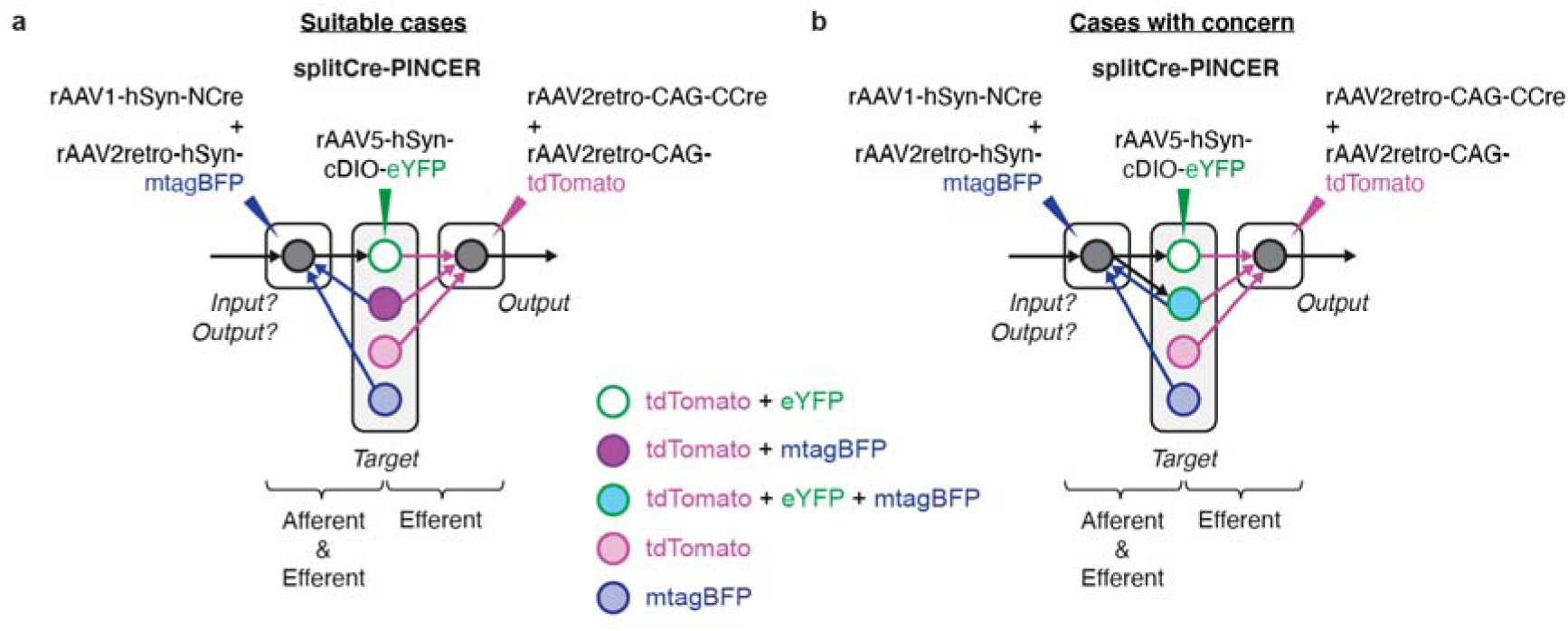
Considerations for applying the PINCER approach in reciprocally connected circuits. **a,** Cases in which PINCER can be used in reciprocally connected circuits in which cells receiving input from a given region do not themselves project back to that region. In these hypothetical cases, there is no overlap in the expression of splitCre-PINCER tagged input-output eYFP+ cells and retrogradely tagged mtagBFP+ neurons. **b,** Cases in which there is a potential issue using PINCER in reciprocally connected circuits. In these hypothetical cases, there would be overlapped expression of splitCre-PINCER tagged input-output eYFP+ cells and retrogradely tagged mtagBFP+ neurons due to reciprocal connectivity of some input-output neurons.

